# High-resolution DNA accessibility profiles increase the discovery and interpretability of genetic associations

**DOI:** 10.1101/070268

**Authors:** Aviv Madar, Diana Chang, Feng Gao, Aaron J. Sams, Yedael Y. Waldman, Deborah S. Cunninghame Graham, Timothy J. Vyse, Andrew G. Clark, Alon Keinan

## Abstract

Genetic risk for common autoimmune diseases is influenced by hundreds of small effect, mostly non-coding variants, enriched in regulatory regions active in adaptive-immune cell types. DNaseI hypersensitivity sites (DHSs) are a genomic mark for regulatory DNA. Here, we generated a single DHSs annotation from fifteen deeply sequenced DNase-seq experiments in adaptive-immune as well as non-immune cell types. Using this annotation we quantified accessibility across cell types in a matrix format amenable to statistical analysis, deduced the subset of DHSs unique to adaptive-immune cell types, and grouped DHSs by cell-type accessibility profiles. Measuring enrichment with cell-type-specific TF binding sites as well as proximal gene expression and function, we show that accessibility profiles grouped DHSs into coherent regulatory functions. Using the adaptive-immune-specific DHSs as input (0.37% of genome), we associated DHSs to six autoimmune diseases with GWAS data. Associated loci showed higher replication rates when compared to loci identified by GWAS or by considering all DHSs, allowing the additional discovery of 327 loci (FDR<0.005) below typical GWAS significance threshold, 52 of which are novel and replicating discoveries. Finally, we integrated DHS associations from six autoimmune diseases, using a network model (bird’-eye view) and a regulatory Manhattan plot schema (per locus). Taken together, we described and validated a strategy to leverage finely resolved regulatory priors, enhancing the discovery, interpretability, and resolution of genetic associations, and providing actionable insights for follow up work.

## Introduction

Most common autoimmune diseases affecting over 4% of the world’ population^1,2^ have a substantial polygenic heritable component^3^. Genome-wide association studies (GWAS) have been successful at linking hundreds of genomic loci to autoimmune diseases^4^, but understanding the molecular mechanisms influencing disease-risk remains challenging^5^. DNaseI hypersensitive site sequencing (DNase-Seq) is a high-throughput technology for genome-wide detection of DNaseI hypersensitive sites (DHSs) in a given cell type^6,7^, and DHSs are an excellent mark for regulatory DNA where transcription factors bind^8,9^. Recently, it has been shown that the majority of autoimmune disease risk alleles reside in DHSs active in adaptive immune cell types, such as T and B cells^10-13^. As recently suggested^11,14^, this presents the opportunity to focus association studies on regulatory DNA of such trait-relevant cell types. However, a large proportion of the regulatory DNA of a given cell type is shared with non-related cell types^8^. Since genes involved in adaptive-immune-specific functions (e.g. T cell receptor signaling^15^) are likely regulated by adaptive-immune-specific regulatory DNA, we suggest further focusing genotype-phenotype studies of autoimmune diseases, on regulatory DNA specific to adaptive immune cell types.

Regulatory marks other than DHSs such as certain histone marks are also available for many cell types^16,17^. However, when compared with DHSs, these other marks typically lack in two key factors we relied on, resolution, about 200-300 base pairs [bps] for a typical DHS compared to thousands of bps for a typical histone mark peak, and sequencing depth, over 200 million reads for each DNase-Seq data we used, compared to 10-30 million reads for a typical histone-mark ChIP-Seq experiment. For these two main reasons, here we used DHSs and not other regulatory marks.

This work focused on developing a framework for analyzing genetic data in light of the growing functional data. Perhaps the largest difference between our approach and many other reports is that we have used the functional data, instead of the genetic data, to dictate which regions of the genome would be evaluated (Fig. 1a,b). This resulted in, 1) a reduced requirement for multiple-testing correction and thereby increased statistical power; 2) tested units became context-specific, short regulatory regions thought to influence the studied trait, instead of all SNPs—the large majority of which have no functional impact; and 3) improved discovery and interpretation of genetic associations from current autoimmune-disease GWAS, afforded by trait-relevant, finely-resolved regulatory priors.

**Figure 1.**
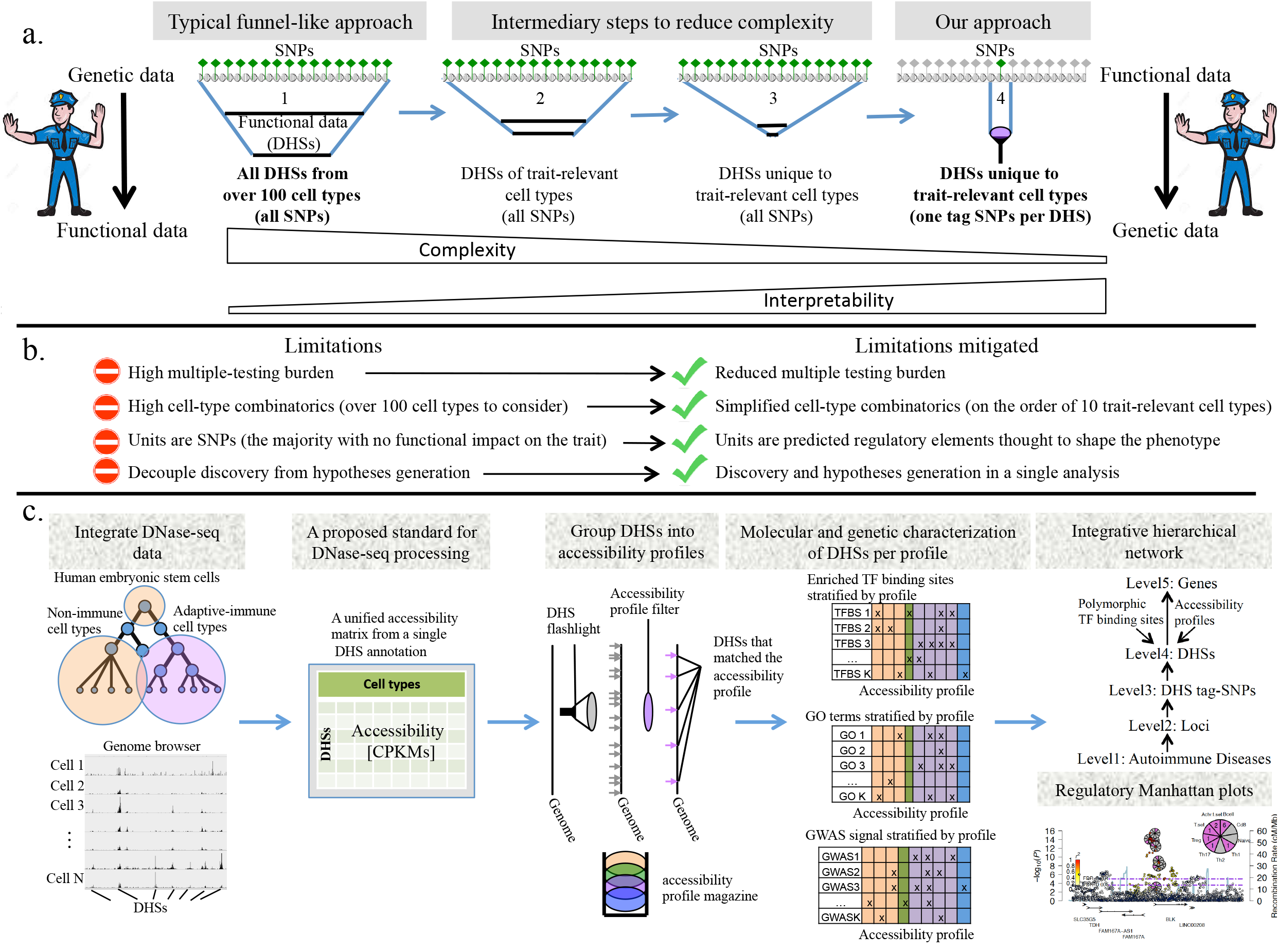
Overview. **a**. From left to right: (1) Functional analyses of GWAS data typically follow a funnel-like strategy starting with genetically associated loci and focusing in them on subsets of functional DNA (e.g. DHSs). That is, the genetic information dictates which DNA regions are considered. (2) A further funneling to smaller subsets of functional DNA may be desired by focusing on trait-relevant cell types. (3) We propose that an additional funneling to functional DNA unique to trait-relevant cell types could be even more advantageous. (4) By reversing the information flow from genetics → function, to function → genetics, and considering only one tag SNP per DHS, we transition from a funnel-like approach, to a flashlight-like approach where the functional information dictates which DNA regions are being considered. **b**. Several limitations of GWAS and functional, funnel-like approach to GWAS are mitigated by the proposed flashlight approach. **c.** DNase-seq data from adaptive-immune and non-immune cell types were integrated to generate a single DHS annotation and quantify DHS accessibility in cleavages per kilo base per million (CPKM) across cell types. Adaptive-immune-specific DHSs were identified and further resolved into accessibility profiles. DHSs belonging to each accessibility profile were characterized in terms of enrichment in binding motifs, GO terms, and GWAS signal in autoimmune diseases and unrelated control traits. Using the adaptive-immune-specific DHSs as input DNA, we associated DHSs to six autoimmune diseases with GWAS data. We integrated combined results from six autoimmune diseases using regulatory Manhattan plots (one per locus) and a hierarchical network model (over all loci).

Finally, since more results were generated than are possible to report in detail, we integrated results from six autoimmune diseases in a human-accessible format, in the hope of encouraging further investigations by others. This was achieved through a network model (integrating over all results) and a regulatory Manhattan plot schema (integrating results per locus). See Fig. 1c for an overview of our approach.

## Results

### Generating a unified DHS annotation from multiple samples and quantifying DHS accessibility

Currently, there is no unified genome-wide annotation for DHSs as there is for example for genes. Instead, each sample comes with its own unique DHS annotation. Moreover, there are currently no standards for quantifying accessibility as there are for quantifying gene expression, e.g. RPKM^18^. This makes working with DHSs from multiple DNase-seq experiments challenging. Here, we generated a single DHS annotation from fifteen available DNase-seq samples^16,17,19^: seven adaptive immune cell types (B, CD8+ T, Naïve CD4+ T, Th1, Th2, Th17, Treg), an innate immune cell (monocytes), and seven non-immune cell types (human embryonic stem cell, hereafter hESC, fetal brain, astrocyte, myoblast, fibroblast, epithelial, and hematopoietic stem cells, hereafter hematoSC). In total, we annotated 348,527 DHSs covering 3.4% of the human genome from DHSs that were significantly accessible in at least one of the above cell types (Methods). We then quantified accessibility per DHS and cell-type in number of cleavages per kilobase per million (CPKMs). We visualized the resulting accessibility matrix as a heatmap (Fig. S1). We saw that over half of all accessible DNA in adaptive-immune cell types is ubiquitously accessible across many other cell types, and that such ubiquitous DHSs tended to be more accessible when compared to adaptive-immune-specific DHSs. If one considered trait-relevant cell types alone, or considered cell types independently from one another, the ubiquitous DHSs might have overshadowed the less accessible but more interesting adaptive-immune-specific DHSs. This demonstrates the insights that can be gained by creating one DHS annotation from trait-relevant and trait-irrelevant cell types, and continuously quantifying accessibility in a single matrix.

### Grouping DHSs by accessibility profiles

Next, we hypothesized that grouping of DHSs by the subset of cell-types they were accessible in, would also group DHSs by distinct regulatory functions. Using the CPKM matrix, we scored DHSs for how well they matched cell-type-specific profiles, or profiles specific to predefined subsets of cell types (Figure 2a). Out of nineteen accessibility profiles we assayed, nine were unique to adaptive-immune cell types (purple), and were designed to group DHSs in broad strokes by salient immune functions: B cells specific DHSs (antibody production^20^), CD8+ T cell specific DHSs (cell-mediated killing^21^), Naïve CD4+ T cell specific DHSs (maintenance of non-activated T cells^22^), CD4+ Th1 specific DHSs (cytokine-induced cell-mediated immunity^22^), CD4+ Th2 specific DHSs (cytokine-induced antibody production^22^), CD4+ Th17 specific DHSs (cytokine-induced mucosal immunity^22^), CD4+ Treg specific DHSs (cytokine-induced anti-inflammatory response and tolerance to self antigens^22^); and for subsets of cell types: DHSs specific to all six T lineages (T cell maintenance), and DHSs specific to all four activated CD4+ T cells, namely: Th1, Th2, Th17 and Treg (CD4+ T cell activation). As examples, we present genome-browser views of two uniformly selected DHSs per accessibility profile (Fig. 2b, Fig. S2). The accessibility annotation, CPKM matrix, and match-scores per accessibility profile are available in Table S1.

**Figure 2.**
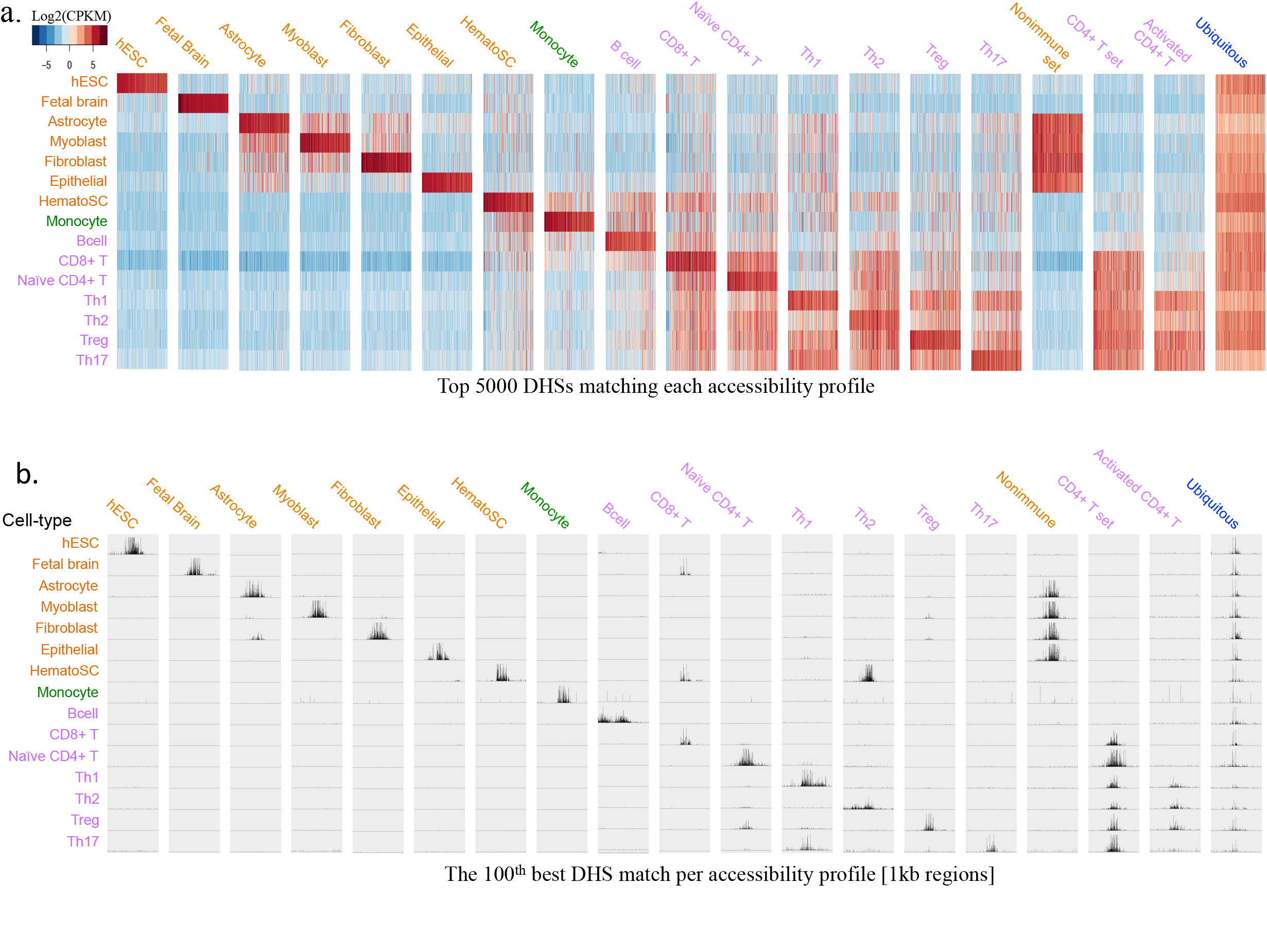
Grouping DHSs by accessibility profiles. **a.** Each DHS received a score for how well it matched one of nineteen predefined accessibility profiles. The top 5000 DHSs matching each accessibility profile (columns) were visualized as a heatmap across all fifteen cell-types (rows). We noted that lineage-specific DHSs for the related T cells were captured quantitatively as having higher accessibility in one T lineage compared to the others, but not in binary terms. **b.** An IGV genome browser view (Broad Institute), across all fifteen cell-types, of a 1kb DNA region surrounding the midpoint of DHSs uniformly selected as the 100^th^ best match per accessibility profile. Reads per millions were used as input to IGV. For each panel the track height was fixed at a single value across all tracks to allow comparisons across cell types.

We also compared the grouping of DHSs into accessibility profiles based on the CPKM matrix, with the alternative of grouping DHSs into the same profiles using a binary matrix in which DHSs were annotated as accessible (1) or not accessible (0)–the common practice for DHS data^8,11,16,19,23,24^. With respect to resolving accessibility profiles among the relate T lineages, the CPKM matrix increased the number of identified DHSs per profile (sensitivity; Fig. S3), proportion of correctly classified DHSs per profile (specificity; Fig. S3), and overall accessibility at identified DHSs per profile (signal-to-noise ratio; Fig. S4; Methods).

### Underlying DHSs of different accessibility profiles are distinct combinations of DNA sequence motifs

The expected molecular basis for DNA accessibility differences between cell types is a corresponding difference in TF occupancy^9,25^. Therefore, to assess whether grouping of DHSs into accessibility profiles was cell type dependent and not strongly confounded by other factors (e.g. genetic or environmental differences between sample donors), we de novo identified enriched DNA sequence-motifs in the top 600 DHSs matching each accessibility profile (Methods). Consistent with unique regulatory functions for DHSs belonging to different accessibility profiles, but not with confounders, distinct combinations of sequence motifs were enriched in each profile, often matching to binding sites of TFs known to be important in the accessible cell types (Fig. 3a). For example, binding sites for the Th2 and Th17 master regulators, GATA3^22^ and RORC^26^, were solely discovered in DHSs specific to Th2 and Th17 cells, respectively. However, we did not recover the binding motifs for the Th1 and Treg master regulators, FOXP3 and TBX21, respectively, suggesting that a lower-resolution was achieved for DHSs specific to these two cell types. Additionally, we found that ubiquitous DHSs were solely enriched for the CCCTC-binding factor (CTCF) – a constitutively expressed DNA-binding protein involved in organizing chromatin into topological domains^27^ – suggesting that these DHSs were marking generic chromatin-to-chromatin contact points found across all cell types.

**Figure 3.**
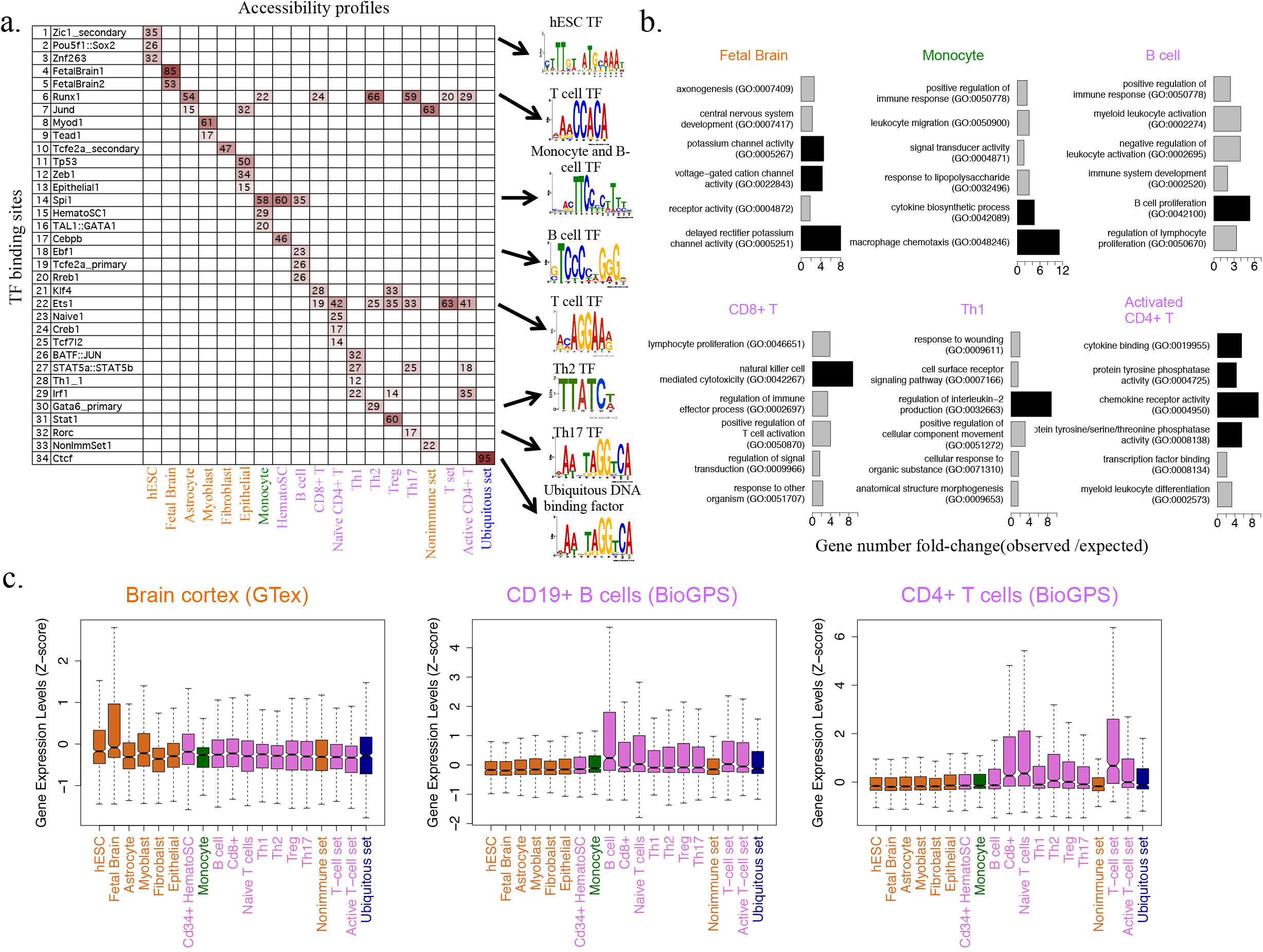
Functional characterization of DHSs grouped by accessibility profiles. **a.** A table summarizing de novo identified DNA sequence motifs (rows) enriched in the top 600 DHSs per accessibility profile (columns). The percent of DHSs containing the enriched motifs is reported. We gave motifs matching known TF binding sites the name of the matching TFs, otherwise, names were reported as the accessibility profile name followed by an index (e.g. see two novel binding sites for fetal-brain-specific DHSs). **b**. We assigned a gene to each DHS by the nearest transcription start site (TSS), and evaluated the top 600 unique genes per accessibility profile for GO term enrichment. The top six enriched terms for six example profiles are shown as barplots. The Bar length marks the fold change (FC) between observed and expected number of genes. Black bars correspond to FC larger than four. Highly enriched GO terms were meaningful and specific to the queried profile. Barplots for all accessibility profiles are presented in Figure S5. **c**. We assigned a gene to each DHS by the nearest TSS, and evaluated the top 600 unique genes per accessibility profile for differential gene expression in profile-related tissues. We show boxplots of *Z* score distributions of gene expression values for each profile (x-axis), in brain cortex, CD19+ B cells, and CD4+ T cells. As can be seen by comparing boxplots, in each sample the most differentially expressed genes came from accessibility profiles of cell types related to the tissue. Similar boxplots for additional tissues are presented in Figure S6 (BioGPS) and Figure S7 (GTEx).

### Accessibility profiles grouped DHSs by coherent regulatory functions

We next determined if DHSs of different accessibility profiles marked regulatory DNA for genes with distinct cellular functions. To this end, we associated DHSs with their nearest transcription-start-site (TSS) gene, as a proxy for regulated genes, and evaluated each profile for gene ontology (GO) term enrichment (Methods)^28,29^. For each accessibility profile, the enriched GO terms were mostly in line with known functions in accessible cell types (Fig. 3b and Fig. S5). Using the BioGPS^30^ and GTEx^31^ gene-expression compendia, we also evaluated if genes assigned to DHSs of a given accessibility profile, where differentially expressed in tissues relevant to accessible cell types (Fig. 3c, Fig. S6, S7). This was indeed the case. For example, in bronchial epithelial cells, genes assigned to epithelial-specific DHSs were most differentially expressed. Additionally, we noted that up regulation in gene expression was much more common than down regulation, indicating that most DHSs marked enhancers.

Taken together, the profile-specific enrichment in TF binding sites, GO terms, and differential gene expression, show that grouping DHSs by accessibility profiles, resulted in grouping of regulatory DNA by coherent, cell-type-dependent regulatory functions.

### GWAS signal stratified by DHS cell-type accessibility profiles

In previous steps, we grouped DHSs into accessibility profiles. We first verified that as expected adaptive-immune-specific DHSs were selectively enriched with risk alleles for autoimmune diseases (Fig. S8). We further resolved which subsets of adaptive-immune-specific DHSs were most relevant to each disease, using a rank-based approach and a permutation-based statistic (Methods; Fig. S9). No single accessibility profile dominated across all autoimmune diseases (Fig. S10a). However, within each disease more strongly enriched cell-type profiles emerged, e.g. DHSs specific to CD8+ T cells, B cells, and Treg cells, were most enriched for systemic lupus erythematosus (SLE). Perhaps of more interest was that Alzheimer’ disease and schizophrenia, two non-autoimmune controls, clustered together with the autoimmune diseases (Fig. S10b). Specifically, we found that schizophrenia was enriched with DHSs specific to CD4+ T cells, and Alzheimer’ was enriched with DHSs specific to monocytes, B cells, and CD8+ T cells, revealing involvement of different immune processes, and providing further resolution to recent results suggesting immune involvement in susceptibility to Alzheimer’s^32,33^ and schizophrenia^34,35^.

### A context-specific, regulatory-wide association study (csRWAS)

Next, we performed an association study between the adaptive-immune-specific DHSs and autoimmune diseases. Specifically, from each GWAS we assigned a single proxy SNP for each DHS, and used that as the DHS association *P*-value (Methods, table S2). Choosing the largest GWAS (by sample size) for each of six autoimmune diseases, we analyzed rheumatoid arthritis^36^ (RA, *n*=80k), ulcerative colitis^37^ (UC, *n*=26k), Crohn’s disease^38^ (CD, *n*=21k), multiple sclerosis^39^ (MS, *n*=15k), systemic lupus erythematosus^40^ (SLE, *n*=11k), and type 1 diabetes^41^ (T1D, *n*=5k). The genome-wide significance typically employed by GWAS (*p*=5x10^-8^) is not appropriate for csRWAS, as all the adaptive-immune-specific DHSs combined constitute ~0.37% of the human genome. Leveraging the groupings of DHSs into accessibility profiles, we estimated a null *P*-values distribution from proxy SNPs assigned to non-immune DHSs (Methods), as we did not expect these to be specifically associated with autoimmune diseases. However, to the extent that non-immune DHSs were tagged by risk alleles, this null is conservative. This allowed matching a *P*-value to a desired FDR threshold. For example, at an FDR of 0.001, P-values ranged from 2x10^-10^ for RA to 3.1x10^-4^ for MS, and at an FDR of 0.005, from 7x10^-5^ for RA to 2.8x10^-3^ for MS. Employing an FDR<0.005 as a significance threshold, we identified between 243 and 839 DHSs (94 to 165 independent loci) per disease. To aid in analysis of these many associated DHSs, we first filtered them based on the GWAS genome-wide significance threshold, as associations above this threshold would have likely already been reported by the corresponding GWAS. Specifically, each associated DHS at FDR<0.005 was assigned to one of four groups, in the following order of precedence:

- ***Genome-wide significant (GW)***, if the associated DHS itself was identified above genome-wide significance (*p*<5x10^-8^).
- ***Locus GW significance (IGW)***, if any GWAS SNP within 0.1cM or 100kb of the associated DHS was identified above genome-wide significance.
- ***True***, if any GWAS SNP within 0.1cM or 100kb of the associated DHS, in the GWAS catalog^4^ for the same disease, was identified above genome-wide significance. Or,
- ***Novel*** otherwise.

That means that DHSs in the Novel and True bins represent associations found below genome-wide significance (with the latter reaching genome-wide significance in other studies). Similarly, after grouping DHSs into independent loci, loci were assigned into these four groups, as determined by the highest-precedence DHS found in each locus.

We initially expected all GW and lGW loci to be reported in the GWAS catalog; after all they had a GWAS SNP with p<5x10^-8^. In practice however we found that this was not the case. For example, out of 644 DHSs assigned to the GW bin (FDR<0.005), 89 were not reported as part of any locus in the catalog. We examined these un-cataloged DHSs manually (Methods) and found that beside a single genome-wide-significance SNP supporting them, typically 0-4 other SNPs in the region were below a nominal significance of p<1x10^-3^. Therefore, these loci were likely flagged as poor in the original GWAS and therefore not reported. Here, we flagged such poorly associated DHSs if around the associated DHS, fewer than four SNPs with *p*<1x10^-3^ were found (Methods). Finally, we determined which DHSs and loci replicated across diseases, as this can provide further proof that an association is genuine, and detect key loci and regulatory DNAs associated to multiple autoimmune diseases (Methods).

We summarized the results for all the non-poor loci, stratified by the above groupings of: GW, lGW, True, and Novel in Table 1 (see table S3 and S4 for details per DHS and per locus, respectively). Out of 529 loci associated to one or more of the autoimmune diseases (FDR<0.005), 322 (60.9%) had further support by either replicating here across diseases (27.2%) or reaching genome-wide significance in the GWAS catalog in the same diseases (55%). Importantly, out of the 529 loci, 327 were discovered below genome-wide significance (True or Novel) in at least one of the diseases a locus was associated with, and 153 (46.8%) of these had replication or catalog support. This shows that using informed regulatory priors allows the identification of many genuine associations below GWAS genome-wide significance; albeit higher significance does lead to improved validation rates (Table 1). It still remains to be evaluated how GWAS compares to csRWAS with respect to replication rates at equal significance thresholds and we will return to this question to conclude the results section. Next, we describe how polymorphisms that disrupt potential TF binding sites inside DHSs were identified, and how we prioritize among them to suggest follow up SNPs for each associated locus.

**Table 1:** Associated loci with cross-replication and GWAS catalog status

| Grouping | a. #Loci | b. #Cataloged | c. #Cross-replicate | d. #(b or c) | e. %Cataloged | f. %Cross-replicate | g. %(e or f) |
| --- | --- | --- | --- | --- | --- | --- | --- |
| GW | 139 | 122 | 71 | 128 | 87.8 | 51.1 | 92.1 |
| IGW | 148 | 119 | 64 | 125 | 80.4 | 43.2 | 84.5 |
| TRUE | 112 | 112 | 42 | 112 | 100 | 37.5 | 100 |
| Novel | 226 | 0 | 52 | 52 | 0 | 23 | 23 |
| **True or Novel** | **327** | **112** | **83** | **153** | **34.3** | **25.4** | **46.8** |
| **All** | **529** | **291** | **144** | **322** | **55** | **27.2** | **60.9** |

We present the number of loci discovered at $FDR < 0.005$ and stratified into four groupings. GW loci had at least one DHS with $p < 5 \times 10^{-8}$ . IGW loci had at least one non-DHS SNP with $p < 5 \times 10^{-8}$ . True loci had no SNP below genome-wide significance here, but reached genome-wide significance when considering all reported studies for that disease in the GWAS catalog. Loci not matching the above criteria were deemed novel. True and Novel loci represent csRWAS associations found below genome-wide significance. Note that cross-replicating loci (loci associated to more than one disease), may belong to more than one grouping, i.e. a locus replicating in RA and SLE, may have been GW with RA, but Novel with SLE. We will show many examples of such cross-replicating loci throughout the text.

### Identifying polymorphisms in DHSs that disrupt predicted TF binding sites

Genetic variants that modify TF binding sites in DHSs are a major source of human gene expression variation^42^. Here, we scanned DHSs for sequences matching one of the enriched TF binding sites found in adaptive-immune-specific accessibility profiles (Fig. 3a), and identified polymorphisms disrupting such binding sites as possible sources of disease risk alleles (Methods). Since many such polymorphic TF binding sites were found, we scored binding sites using a three-letter grade to allow prioritizing among them (Methods). The first letter grade measured how well the predicted binding-site matched to the cognate TF binding site and cell-type context (+A being the best). The second and third letters measured the SNP MAF in Europeans and Asians (A being common, B intermediate, and C rare), respectively (table S5). We will see examples of these grades in action below.

We next describe how we integrated the regulatory and genetic information in a network model.

### A hierarchical integrative network model

Through disease-associated DHSs our approach connected diseases, SNPs, loci, DHSs, polymorphic TF-binding-sites, DHS accessibility profiles, and genes by most proximal TSS. Using the accessibility profiles as a functional guide, we constructed a hierarchical network model with five levels. The network for associated DHSs at an FDR<0.005 is visualized in Figure 4 and is available for navigation in Cytoscape^43^ (supplementary file 1). Although gene assignment to DHSs by nearest TSS is prone to false negatives and false positives, particularly in gene-dense regions, this simple procedure clearly matched many DHSs to correct regulated genes, as supported by the GO term enrichments and differential gene expression analyses, and as shown next by KEGG^44^ pathway enrichment. We first examined the 528 genes in the top level of the network. We asked if the associated genes were enriched with functional pathways, above what was expected for genes most proximal to adaptive-immune-specific DHSs (Methods). The top three enriched KEGG pathways revealed canonical T-cell signaling pathways (hypergeometric test): JAK/STAT signaling pathway^45^ (fold-change[observed/expected]=5.56, *p*=2.86*x*10^-12^), Cytokine-cytokine receptor interaction^45^ (fold-change =4.08, *p*=3.43*x*10^-11^), and T-cell receptor signaling^46^ (fold-change =4.17, *p*=1.65*x*10^-06^). No unexpected pathway enrichments were found. We further examined whether genes found in two or more autoimmune diseases (96 such genes) were more enriched for one of those three pathways. Cytokine-cytokine receptor interaction^45^ and T-cell receptor signaling showed similar fold enrichments (fold-changes of 4.7 and 4.48, respectively), however, JAK/STAT signaling pathway had almost doubled its fold enrichment (fold-change of 10.04, p=9.56x10^-07^) (Fig. 4b). The JAK/STAT genes along with their DHS-derived cellular-contexts were: 1. Janus-kinase 2 (*JAK2*, Activated CD4+ T set^47^), 2. Suppressors of cytokine signaling 1 (*SOCS1*, Activated CD4+ T set^22^), *3*. Signal transducer and activator of transcription 1 (*STAT1*, T set^48^), 4. Protein tyrosine phosphatase non-receptor type 2 (*PTPN2*,T set^49^), 5. Interleukin 23 receptor (*IL23R*, T set or Th17^50^), 6. Interleukin 12 Receptor beta 2 (*IL12RB2*, Th1^51^), 7. Leukemia inhibitory factor (*LIF*, Th1^52^), 8. Sprouty-related EVH1 domain containing 2 (*SPRED2*, Treg), 9. Signal transducer and 10. Activator of transcription 3 *(STAT3*, Treg^22,53^), and interleukin 21 (*IL21*, Th17^54^), most having reported functions in the predicted cell-type context (references provided). Five of these genes form protein-protein complexes through interactions with JAK2, namely: *IL12RB2, STAT3, STAT1* and *SOCS1^22,55^’^57^*. This suggests that in addition to the highly autoimmune-relevant HLA region (which we excluded from all of our analyses), the dysregulation of genes involved in JAK/STAT signaling is the second most prevalent pathway in autoimmunity.

**Figure 4:**
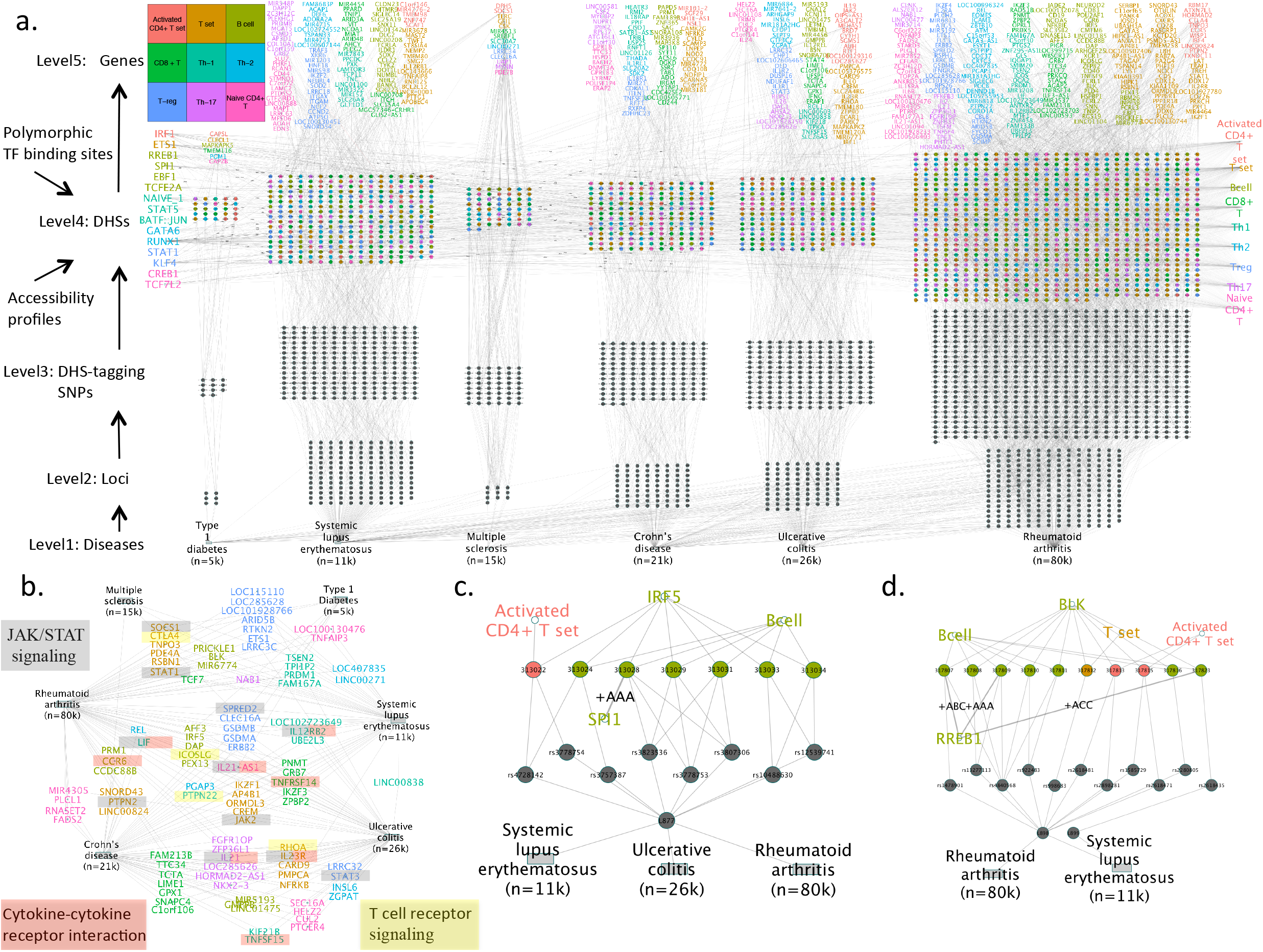
A cross disease hierarchical network model. **a**. We display a hierarchical network model constructed of DHSs associated with six autoimmune diseases at an FDR<0.005. Reading the gene names in the network is possible by either using the digital copy of this manuscript and zooming in, or by navigating the Cytoscape network used to produce this image (supplemental file 1). The network has five levels indicated in the left diagram. DHSs were matched to accessibility profiles (rightmost in the network) and connected to harbored polymorphic TF binding sites (leftmost in the network). Nodes from lower levels connected to nodes in the immediate upper level. DHSs, TF binding sites, and genes were colored to indicate their most well supported accessibility profile (color map on the top left). Loci associated with multiple diseases appear above the associated disease with the largest GWAS sample size. **b**. A disease to gene network of genes associated with two or more diseases (the more diseases a gene was associated with the more central it is). These cross-disease genes were particularly enriched with the JAK/STAT signaling pathway (see main text). **c**. We highlight a sub network generated by selecting *IRF5* as a gene of interest and descending down the network to diseases. Underlying the association is a stretch of B-cell-specific associated DHSs, one of which harbors a good match for SPI1 - a TF binding site enriched in B-cell-specific DHSs. Furthermore, a common SNP in Europeans and Asians disrupt this binding site, providing a testable hypothesis in search of a causal SNP. We graded such good match polymorphic TF binding sites, found in DHSs assigned to accessibility profiles where the TF binding site was also enriched in, and intersected by a common SNP in Europeans and Asians, as ‘+AAA’ to allow prioritizing regulatory polymorphism for follow up work (see methods to better understand our motivation and nomenclature for the three letter grading system). d. We highlight the sub network generated by selecting *BLK* as a gene of interest and descending down the network to diseases. This gene was assigned to a B-cell context, as mainly B-cell-specific associated DHSs were underlying it. Three good motif matches were found for RREB1, a TF binding site enriched in B-cell-specific DHSs - one of which was intersected by a common SNP in Europeans and Asians (+AAA), and two were intersected by less frequent SNPs (scored as +ABC and +ACC). Together, these and other polymorphic TF binding sites provide a prioritizable search space for causal regulatory SNPs.

Note that there is a bias in which cellular contexts we can find. For example, *IL21* was associated with a Th17 context, but is also important for T follicular helper cells^58^, for which we had no accessibility data. Moreover, for T cells we have six lineages, allowing fine resolution into cellular contexts, whereas for B cells we have only one sample, a composite of B cells at different developmental stages, in which case cellular contexts can only be identified as an aggregate.

Next, we provide two examples for how the network can be used to extract actionable insights for two genes of interest, *IRF5* and *BLK* (Fig. 4c,d). For both genes, we show that stretches of B-cell-specific DHSs underlie the association, with at least one of these DHSs harboring a ‘+AAA’ polymorphic TF binding site. Thereby suggesting candidate causal regulatory SNPs, molecular mechanisms of action, and cellular contexts of associations. In support, both genes were previously described as important for B cell development and function^59,60^ and in the same regions super enhancers in B-cells were previously reported^10^.

### Regulatory Manhattan plots (RMPs)

A regulatory Manhattan plot (RMP) highlights DHS tagging SNPs above a set significance threshold, across related traits, and visualizes their matched accessibility profile. The profiles in turn help determine the likely cellular context of the locus. For example, we show the RMPs for the *IRF5* and *BLK* example loci discussed above (Fig. 5). Note that the signal over the *IRF5* TSS region is not in LD with the signal above the *TNPO3* gene body (Fig. S11). As additional examples, we present thirteen associations in three loci that replicated across diseases, *IL2RA, PTPN22*, and *IKZF1*, including associations below GWAS genome-wide significance (True and Novel) (Fig. 6). For *IL2RA* and *IKZF1*, csRWAS resolved associations to a single dominant accessibility profile: *PTPN22* with T set, and *IL2RA* with Treg. Clear evidence that a locus is not associated is also important, so we provide RMPs for all loci across all diseases, including non-associated ones (supplementary files 2 and 3). Such RMPs reveal that csRWAS often could not associate loci with some diseases, not because an association signal was not present, but because no DHS proximal SNPs were available. This highlights the importance of high-quality dense imputations for csRWAS, and means that we could only provide a lower bound for the true cross-replication rates across these six autoimmune diseases.

**Figure 5:**
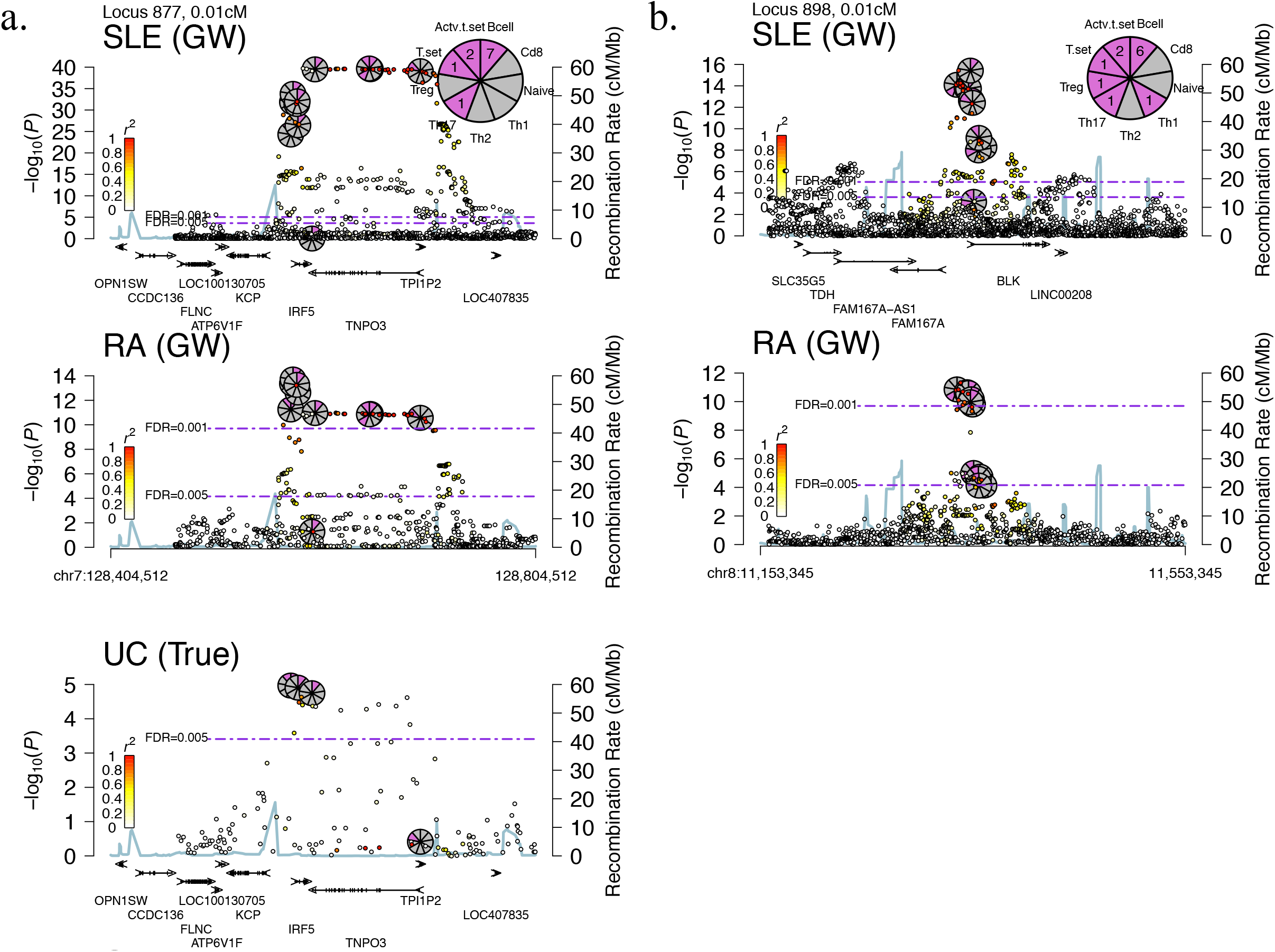
Regulatory Manhattan plots for the *IRF5 and BLK* loci. **a. and b.** We present regulatory Manhattan plots (RMPs) for the *BLK* and *IRF5* loci, complementing the subnetworks in Fig. 4. The input GWAS name is indicated on the top left of each plot together with the loci association bin (GW, lGW, True, or Novel). For cross-disease associations, RMPs were ordered by loci significance (from top to bottom) and centered on the same chromosomal position. DHS tagging SNPs above an FDR threshold of 0.005, in any of the diseases, are visualized as small pie charts with one wedge per accessibility profile, in all of the diseases (as long as a tag SNP was available). Purple wedges indicate the accessibility membership of an associated DHS. A larger summary pie chart that counts the total times each accessibility profile was observed over the entire locus is shown on the top left. Often this summary pie chart can suggest the mostly likely cell-type context for an association. Accessibility profiles in pie charts were ordered as indicated outside the circumference of the summary pie-chart (moving clockwise form 12 o’clock): B cell, CD8+ T cell, Naïve CD4+ T, Th1, Th2, Treg, Th17, T set, and Activated CD4+ T set. For example, for the *IRF5* locus, B-cell-specific accessibility profile was matched by seven associated DHSs, followed by two matches to Activated CD4+ T set profile and a single match to T set and Th17. Therefore, B cell is the predicted cellular context for the *IRF5* locus association. Note that the subnetworks from figures 4c and 4d connect only to the DHSs most proximal to the TSSs of *IRF5* or *BLK*, whereas the Manhattan plots displays all DHSs in a given locus. LD was calculated for all SNPs within 50kb of their nearest DHS tagging SNP. Genes were visualized from their longest Refseq isoform and in cross-disease associations shown under every other RMP.

**Figure 6.**
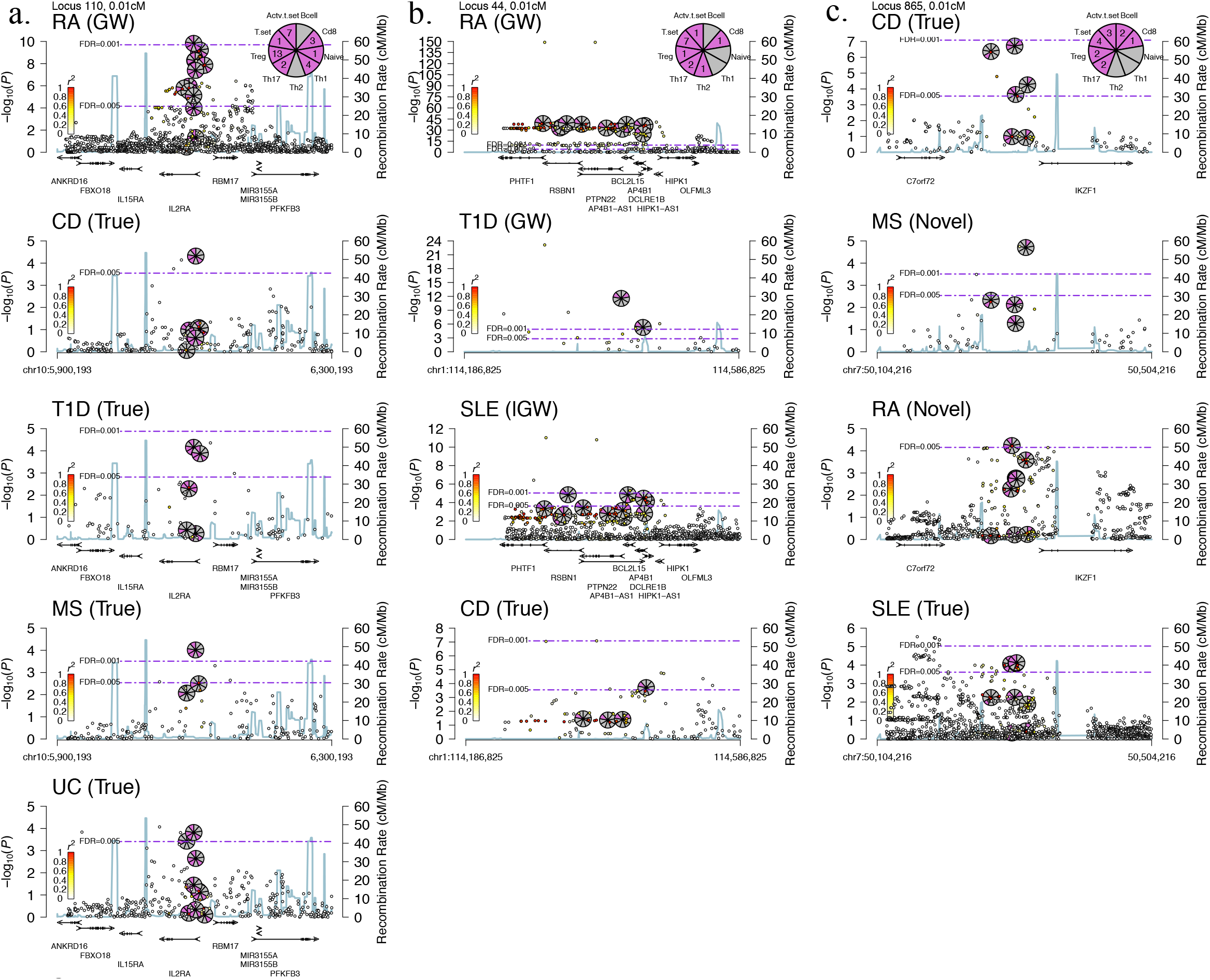
Examples of prevalent loci associated with autoimmunity. We present three loci associated at FDR<0.005 across four or more diseases (13 associations in total). For **a.** and **b.** the cellular context could be resolved to a single most-likely accessibility profile, and five associations were detected below genome-wide significance. For **c.** we report two novel associations in MS and RA upstream to *IKZF1* that replicated in SLE and CD.

In support of the two novel *IKZF1* associations reported above with MS and RA, csRWAS linked three additional members of the ikaros hematopoietic transcription factor family^61^, namely, *IKZF2, IKZF3*, and *IKZF4*, to autoimmune diseases (Fig. S12), further resolving the cellular context of the RA-association with *IKZF4* to Treg, and the RA-UC-CD-SLE-T1D association with *IKZF3* to a T-set or CD8+ T cell accessibility profiles. Note that *IKZF3*, was associated with five of the six autoimmune diseases, suggesting that dysregulation of *IKZF3* is a prevalent cause for autoimmunity, highlighting these TF for possible drug intervention. Additional RMPs for eight loci including the gene-regions for *IL12RB2, TCF7, CTLA4, CD28, CCR6*, and *ETS1* (Fig. S13-S14), reveal more cellular contexts and identify four novel cross-replicating associations, two for *IL12RB2* with RA and SLE, one for *ETS1* (upstream) with SLE, and one for *TCF7* with RA, as well as one non-replicating association for *ETS1* (gene body) with UC. We also note that several genome-wide significant loci identified here were not within 0.1cM of a catalog lead-SNP for the respective diseases, and were not flagged by us as poor. Specifically, two cross-replicating loci were found for RA near *ZFP36L1* and *FAM213B* (Fig. S15), two loci for CD, the first cross-replicating with UC near *BRD7* and the second not-replicating near *TTC33* (Fig. S16), and one locus for MS near *SOCS1* that cross-replicated with T1D and CD (Fig. S17). To summarize, we highlight the more promising novel and replicating loci in Table 2 and Figures S18-S27, providing an array of new insights for six common autoimmune diseases. Furthermore, in order to assist in prioritizing polymorphic TF binding sites for follow up work, each RMP is accompanied by a table (in the same.pdf file) providing additional information similar to that found in the network, but in a tabular format and on the locus scale (supplementary files 2 and 3, see example in Fig. S28). We next contrast the loci replication rates of GWAS and csRWAS to conclude the results section.

**Table 2:**
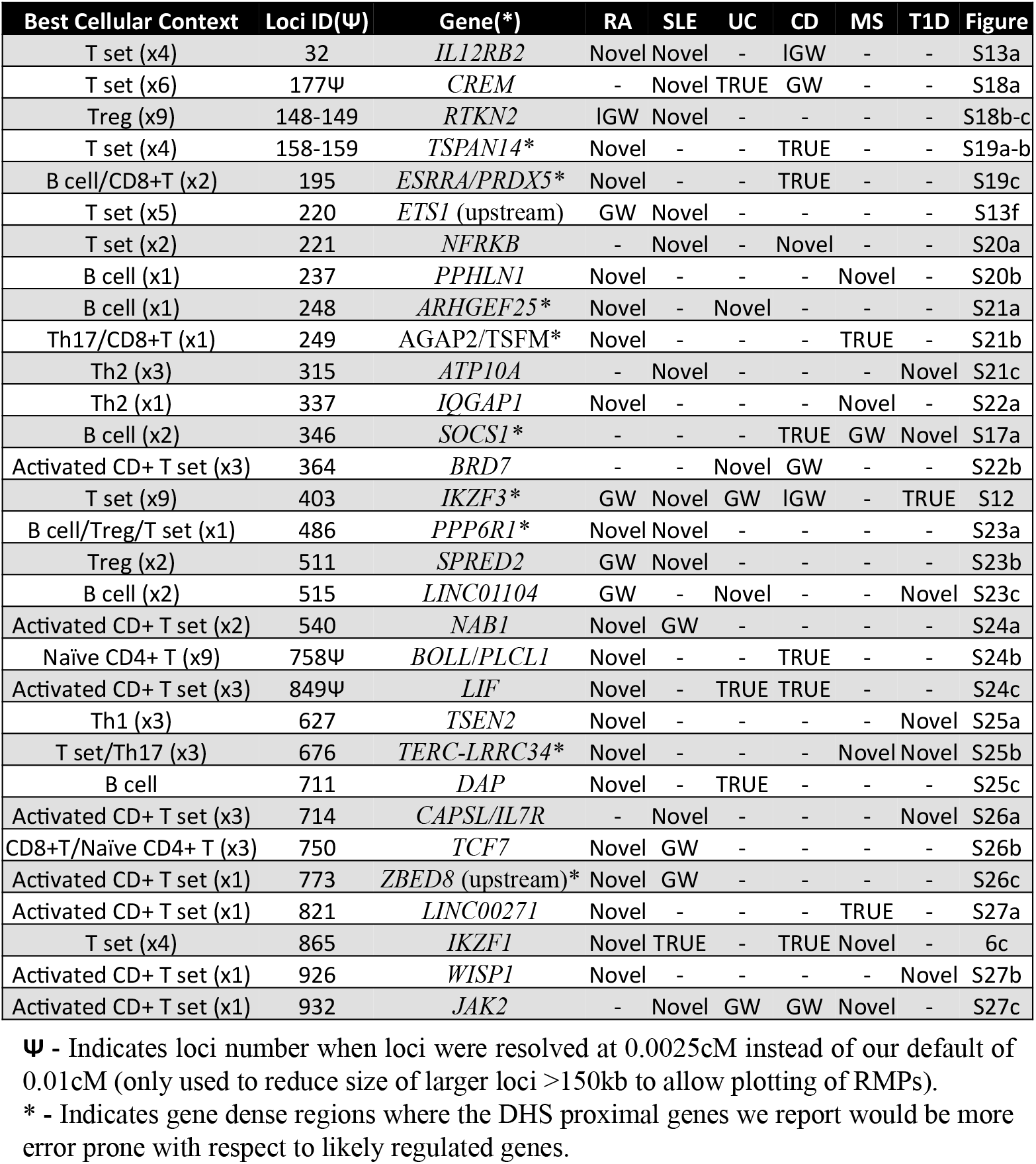
Novel replicating loci, cellular contexts, and DHS proximal genes

| Best Cellular Context | Loci ID($\Psi$) | Gene(*) | RA | SLE | UC | CD | MS | T1D | Figure |
| --- | --- | --- | --- | --- | --- | --- | --- | --- | --- |
| T set (x4) | 32 | *IL12RB2* | Novel | Novel | - | IGW | - | - | S13a |
| T set (x6) | 177Ψ | *CREM* | - | Novel | TRUE | GW | - | - | S18a |
| Treg (x9) | 148-149 | *RTKN2* | IGW | Novel | - | - | - | - | S18b-c |
| T set (x4) | 158-159 | *TSPAN14\** | Novel | - | - | TRUE | - | - | S19a-b |
| B cell/CD8+T (x2) | 195 | *ESRRA/PRDX5\** | Novel | - | - | TRUE | - | - | S19c |
| T set (x5) | 220 | *ETS1* (upstream) | GW | Novel | - | - | - | - | S13f |
| T set (x2) | 221 | *NFRKB* | - | Novel | - | Novel | - | - | S20a |
| B cell (x1) | 237 | *PPHLN1* | Novel | - | - | - | Novel | - | S20b |
| B cell (x1) | 248 | *ARHGEF25\** | Novel | - | Novel | - | - | - | S21a |
| Th17/CD8+T (x1) | 249 | *AGAP2/TSFM\** | Novel | - | - | - | TRUE | - | S21b |
| Th2 (x3) | 315 | *ATP10A* | - | Novel | - | - | - | Novel | S21c |
| Th2 (x1) | 337 | *IQGAP1* | Novel | - | - | - | Novel | - | S22a |
| B cell (x2) | 346 | *SOCS1\** | - | - | - | TRUE | GW | Novel | S17a |
| Activated CD+ T set (x3) | 364 | *BRD7* | - | - | Novel | GW | - | - | S22b |
| T set (x9) | 403 | *IKZF3\** | GW | Novel | GW | IGW | - | TRUE | S12 |
| B cell/Treg/T set (x1) | 486 | *PPP6R1\** | Novel | Novel | - | - | - | - | S23a |
| Treg (x2) | 511 | *SPRED2* | GW | Novel | - | - | - | - | S23b |
| B cell (x2) | 515 | *LINC01104* | GW | - | Novel | - | - | Novel | S23c |
| Activated CD+ T set (x2) | 540 | *NAB1* | Novel | GW | - | - | - | - | S24a |
| Naïve CD4+ T (x9) | 758Ψ | *BOLL/PLCL1* | Novel | - | - | TRUE | - | - | S24b |
| Activated CD+ T set (x3) | 849Ψ | *LIF* | Novel | - | TRUE | TRUE | - | - | S24c |
| Th1 (x3) | 627 | *TSEN2* | Novel | - | - | - | - | Novel | S25a |
| T set/Th17 (x3) | 676 | *TERC-LRRC34\** | Novel | - | - | - | Novel | Novel | S25b |
| B cell | 711 | *DAP* | Novel | - | TRUE | - | - | - | S25c |
| Activated CD+ T set (x3) | 714 | *CAPSL/IL7R* | - | Novel | - | - | - | Novel | S26a |
| CD8+T/Naïve CD4+ T (x3) | 750 | *TCF7* | Novel | GW | - | - | - | - | S26b |
| Activated CD+ T set (x1) | 773 | *ZBED8* (upstream)* | Novel | GW | - | - | - | - | S26c |
| Activated CD+ T set (x1) | 821 | *LINC00271* | Novel | - | - | - | TRUE | - | S27a |
| T set (x4) | 865 | *IKZF1* | Novel | TRUE | - | TRUE | Novel | - | 6c |
| Activated CD+ T set (x1) | 926 | *WISP1* | Novel | - | - | - | - | Novel | S27b |
| Activated CD+ T set (x1) | 932 | *JAK2* | - | Novel | GW | GW | Novel | - | S27c |

$\Psi$ - Indicates loci number when loci were resolved at 0.0025cM instead of our default of 0.01cM (only used to reduce size of larger loci >150kb to allow plotting of RMPs).
\* - Indicates gene dense regions where the DHS proximal genes we report would be more error prone with respect to likely regulated genes.

### High rate of validation for genetic associations identified by csRWAS

Often, an initial GWAS suffers from what is known as the “winner’ curse”, where associations found are likely stronger in that GWAS sample than in the general population that is assayed^62^. This problem is less significant as the GWAS sample size increases. Therefore, the gold standard for validation of any genetic study is replication in an independent sample, which is recommended to be larger to correctly identify false positives^63^. Following these GWAS guidelines, we examined the rate at which loci discovered in a smaller GWAS, replicated in a larger and independent GWAS for the same autoimmune disease. We compared the replication rates attained by GWAS (considering all SNPs), RWAS (considering all DHS tagging SNPs), and csRWAS (considering all adaptive-immune-specific DHS tagging SNPs) (Methods). We show that the replication rates were overall higher for csRWAS when compared to RWAS or GWAS (Fig. 7a,b). The difference in replication rate was more pronounced for the smaller CD discovery data (*n*=4,664), when compared to the larger RA discovery data (*n*=22,515), consistent with prior information having larger impact on underpowered GWAS. Since emerging whole-genome sequencing-based GWAS commonly have such lower sample sizes^64–70^, we propose that csRWAS is well suited for their analysis as well, as it can leverage the improved SNP resolution offered by sequencing data, while avoiding the pitfalls of greatly increasing the number of variants considered.

**Figure 7.**
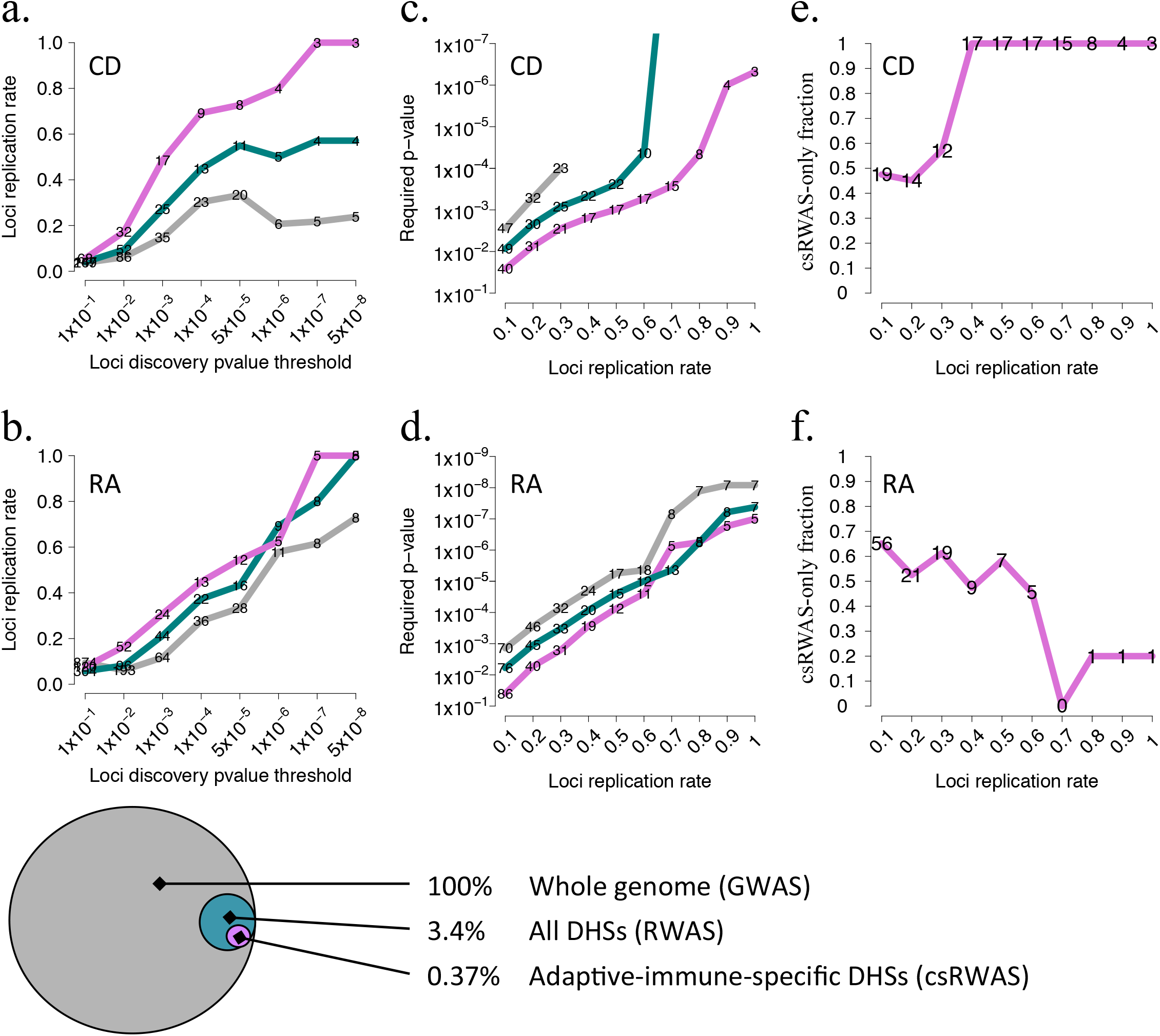
Contrasting the validation rates of csRWAS and GWAS. DHSs were assigned to their nearest GWAS reported SNP. We compared loci replication rates for GWAS (all SNPs, grey), RWAS (all DHS tagging SNPs, blue), and csRWAS (all adaptive-immune-specific DHS tagging SNPs, purple). We considered a GWAS locus as replicating if any of the discovered SNPs within it reached nominal significance in the replication study (*p*<0.01). For csRWAS or RWAS, a locus was considered as replicating if any of its discovered DHS tagging SNPs replicated with such nominal significance. **a. and b.** Loci replication rates as a function of discovery *P* value thresholds for CD and RA, respectively. On the lines we display the number of discovered loci that replicated at each threshold. **c. and d.** The inverse of a. and b., estimating the P-value thresholds required for attaining increasing replication rates. The significance threshold required for csRWAS to attain equal replication rates with GWAS, was often much lower. As before, the entries on the lines specify the number of replicating loci. **e. and f.** The fraction of all csRWAS associations (total csRWAS discoveries) that were not detected by GWAS at equivalent replication rates. The entries on the line specify the number of such discoveries unique to csRWAS.

Considering the reciprocal of the results above, we show that to achieve a desired replication rate, csRWAS required a reduced significance threshold when compared to GWAS (Fig. 7c,d). For example, if the desired replication rate was 0.5 for RA, csRWAS required loci to reach *p*<7.4x10^-05^, while GWAS required loci to reach *p*<5.6x10^-06^. This means that at the intermediate *P*-values (in which significance threshold was reached for csRWAS but not for GWAS) additional loci could be discovered from csRWAS, without loss of accuracy. We quantified the number of such discoveries unique to csRWAS for a range of replication rates (Fig. 7e,f). Keeping with the same example as before, for RA at a 0.5 replication rate, csRWAS discovered 7 additional loci to GWAS, accounting for ~58% of all twelve csRWAS discoveries at that replication rate. The results for CD were more striking, likely because of the higher relevance of priors for underpowered GWAS. These results remained consistent when varying parameters (Methods, Fig. S29-S30).

## Discussion

We have demonstrated that genome-wide association testing can be made considerably more powerful and precise by making use of chromatin accessibility and other functional data, and by focusing on genomic regions relevant to the trait or disease of interest. We illustrate the method by reanalyzing existing genetic data from six autoimmune diseases, where the approach led to additional discoveries and increased interpretability. Note that even if associated DHSs did not harbor any causal allele, the regulatory information they provide can still be used to highlight likely cellular contexts, and therefore help determine candidate genes (e.g. genes with function or high expression in suggested cell types), and guide further experiments to prove causal relationships. We propose that when accessibility data is available for trait-relevant cell types, csRWAS would be a valuable complementary analysis to GWAS.

An important technical advancement made here is in defining a single annotation from multiple DNase-seq samples, and generating a continuous accessibility matrix from multiple DNase-Seq data. This simplifies analysis of large collections of DNase-seq experiments, and can be generalized to other peak-like functional data. This also suggests that a unified DHS annotation from all DNase-seq samples can be defined, facilitating comparative studies of accessibility across a larger accessibility matrix (analogous to GTEx^31^ for gene expression).

As used here, csRWAS had limitations. Some were data related, e.g. not having DNase-seq data for every trait-relevant cell type, or not having biological replicates to estimate the contribution of donor-to-donor variation. As a result of the latter, for example, some of the differences we attributed to cell type specificity, may instead be due to donor-specific genetic or environmental differences. However, the multitude of tests we performed to validate that DHSs grouped by accessibility profiles, also grouped regulatory DNA by coherent regulatory functions, suggests that DHS accessibility was driven primarily by cell-type differences. Other limitations were analysis related, e.g. the TF binding sites we tested here, likely comprise only a fraction of the known and unknown binding motifs important for adaptive-immune cell types. With that respect, the modular design of the analysis makes it possible to incorporate additional DHS related data in the future. Also, although a useful first approximation, the assignment of the most proximal gene’ TSS to DHSs as a likely target was error prone. As such, when examining a locus of interest we recommend carefully reviewing the RMP and information table for that locus, and applying domain knowledge before making a decision on candidate genes for follow up.

Extensions to this work are also possible. For example, the measured trait, instead of being disease status, could be gene expression in relevant cell types^71^. The large number of studied “traits” generally further diminishes power of such expression quantitative trait loci (eQTL) mapping. Hence, focusing on regulatory DNA specific to trait-relevant cell types is promising for greatly improving both statistical power and functional insights of eQTL studies.

To conclude, we identified 1975 DHSs (FDR<0.005) associated to one or more of six autoimmune diseases, grouped into 529 independent loci across the genome. The original GWAS of these six diseases, employing standard genome-wide significance thresholds and not integrating non-genetic data, would have missed 327 of these loci, although 153 of those (46.8%) readily replicated here or in the GWAS catalog. Also, as a result of the functional prior, this study identified and replicated 52 novel loci. But perhaps more importantly, csRWAS provides actionable insights for many associations, e.g. suggests polymorphic TF binding sites only in accessible DNA unique to trait-relevant cell types, and proposes cellular contexts for follow up experiments.

Taken together we presented compelling evidence that trait-tailored approaches to functional and genetic data, can provide a structured path to better understanding how genotypes are translated to phenotypes.

## Acknowledgements

We thank Leonardo Arbiza, Kaixiong Ye, and Cris Van Hout for helpful comments and discussions about this work. This work was supported in part by NIH grant R01- HG006849 (A.K.) and by an award from the Ellison Medical Foundation (A.K.). This work would not have been possible without the sharing of data through public resources. We thank the Encyclopedia of DNA Elements (ENCODE) consortium and Roadmap Epigenomics consortium and particularly the work carried out by the Stamatoyannopoulos lab for collecting the DNaseI-seq data used in this manuscript. We thank the following consortia for sharing GWAS summary statistics: Rheumatoid Arthritis Consortium International for Immunochip (RACI), International Inflammatory Bowel Disease Genetics Consortium (IIBDGC), The Coronary Artery Disease Genetics Consortium (C4D), International Consortium for Blood Pressure (ICBP), DIAbetes Genetics Replication And Meta-analysis (DIAGRAM), The Genetic Investigation of ANthropometric Traits (GIANT), International Genomics of Alzheimer’ Project (IGAP), Psychiatric Genomics Consortium (PGC), and the Euro-Canadian systemic lupus erythematosus consortium (EC-SLE). This study makes use of data generated by the Wellcome Trust Case Control Consortium (WTCCC). A full list of the investigators who contributed to the generation of the data is available from www.wtccc.org.uk. Funding for the WTCCC project was provided by the Wellcome Trust under award 076113. Finally, we would like to thank all the participants and staff involved in the collection of the genotype-phenotype data used in this manuscript.

## Supplemental Figures

**Figure S1.**
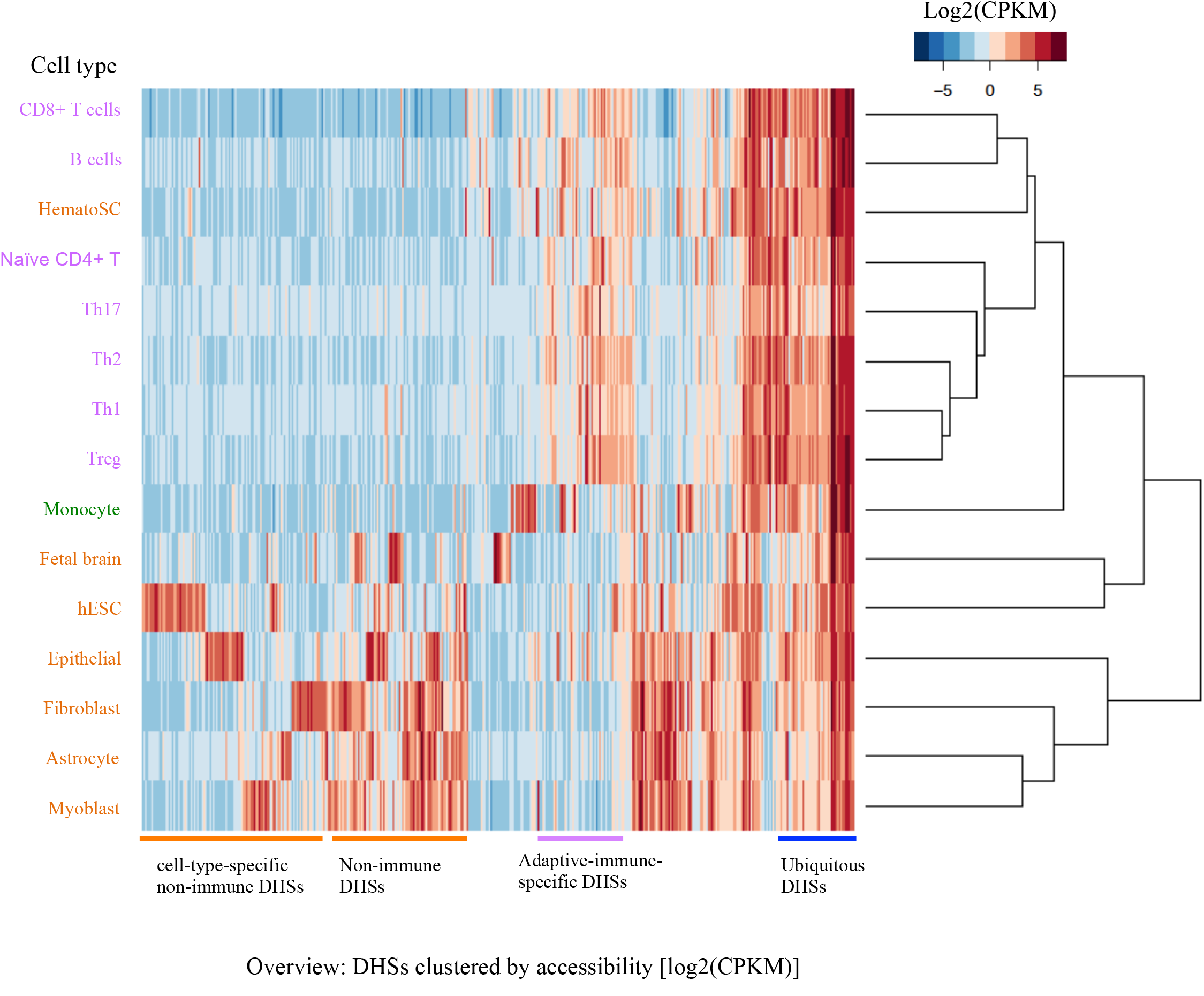
DHS accessibility across fifteen cell types. Accessibility heatmap for a random subset of 30,000 DHSs measured across fifteen cell types. The cell types (rows) largely clustered based on their expected developmental relationships. The innate immune Monocytes (green) serves as a developmentally related control for the adaptive immune cell types (purple). The non-immune cell types (orange) serve as more distant controls. Colored lines demarcating four robust accessibility profiles are shown below the heatmap. Accessibility was measured in cleavages per kilobase per million (CPKMs).

**Figure S2.**
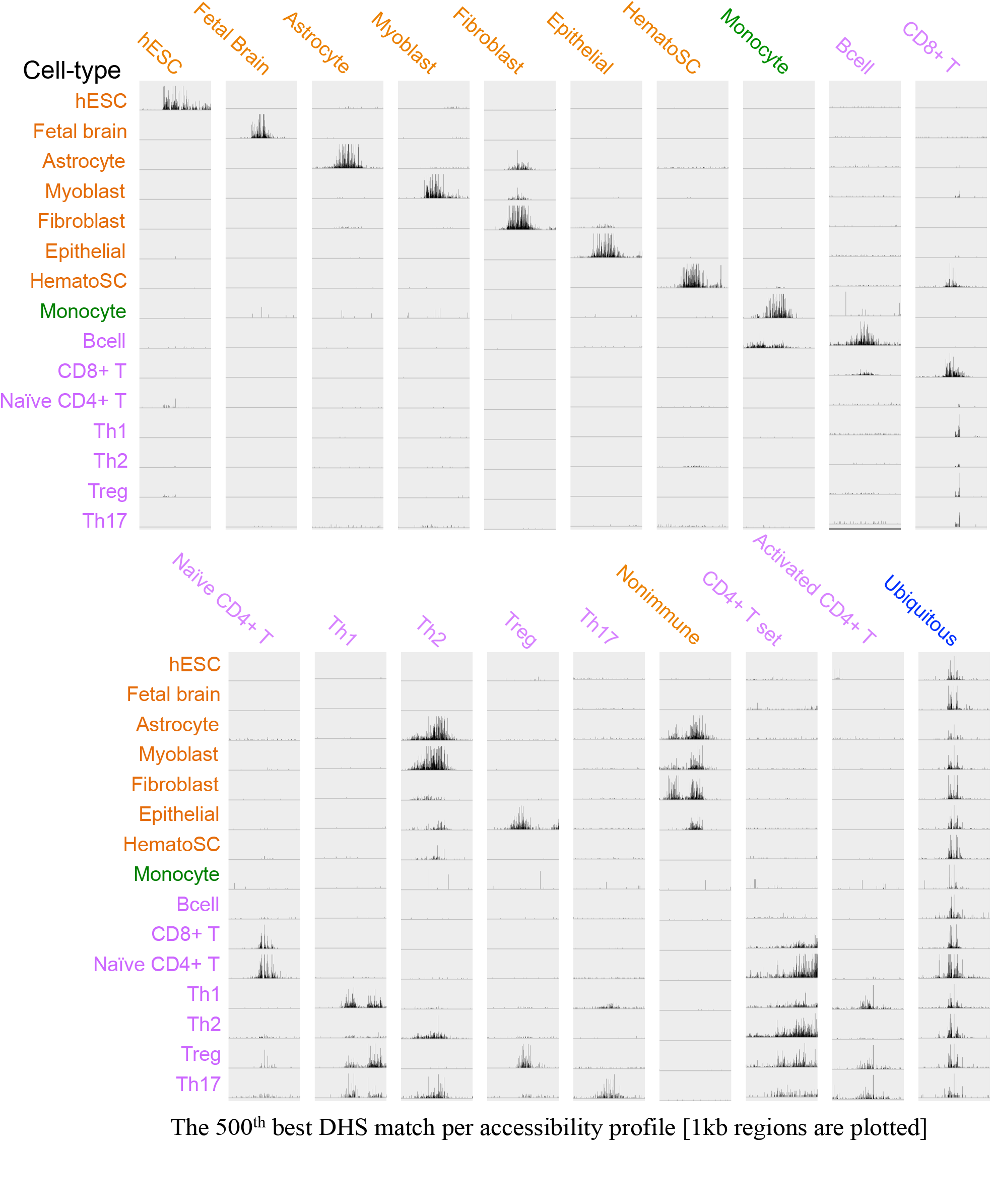
Genome browser view of an example DHS assigned to each accessibility profile. An IGV genome browser view (Broad Institute) of a 1kb DNA region surrounding the midpoint of the DHSs ranking as the top 500^th^ match per accessibility profile. For each panel the track height was fixed to a single value to allow comparisons across all cell types. Reads per millions were used as input to IGV to normalize for inter-sample variations in sequencing depth.

**Figure S3.**
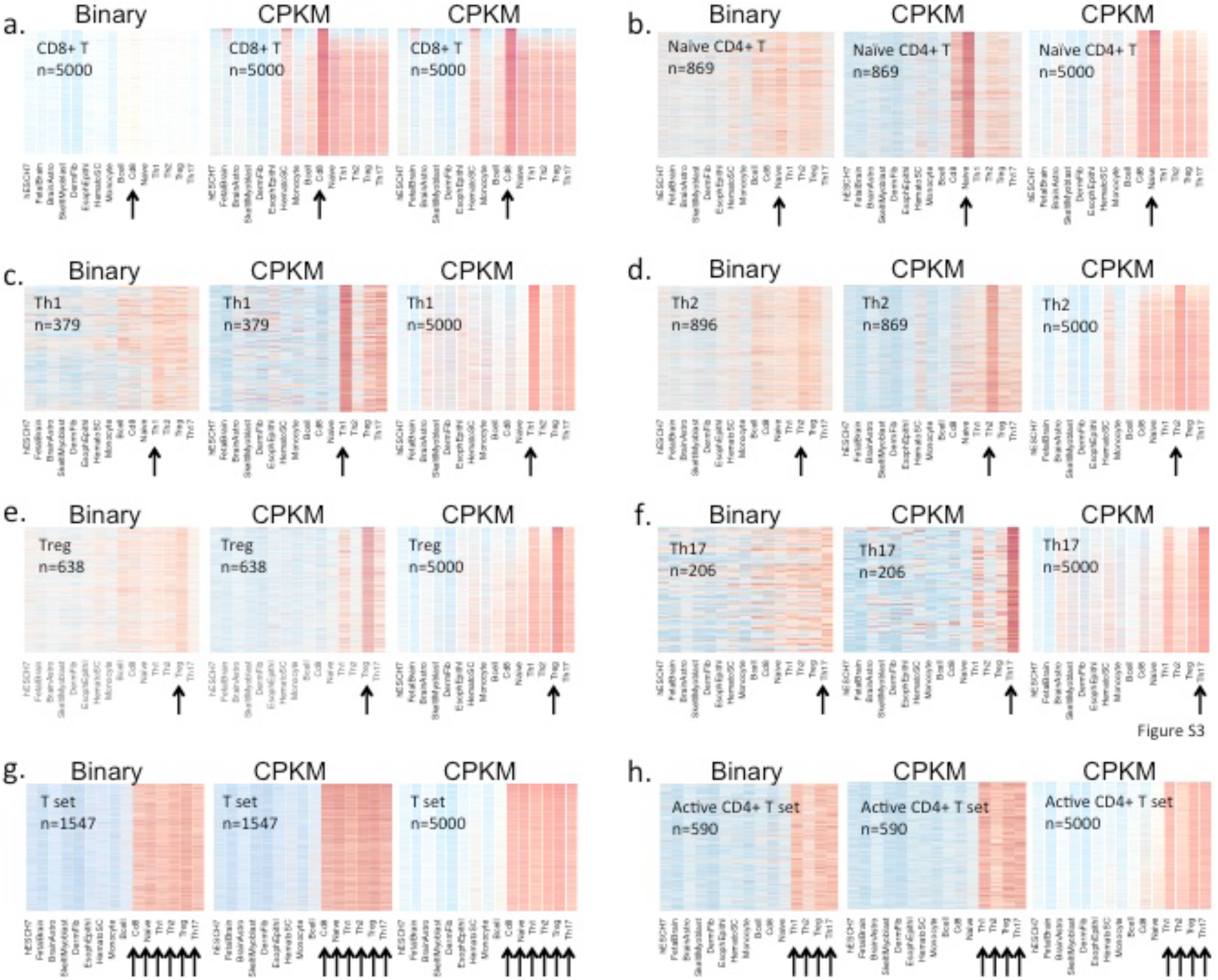
Continuous vs. binary accessibility matrices: Specificity and sensitivity of resolving T-lineage-specific DHSs. **a-h.** For each of the eight T-lineage related accessibility profile, namely: **a.** CD8+ T cell, **b.** Naïve CD4+ T cell, **c.** Th1, **d.** Th2, **e.** Treg, **f.** Th17, **g.** T set, and **h.** Activated CD4+ T set, we assigned DHSs based on the CPKM or binary matrices (Methods). For each profile we show three heatmaps for the identified DHSs based on the binary matrix (**left**), an equal number of DHSs based on the CPKM matrix (**middle**), and the top 5000 DHSs based on the CPKM matrix (**right**). The number of DHSs plotted is indicated on each heatmap. For the CD8+ T-cell accessibility profile the binary approach assigned 6386 DHSs, in that case we plotted the first 5000 of these (there is no notion of top DHSs from binary data). Black arrows mark the cell types comprising each accessibility profile, where one would expect to see the highest accessibility signal. DHSs assigned by the binary matrix showed lower specificity, as seen by lower accessibility in cell types comprising each accessibility profile when compared to other cell types (compare left to middle heatmaps). Furthermore, many more DHSs, of visually better or equal accuracy, were identified from the continuous matrix (compare left with right heatmaps).

**Figure S4.**
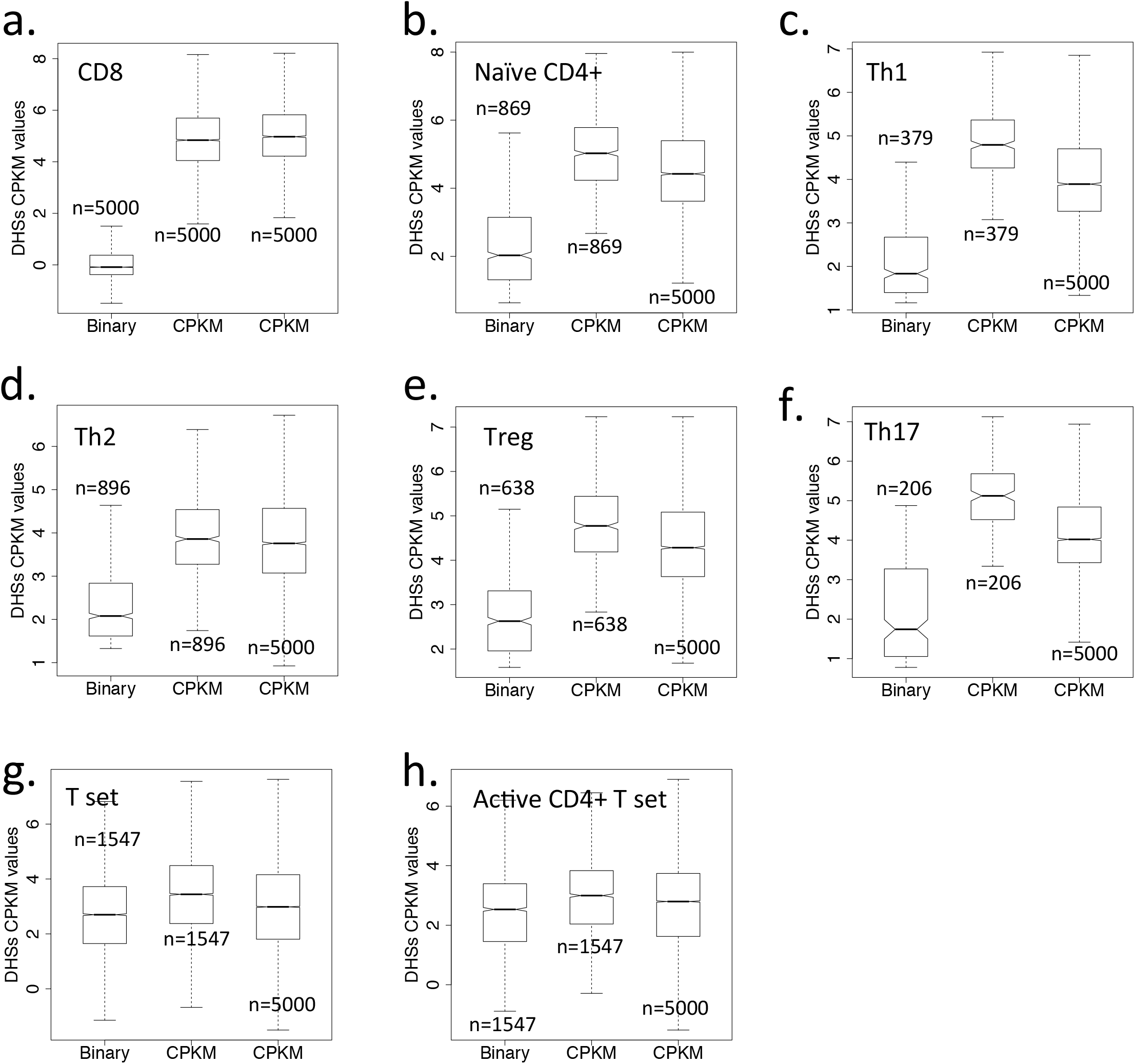
Continuous vs. binary accessibility matrices: Signal to noise of resolving T-lineage-specific DHSs. **a-h.** For each of the eight T-lineage related accessibility profile, namely: **a.** CD8+ T cell, **b.** Naïve CD4+ T cell, **c.** Th1, **d.** Th2, **e.** Treg, **f.** Th17, **g.** T set, and **h.** Activated CD4+ T set, we assigned DHSs based on the CPKM or binary matrices (Methods). For each profile, we show three CPKM boxplots of: DHSs identified from the binary matrix (**left**), an equal number of top DHSs based on the CPKM approach (**middle**), and the top 5000 DHSs based on the CPKM approach (**right**). In each boxplot we show DHS accessibility values from cell types comprising that accessibility profile (i.e. cell types and DHSs marked by black arrows on the corresponding heatmaps in Figure S3). The number of DHSs in each boxplot is shown above or below the boxplots. DHSs assigned by the binary matrix showed lower accessibility when compared to equal or even larger number of DHSs assigned by the CPKM approach, indicating that a lower signal to noise ratio is afforded by the binary accessibility matrix.

**Figure S5.**
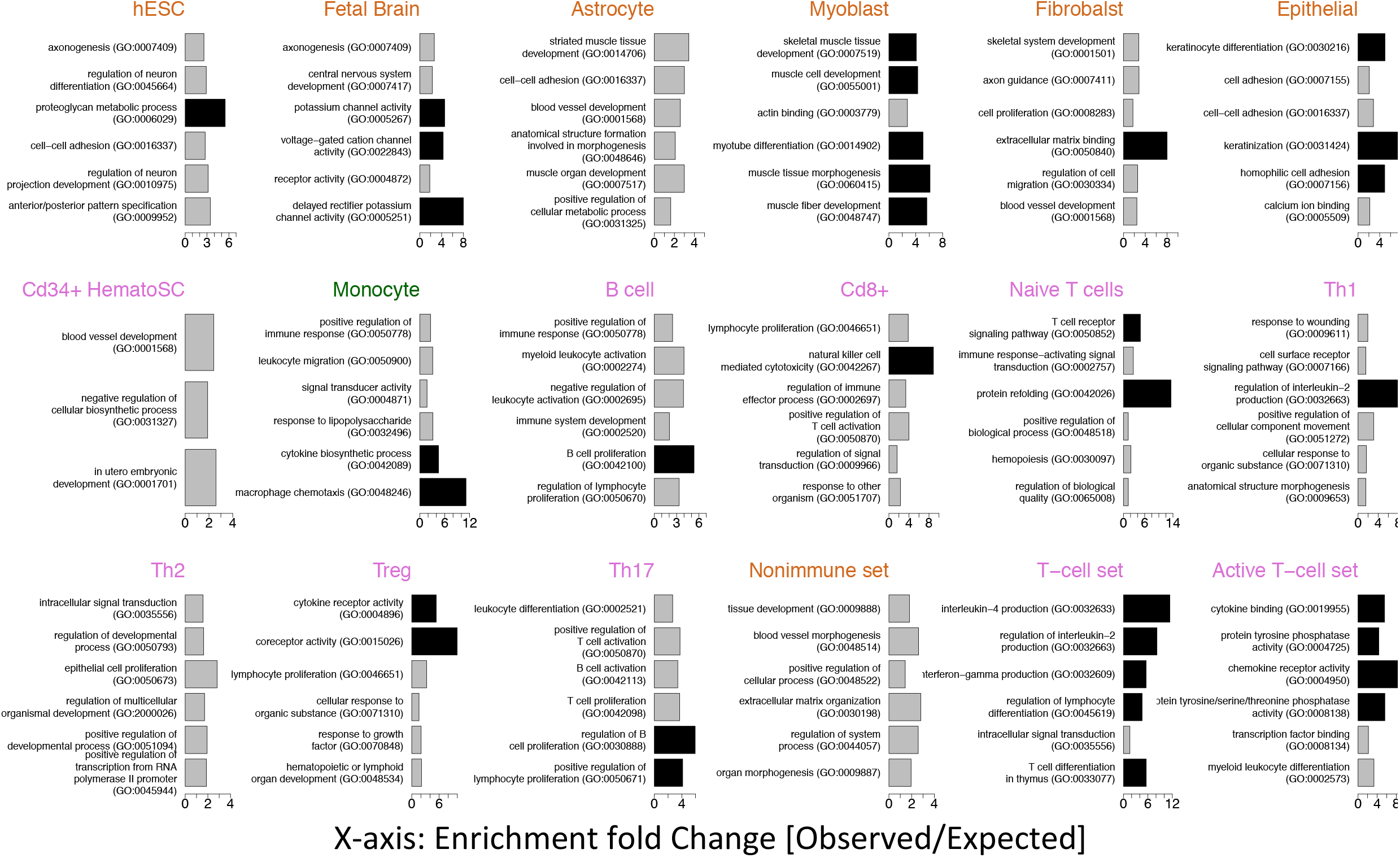
GO term enrichments per accessibility profile. We assigned a gene to each DHS by the nearest TSS, and evaluated the top 600 unique genes per accessibility profile for enrichment with Gene Ontology (GO) terms. The top six enriched terms per profile are shown as barplots. The Bar length marks the fold change (FC) between observed and expected number of genes. Black bars correspond to FC larger than four. There were no GO terms enriched for the ubiquitous set profile. Most enriched GO terms were meaningful and specific to the queried profile: e.g. positive regulation of muscle tissue morphogenesis for Myoblasts specific DHSs, B cell proliferation for the B-cell specific DHSs, natural killer cell mediated cytotoxicity for CD8+ T cell specific DHSs, regulation of interleukin-2 production for Th1 specific DHSs, coreceptor activity for Treg specific DHSs, and chemokine receptor activity for the Activated CD4+ T set DHSs.

**Figure S6.**
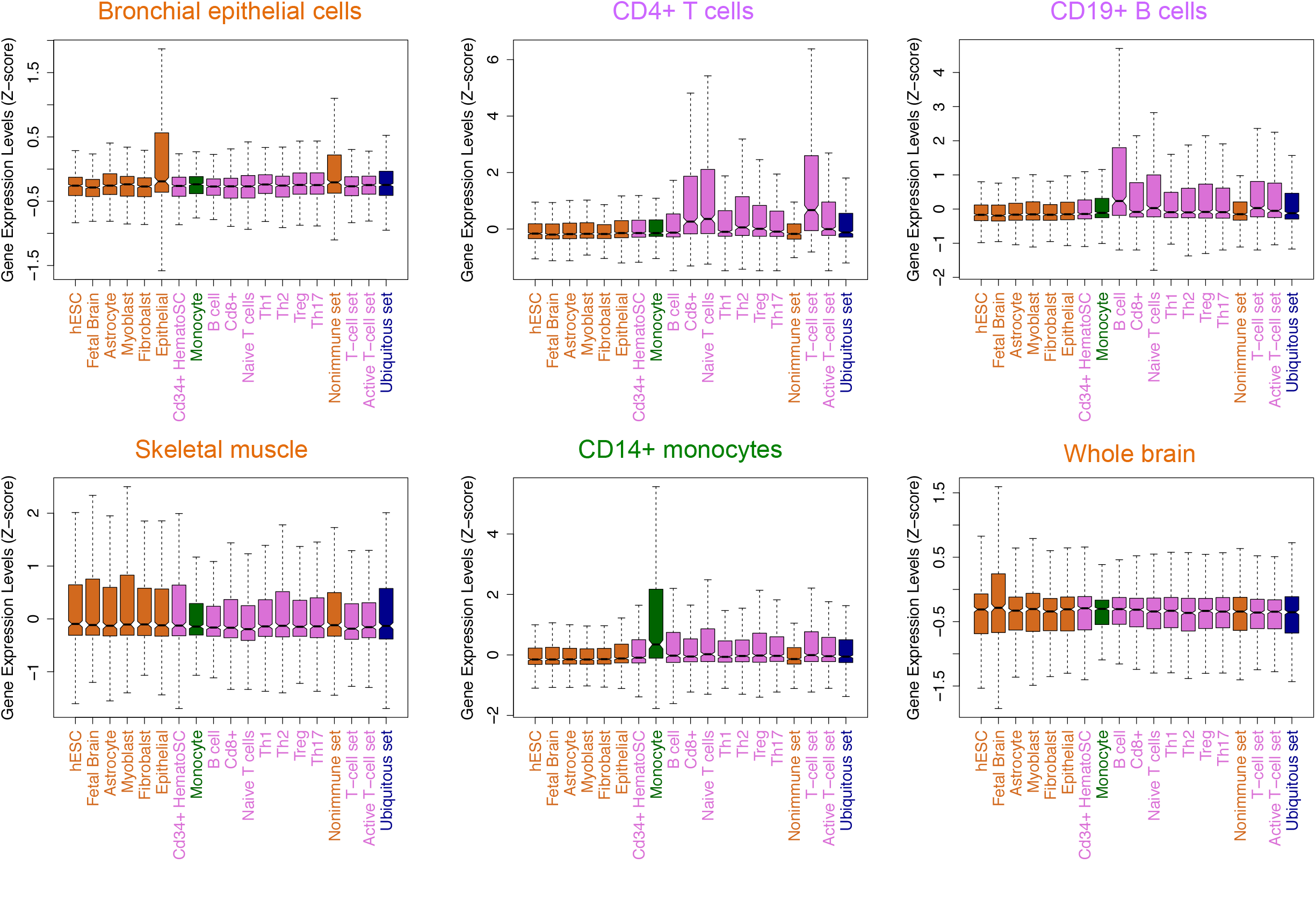
BioGPS: gene expression as a function of accessibility profiles. We assigned a gene to each DHS by the nearest TSS and evaluated the top 600 unique genes per accessibility profile for differential gene expression in profile-related tissues. We show a boxplot for the *Z*-score distribution of gene expression values in each profile (x-axis), for six BioGPS samples. For each sample, the most differentially expressed genes (as can be seen by comparing boxplots) came from accessibility profiles directly related to that tissue sample. For example, genes most proximal to epithelial specific DHSs were most differential in the Bronchial epithelial BioGPS sample.

**Figure S7.**
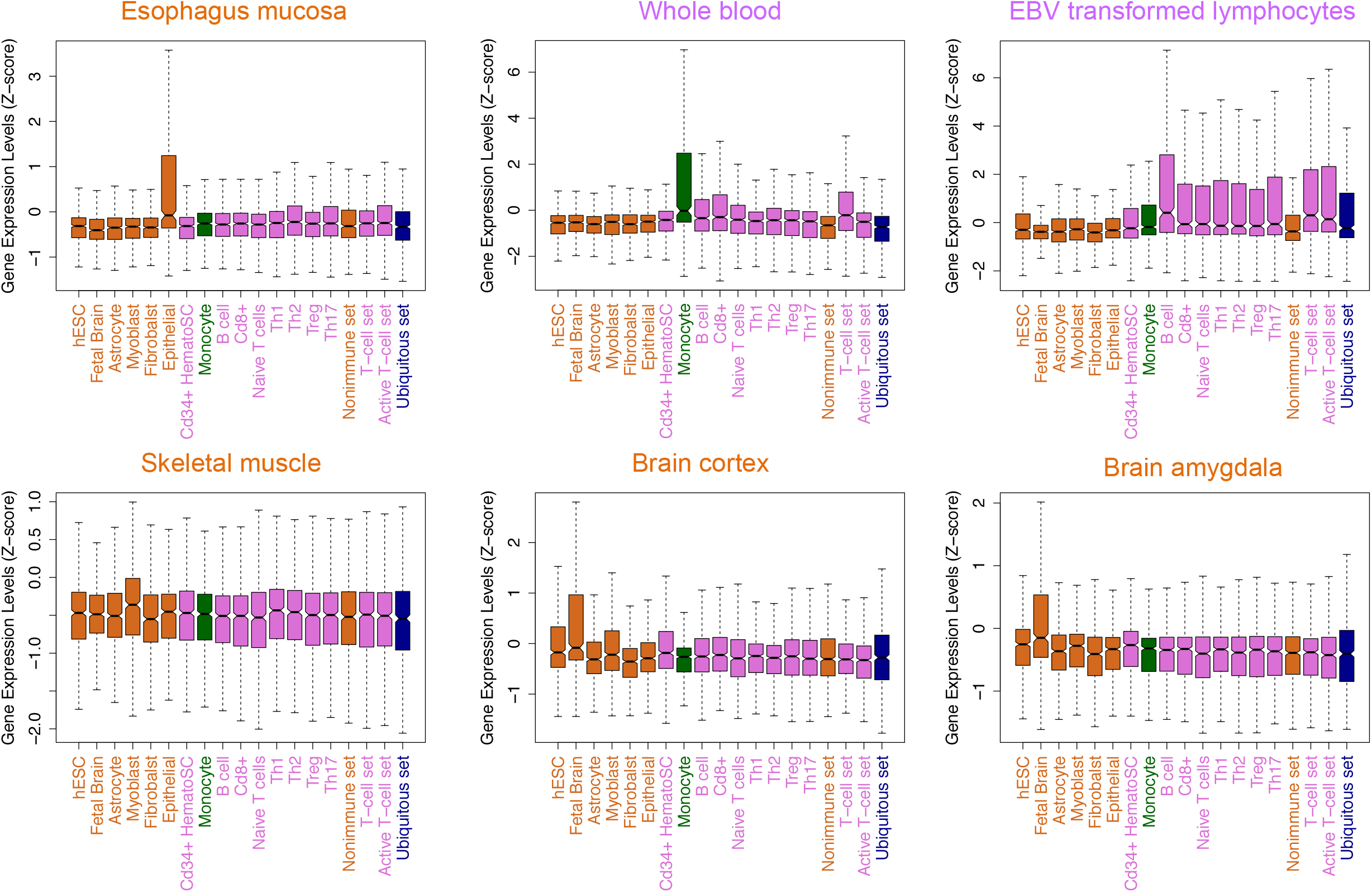
GTEx: gene expression as a function of accessibility profiles. We assigned a gene to each DHS by the nearest TSS and evaluated the top 600 unique genes per accessibility profile for differential gene expression in profile-related tissues. We show a boxplot for the *Z*-score distribution of gene expression values in each profile (x-axis), for six GTEx samples. For each sample, the most differentially expressed genes (as can be seen by comparing boxplots) came from accessibility profiles directly related to that tissue sample. For example, genes most proximal to fetal brain specific DHSs were most differential in the brain cortex and amygdala GTEx samples.

**Figure S8.**
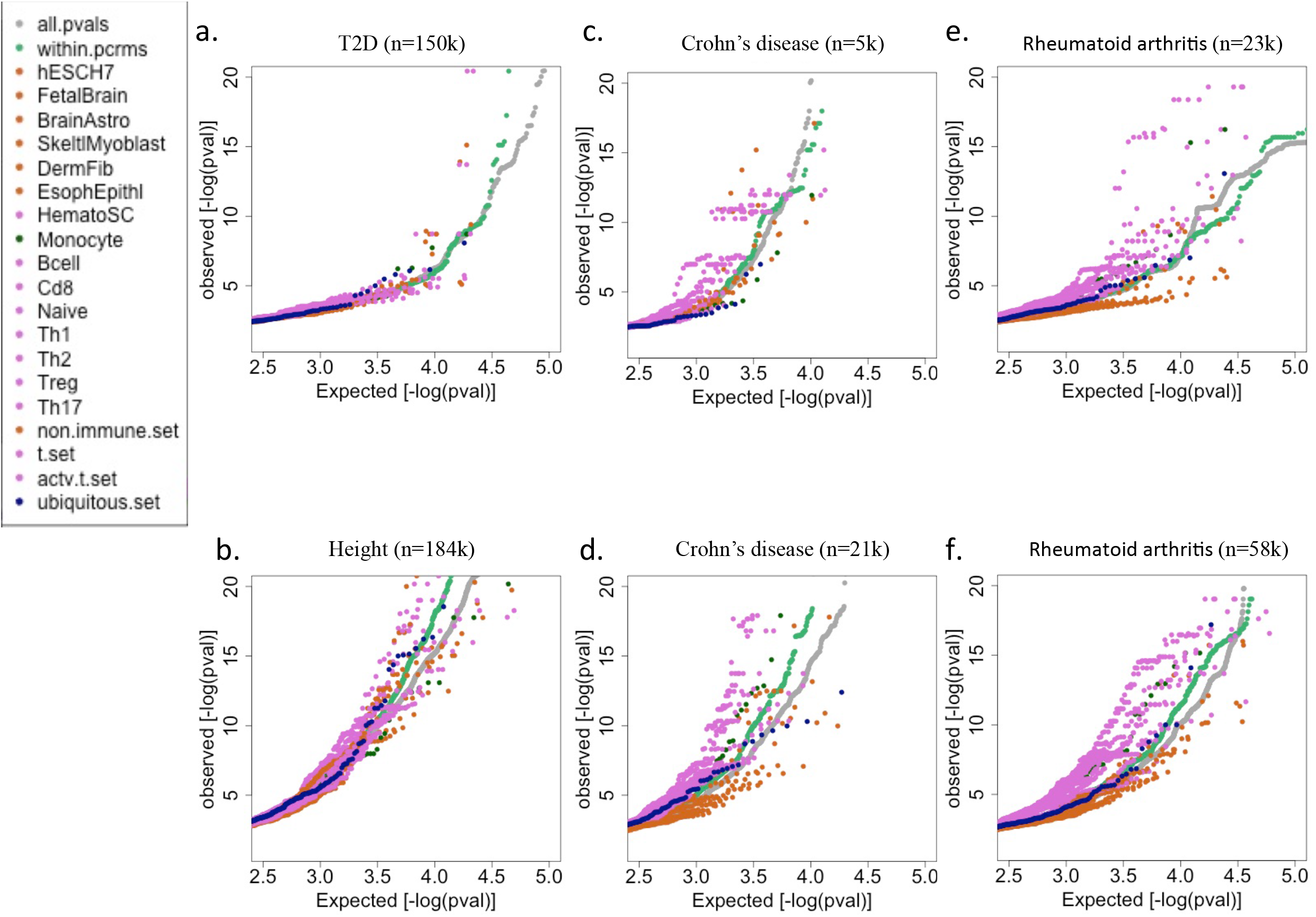
QQ plots stratified by DHS accessibility profiles. DHSs were assigned the nearest GWAS SNP and retained if a SNP was found within 2.5kb of their midpoint. We compared QQ plots for all GWAS SNPs (grey), all DHS SNPs (green), DHS SNPs matching non-immune accessibility profiles (orange), DHS SNPs matching adaptive-immune-specific profiles (purple), and DHS SNPs matching the ubiquitous set (blue). **a. and b.** QQ plots for two non-autoimmune GWAS of height and type 2 diabetes (T2D) serving as controls. We found the adaptive-immune-specific DHS SNPs (purple) distributed above and below all DHS SNPs (green), showing no selective enrichment. **c. and d.** Two independent (no shared participants) GWAS QQ plots for Crohn’ disease. In both cases, adaptive-immune-specific DHSs (purple) were enriched with GWAS, low *P*-value SNPs (i.e. purple above the green dots) and non-immune DHSs were depleted (orange below green dots). **e. and,f.** Two independent GWAS QQ plots for rheumatoid arthritis. As for CD, adaptive-immune-specific DHSs were enriched with GWAS, low *P*-value SNPs and non-immune DHSs were depleted. GWAS samples sizes are indicated in parentheses.

**Figure S9.**
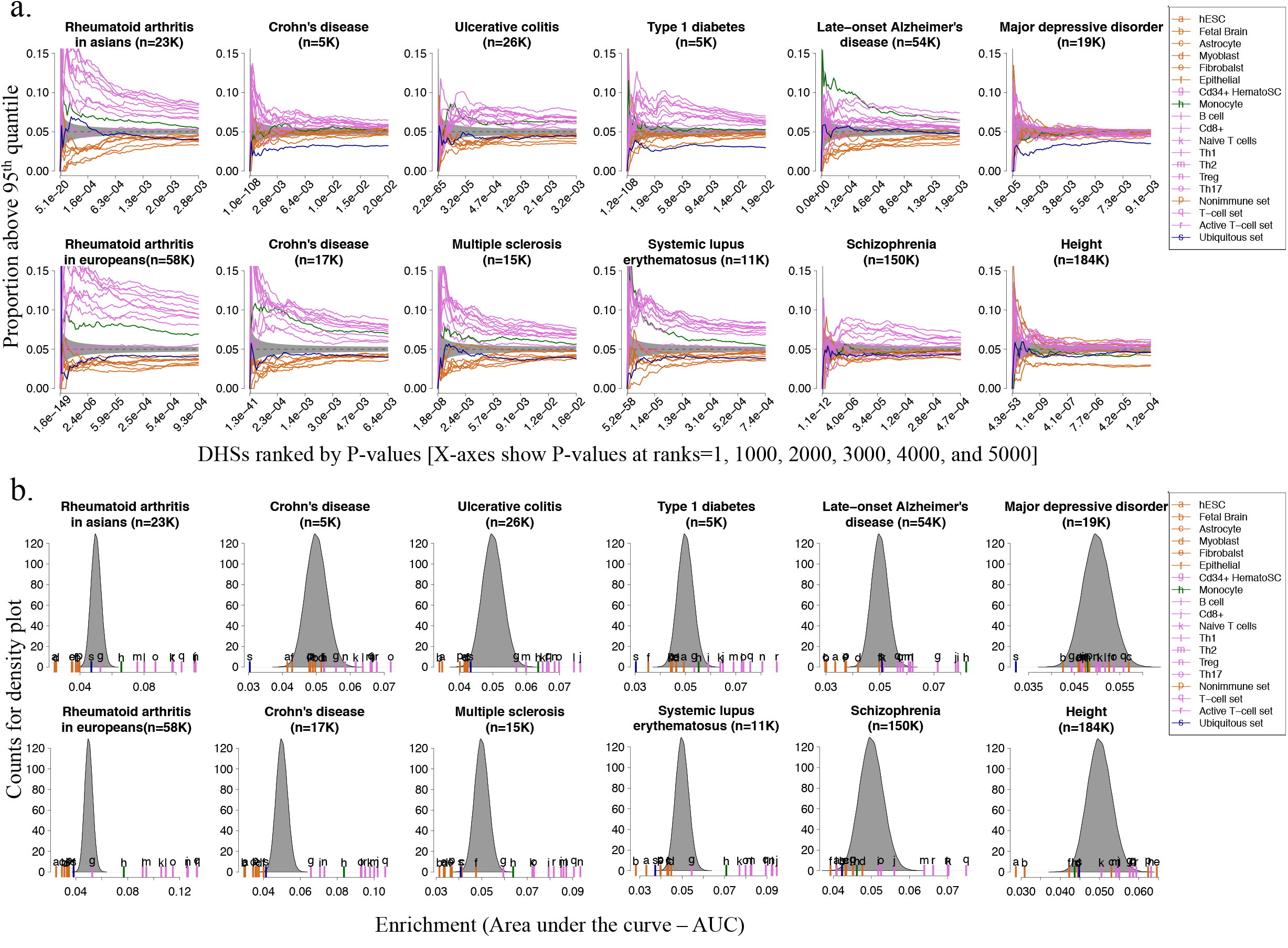
Enrichment of accessibility profiles among low *P*-value DHSs. For each GWAS, DHSs were assigned the minimum GWAS SNP *P*-value, found among all SNPs within 2.5kb from DHS midpoints. If a DHS had no GWAS reported SNP within 2.5kb, it was discarded from the analysis of that GWAS. **a.** Per GWAS, the top 5000 DHSs with minimum SNP *P*-values were ordered increasingly on the x-axis. For each accessibility profile we plotted an enrichment curve defined as the cumulative proportion of DHSs with a profile match score in the top 5 percent of all match scores for that profile (Methods). Hence, the cumulative proportion is expected to distribute around 0.05, where we also added a one standard deviation in grey (based on binomial sampling). Curves above or below this grey shaded area indicate profiles that were enriched or depleted, respectively. **b.** For each GWAS and profile, we calculated a summary statistic defined as the area under the enrichment curve (AUC), with the total possible area (the square of the full plot) normalized to 1. The expected AUC distribution (grey) was estimated from 10,000 simulations (Methods). Short colored lines and letters above them indicate the observed AUCs for each accessibility profile.

**Figure S10.**
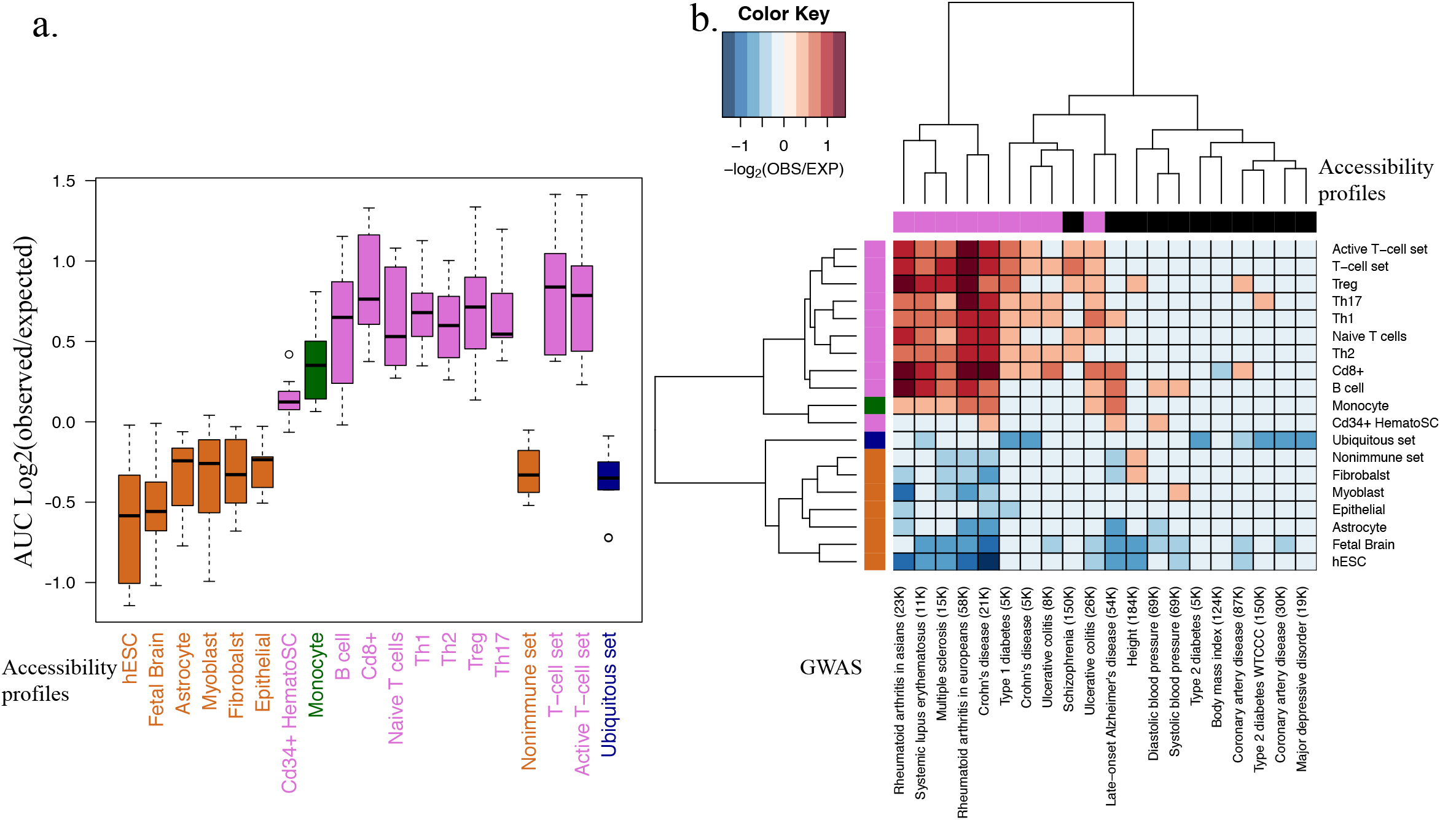
Summary statistics of accessibility profiles among low *P*-value DHSs. Accessibility profile enrichments were measured using the AUC summary statistic, as described in Fig. S9. **a.** Profiles that were enriched or depleted after Bonferonni correction (*P*-value < 0.05) were visualized in a heatmap with the heat color matched to the Log2 fold-change (observed/expected) of the AUCs. Top and side dendrograms were based on clustering all fold-change values, including values from non-Bonferonni significant entries. **b.** We visualize the distribution of Log2 fold-changes for all autoimmune diseases as boxplots (one per accessibility profile). Non-adaptive-immune DHSs were depleted with associations (orange below zero). Adaptive-immune-specific DHSs were enriched with associations (purple above zero).

**Figure S11.**
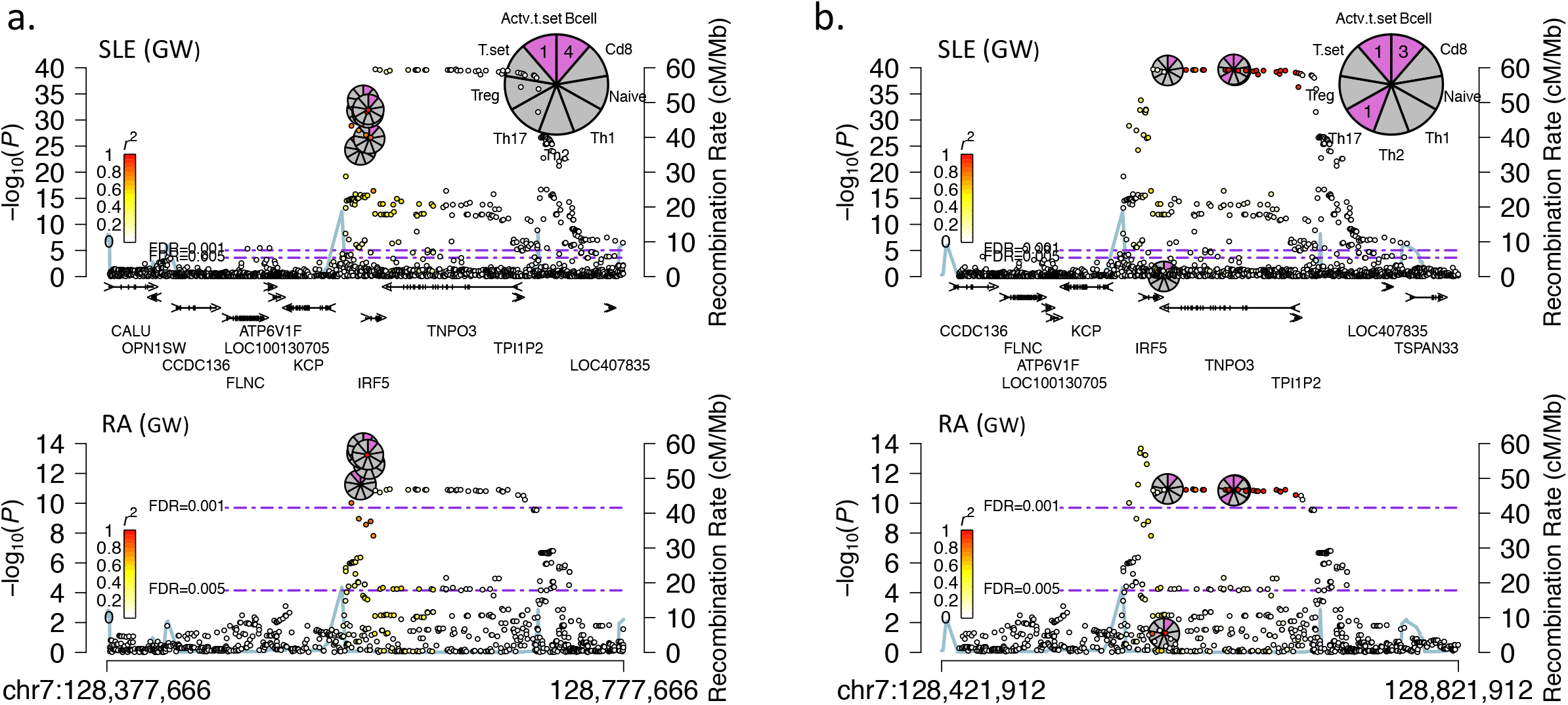
The *IRF5* TSS signal is not in LD with the *TNPO3* gene-body signal. Linkage disequilibrium, as measured in r^2^ from the CEU samples in the 1000 Genomes Project, is visualized using a white-yellow-red gradient, showing that the TSS signal for *IRF5* (**a.**) was LD independent from the signal over the *TNPO3* gene body (**b.**). In both cases r^2^ was visualized for all SNPs within 50kb of their nearest associated DHS-tag SNP. These RMPs were generated using the alternative 0.0025cM cutoff for building loci, to allow breaking down of larger loci annotated with the 0.01cM cutoff (supplementary file 3).

**Figure S12.**
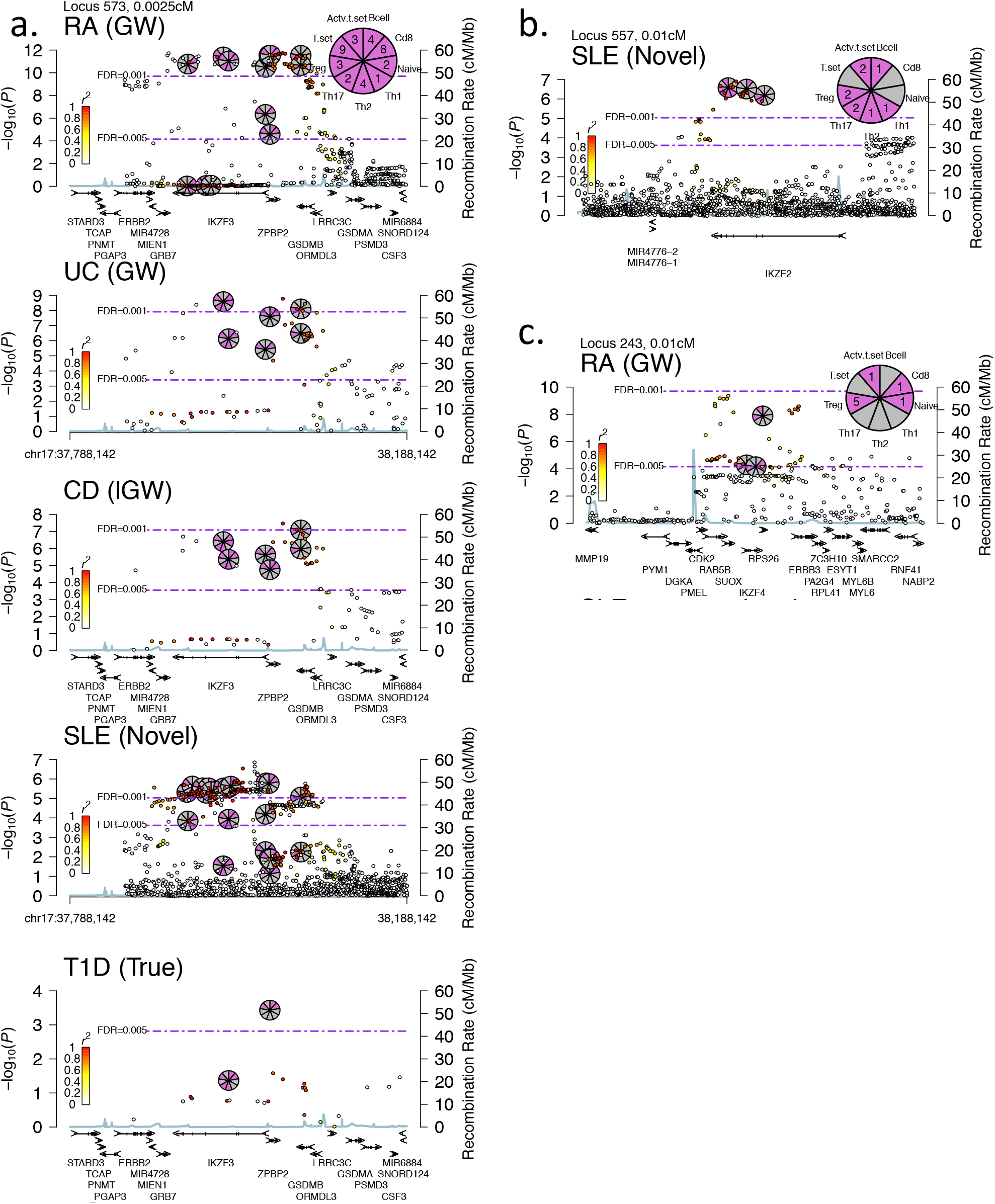
RMPs for autoimmune associations with the ikaros hematopoietic transcription factors: *IKZF2, IKZF3*, and *IKZF5*. Supplementing the association found for *IKZF1*. We present RMPs for the associations made with autoimmune diseases for three other members of the Ikaros transcription factor family. **a.** RMPs for the highly cross-replicating association found for *IKZF3* (T set or CD8+ T-cell context). **b.** RMP for the association found for *IKZF2* with SLE. **c.** RMP for the association found for *IKZF4* with RA (Treg context).

**Figure S13.**
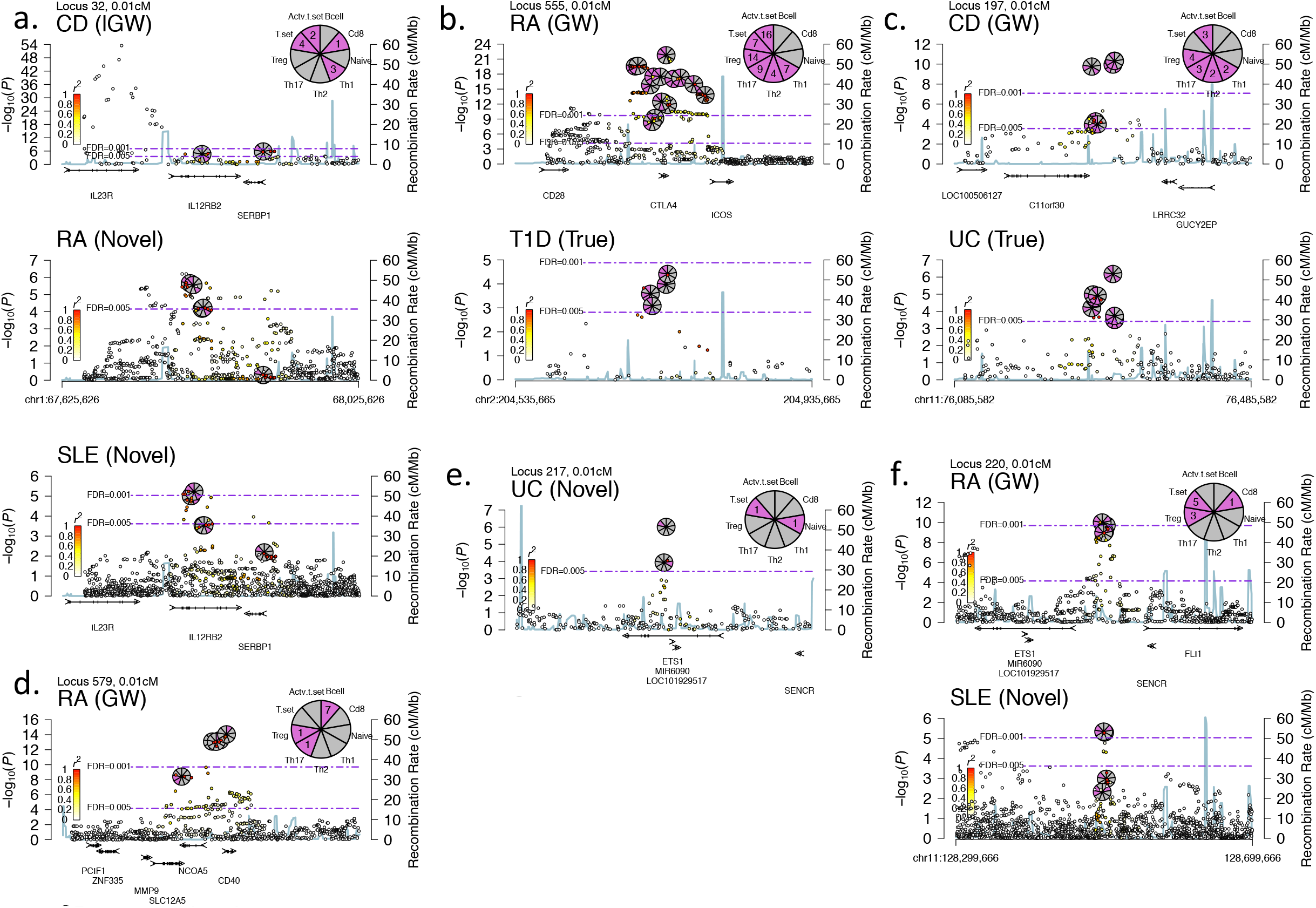
Novel, True, and genome-wide significance associations for *IL12RB2, CTLA4, LRRC32, CD40*, and *ETS1*. We present RMPs for eleven associations made in five gene regions. Five of the associations were GW, two were True, and four were Novel. Two of the novel associations were for *IL12RB2* with RA and SLE (**a.**), which replicated with each other and with CD, one for *ETS1* (upstream) with SLE, which replicated in RA, and one was non-replicating association for *ETS1* (gene body, independent of the upstream signal) with UC (**e.**). Accessibility profiles provided further resolution into the cellular context of several of these associations, namely: *CTLA4* to Treg or Active CD4+ T set (**b.**), *CD40* to B cells (**d.**), and *ETS1* to T-set or Treg accessibility profiles (**f.**).

**Figure S14.**
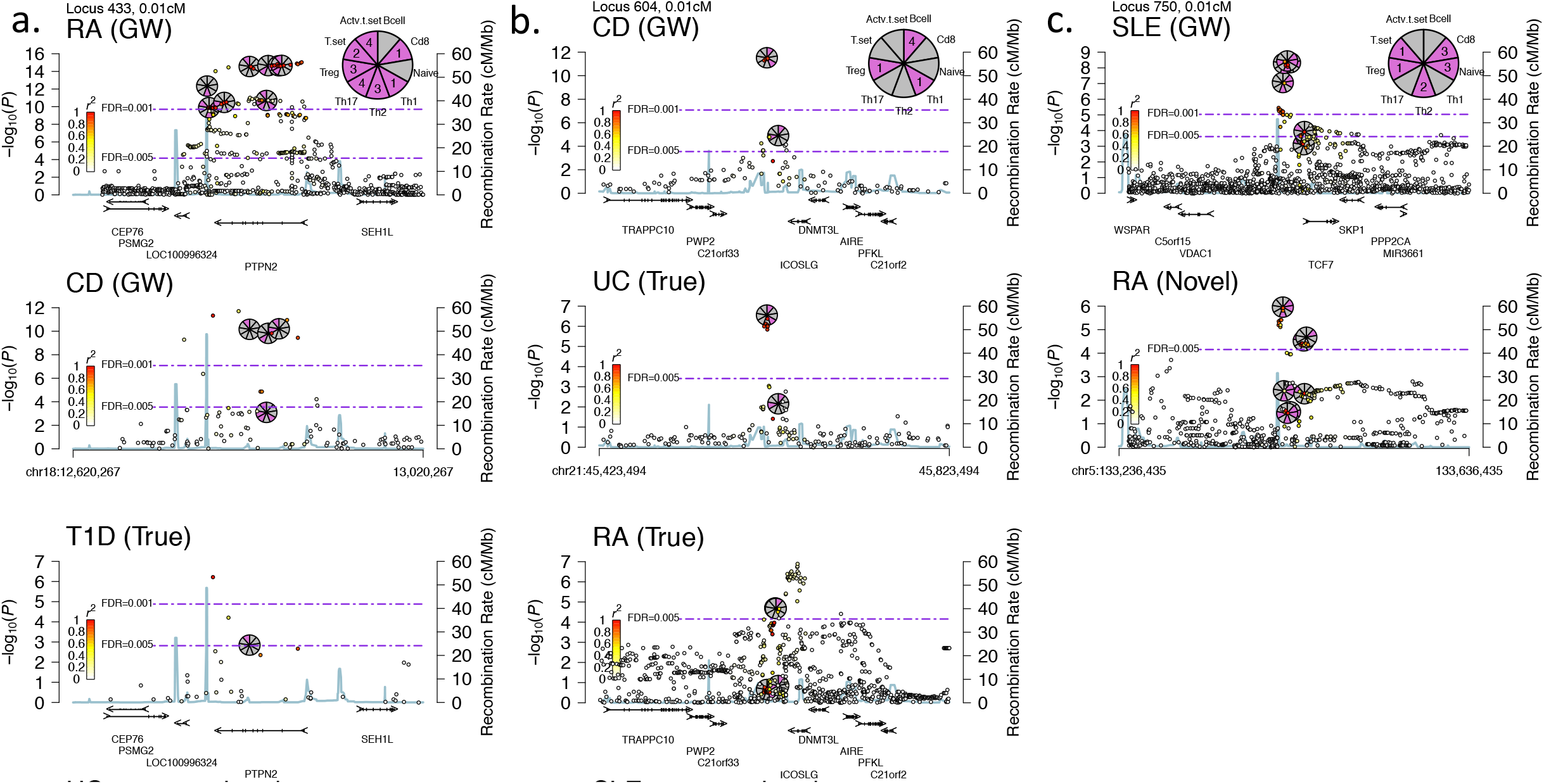
Novel, True, and genome-wide associations for *PTPN2, ICOSLG*, and *TCF7*. We present RMPs for eight associations made in three gene regions. Four of the associations were GW, three were True, and one was Novel for *TCF7* with RA that replicated in SLE (**c.**). Accessibility profiles suggest that the cellular context of the *PTPN2* association is one of the activated CD4+ T cells, namely: Th1, Th2, Th17, and Treg, or all together, as in the Activated CD4+ T set (**a.**), *ICOSLG* is B cells (**b.**), and *TCF7* is CD8+ and naïve CD4+ T cells (**c.**).

**Figure S15.**
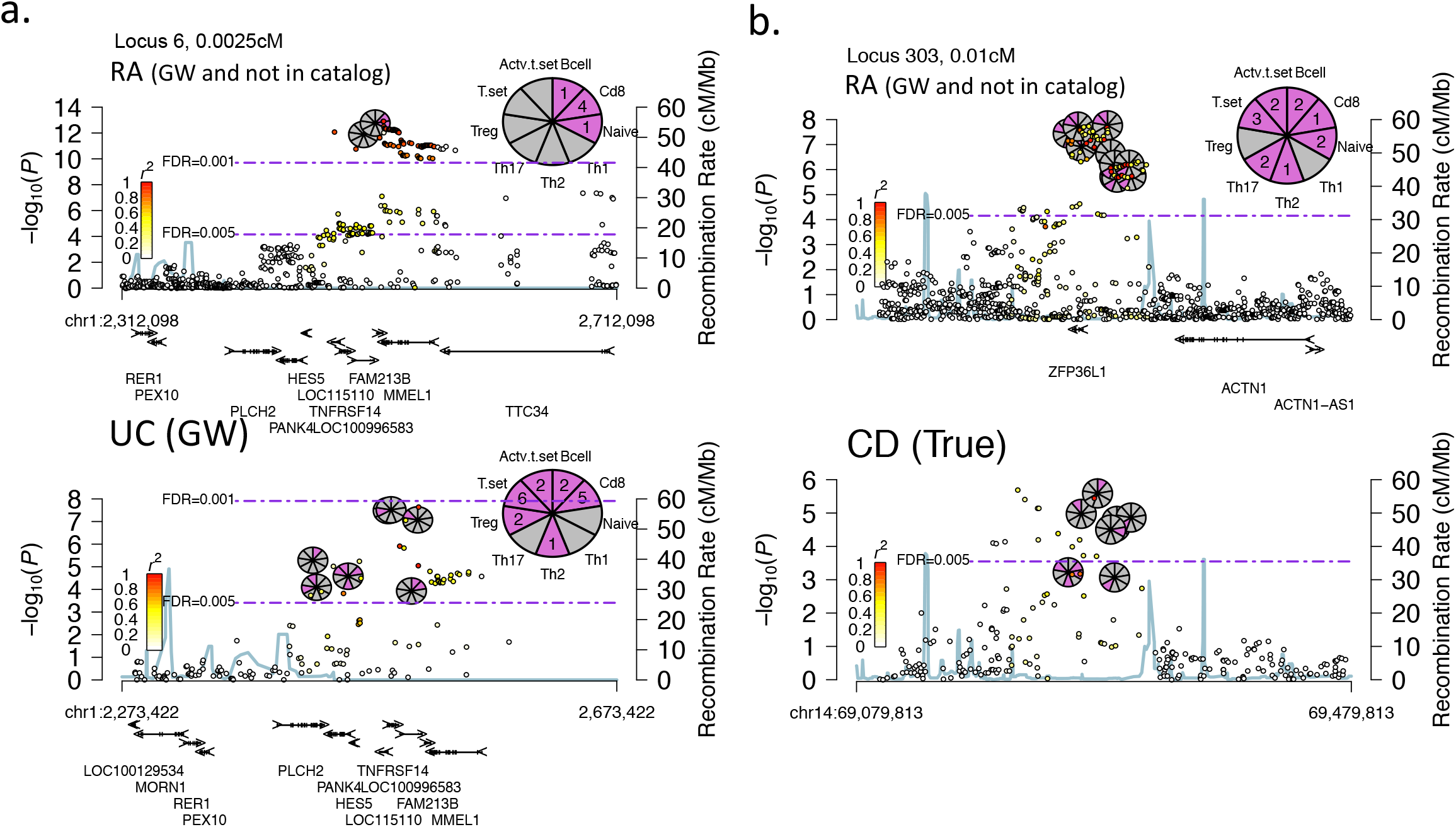
Two rheumatoid arthritis loci with genome-wide significance not near a GWAS catalog, lead SNP. We present RMPs for two RA associations of genome-wide significance, which were not within 0.1cM of a GWAS cataloged, lead SNP for RA. **a.** The first locus surrounding *FAM213B* (CD8+ T cell context) was reported by the authors of the original GWAS, and cross-replicated here with UC. Therefore, this association should be inserted into the catalog. **b.** The second locus surrounding *ZFP36L1* was not reported by the authors and not found in the catalog, however it looks genuine to us given that it reached genome-wide significance, and cross replicates here with CD.

**Figure S16.**
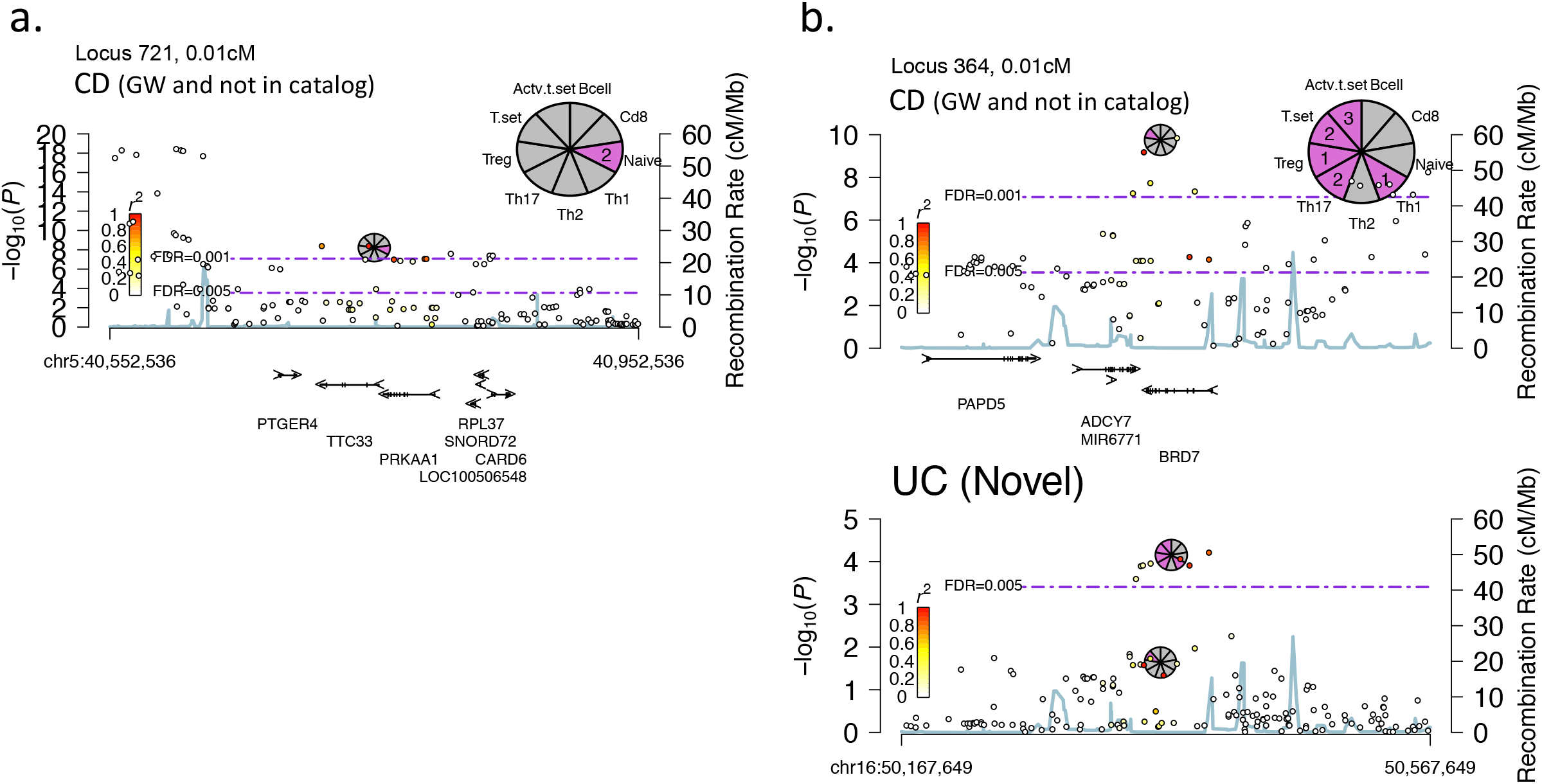
Two Crohn’ disease loci with genome-wide significance not near a GWAS catalog, lead SNP. We present RMPs for two CD associations of genome-wide significance, which were not within 0.1cM of a GWAS cataloged, lead SNP for CD. **a.** The first locus near *TTC33* (Naïve CD4+ T cell context) was reported in the original study as part of a larger locus upstream of the *PTGER4* gene, but was too far in terms of genetic and physical distance to the associated DHS we found with our distance thresholds (Methods). **b.** The second locus near *BRD7* (Activated CD4+ T cell context) cross-replicated in our study with UC, but was not reported in the original study or in the GWAS catalog. Given that this locus reached genome-wide significance and replicated with UC, we suggest that this is a true genome-wide association. The nearest reported locus to it by the authors of the original GWAS was with *NOD2*, but the genomic interval they reported for the *NOD2* association did not overlap with this region (accounting for the usage of hg18 coordinates in the original study).

**Figure S17.**
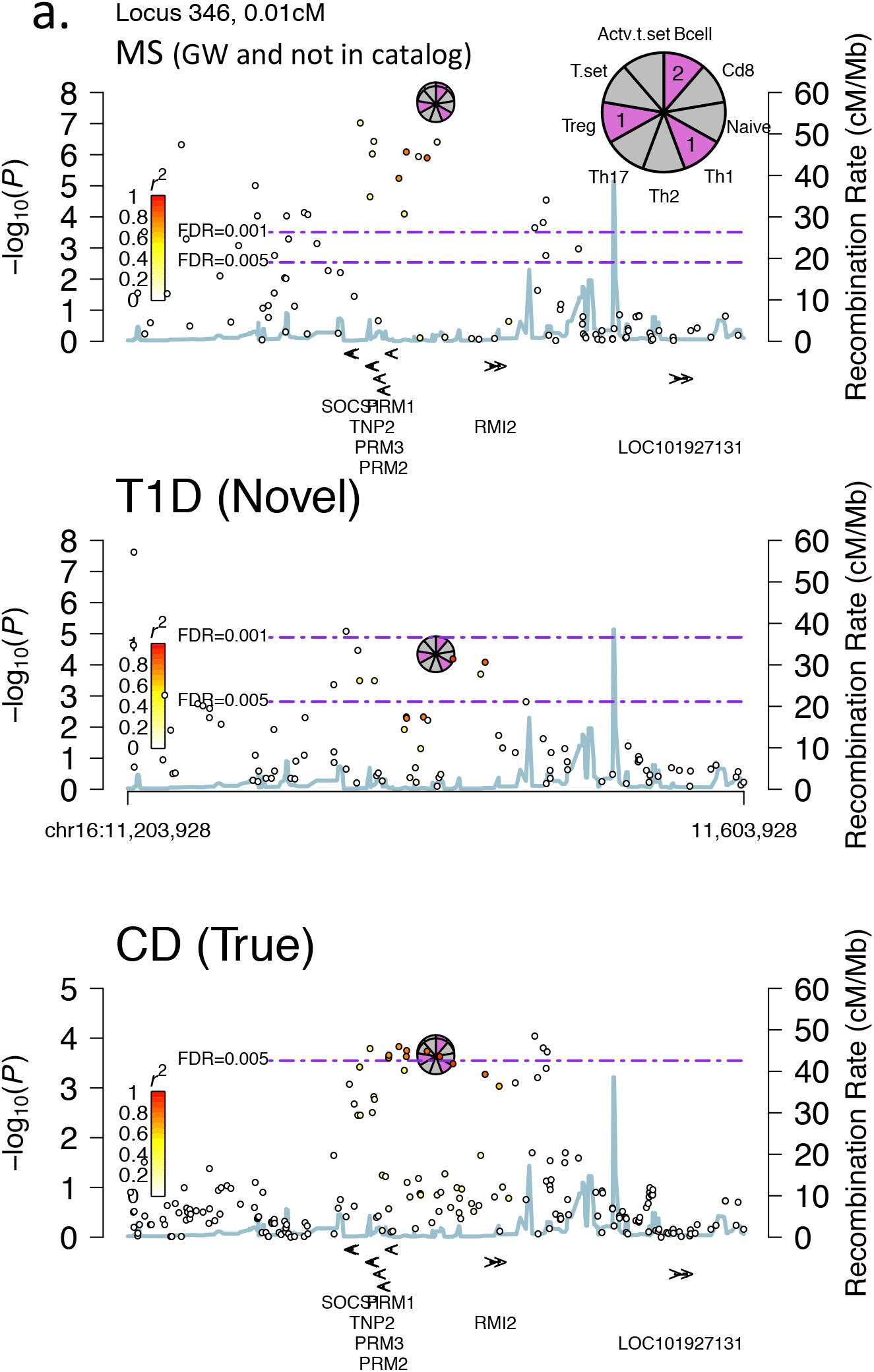
One Multiple sclerosis locus with genome-wide significance not near a GWAS catalog, lead SNP. We present an RMP for one MS association of genome-wide significance, which was not within 0.1cM of a GWAS cataloged, lead SNP for MS. **a.** The locus was near the *SOCS1* gene TSS and was reported in the original study but was not incorporated into the GWAS catalog as far as we could tell.

**Figure S18-S27.**
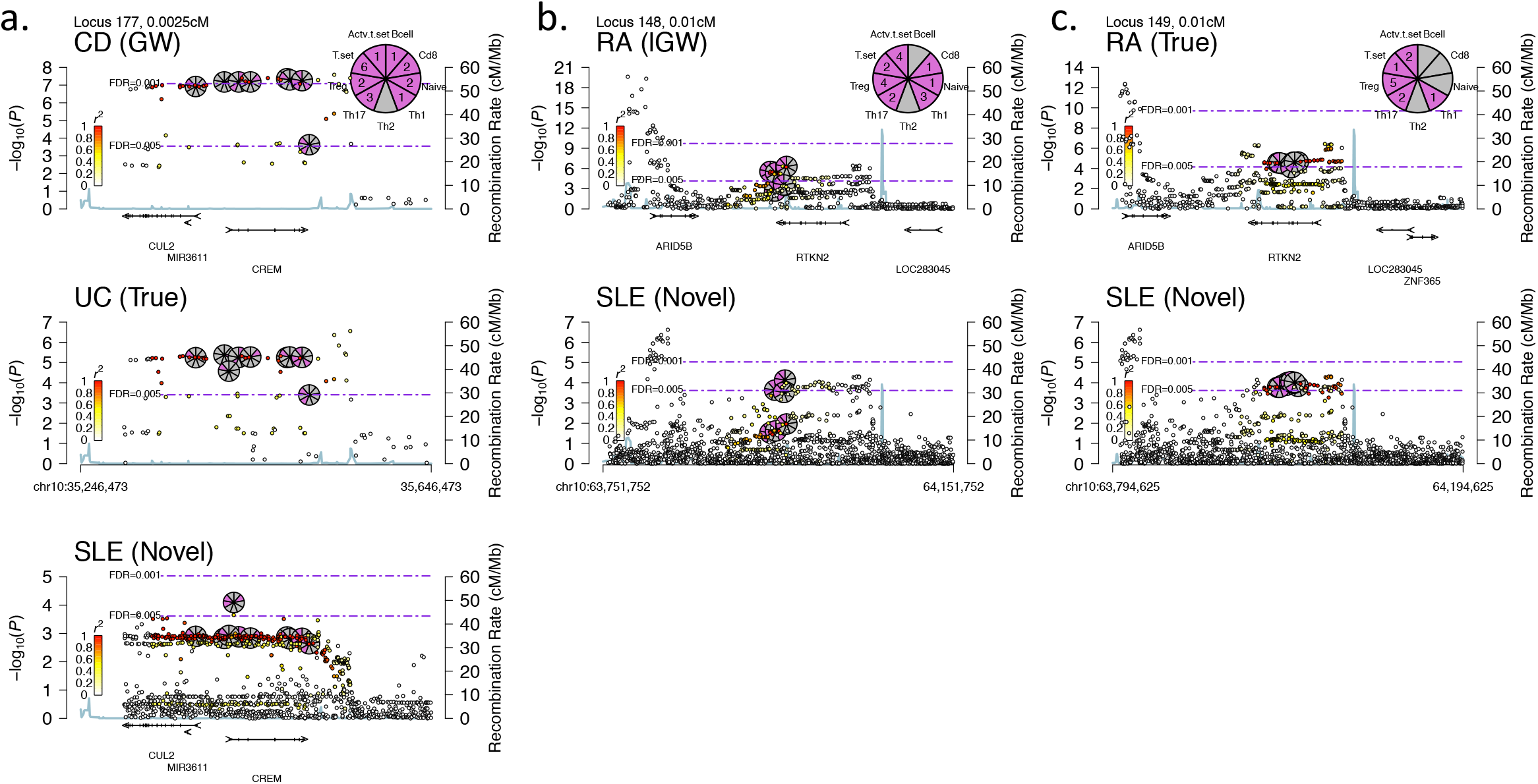
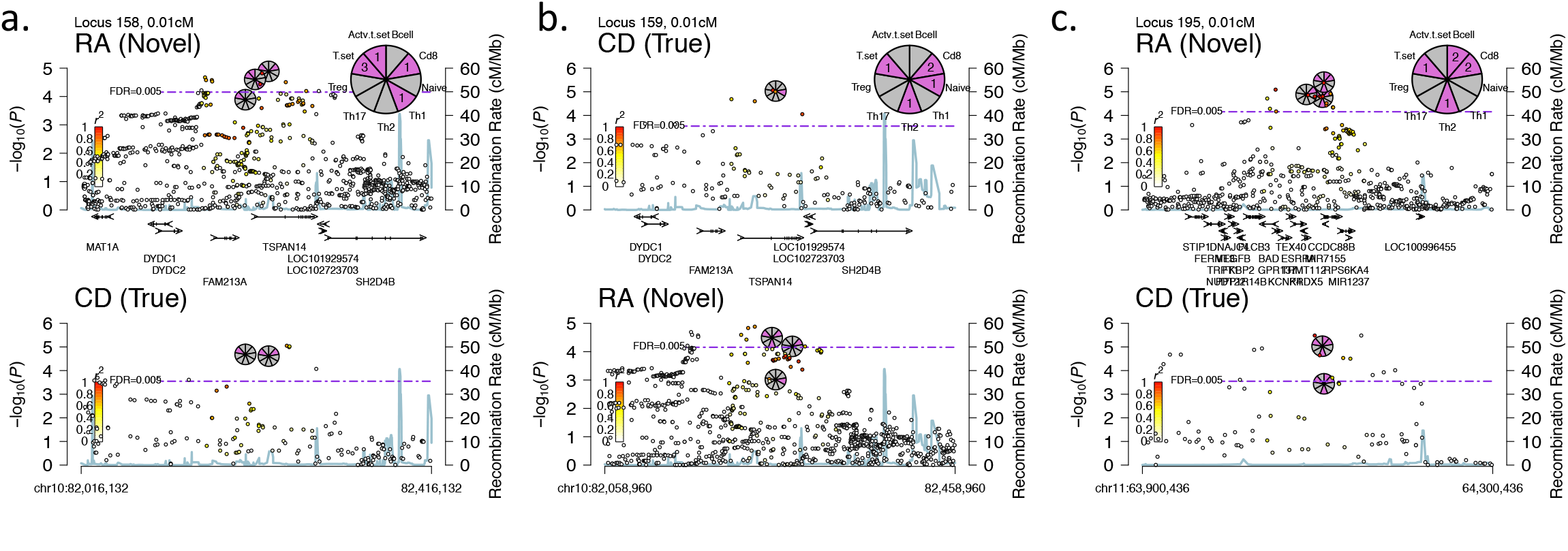
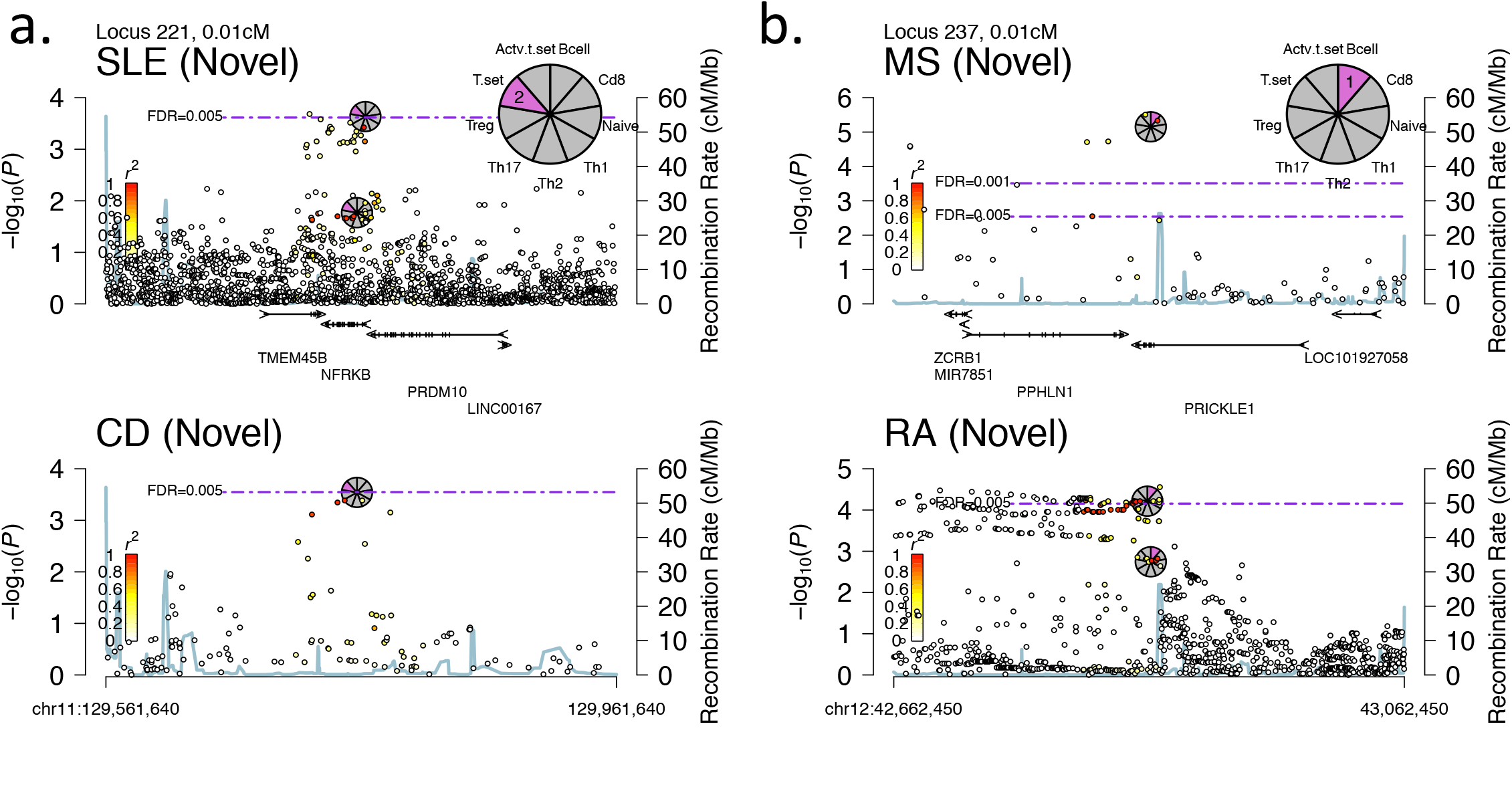
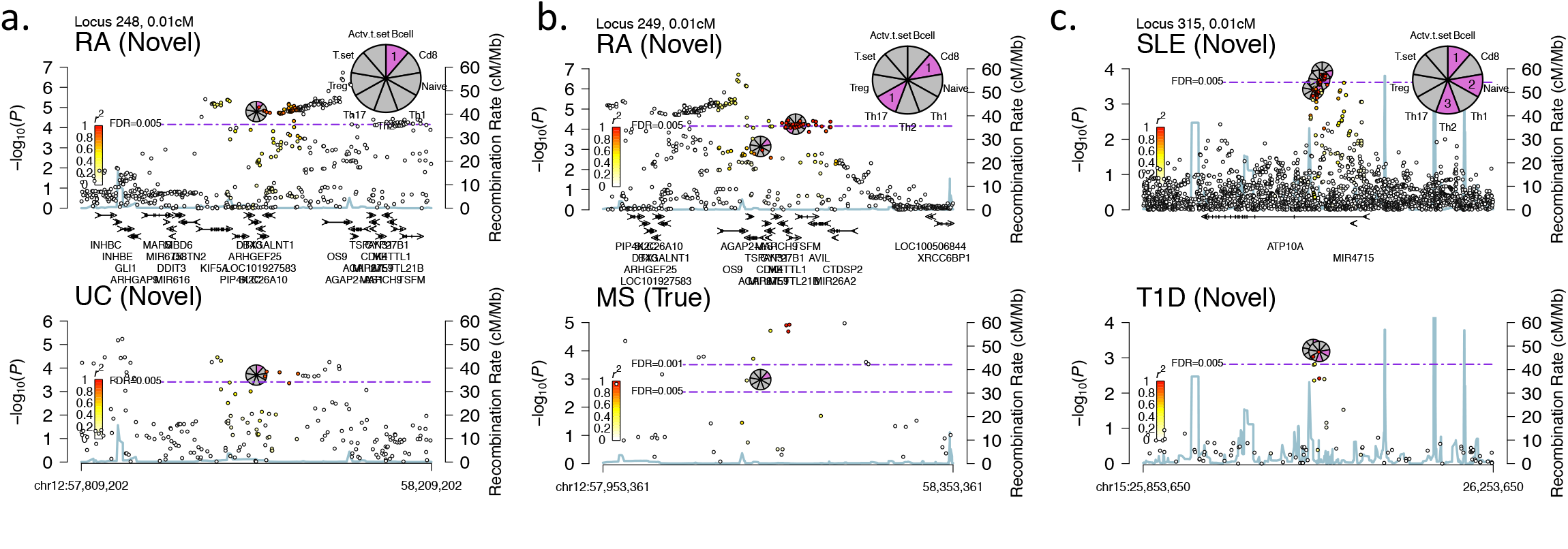
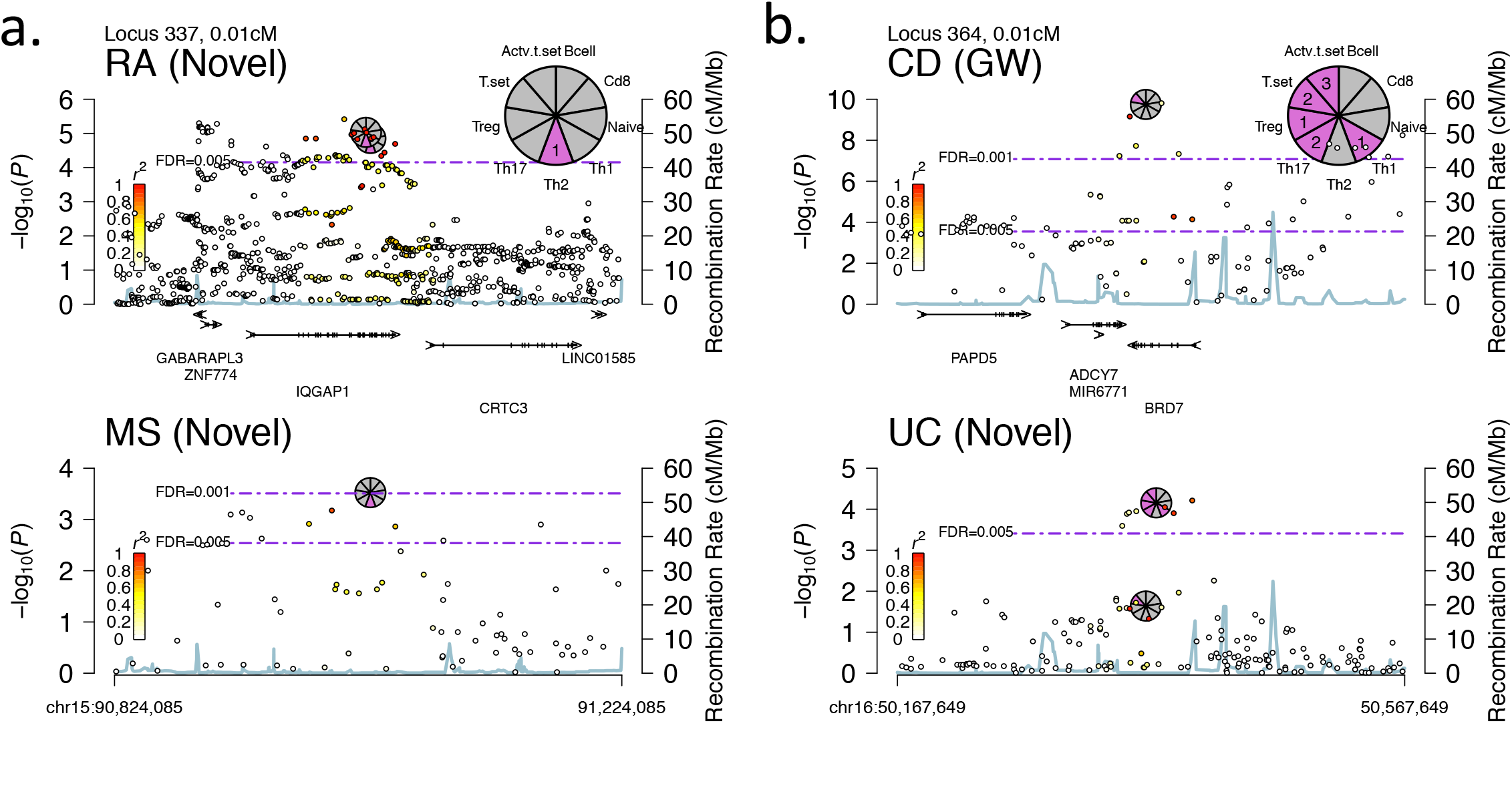
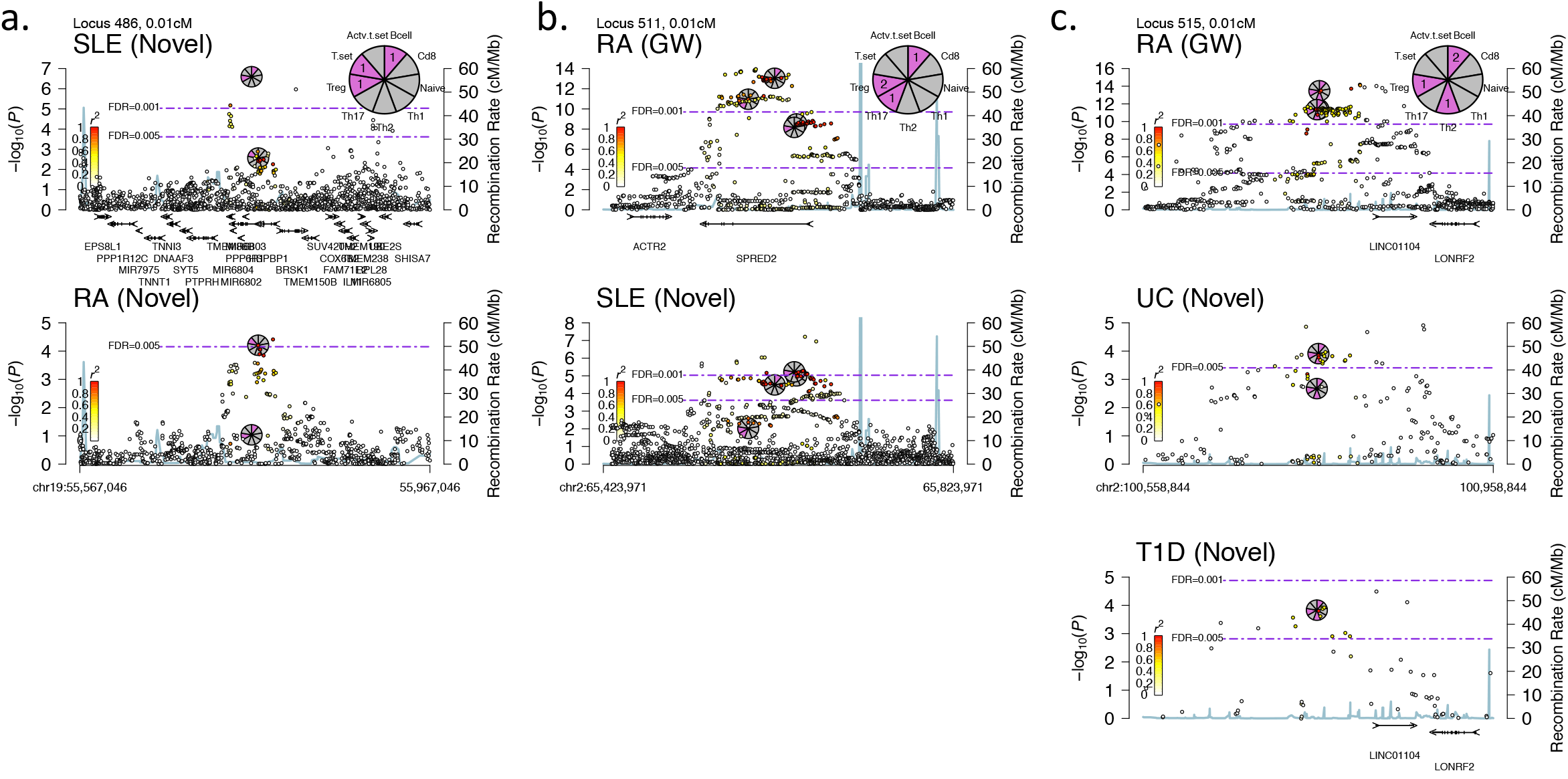
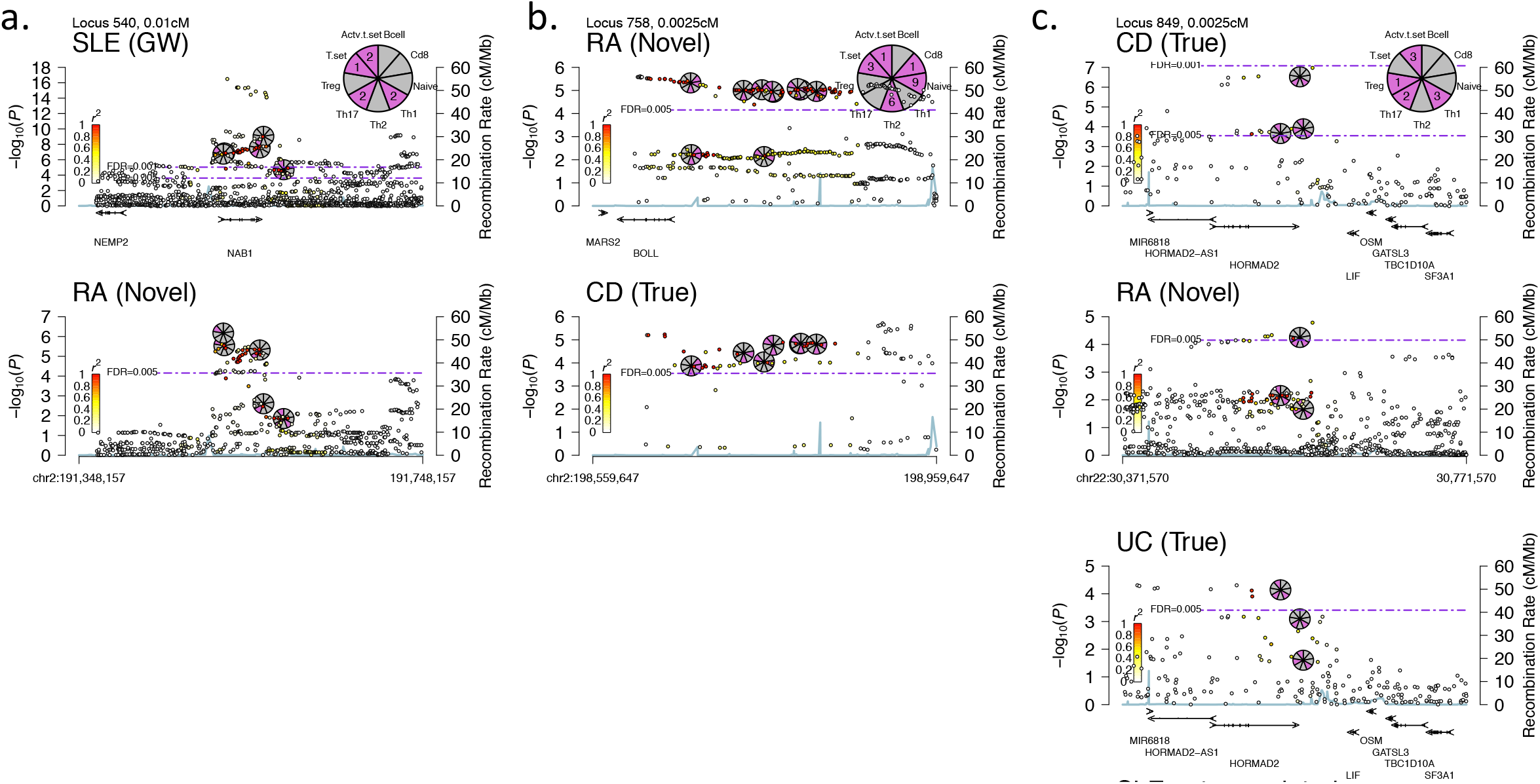
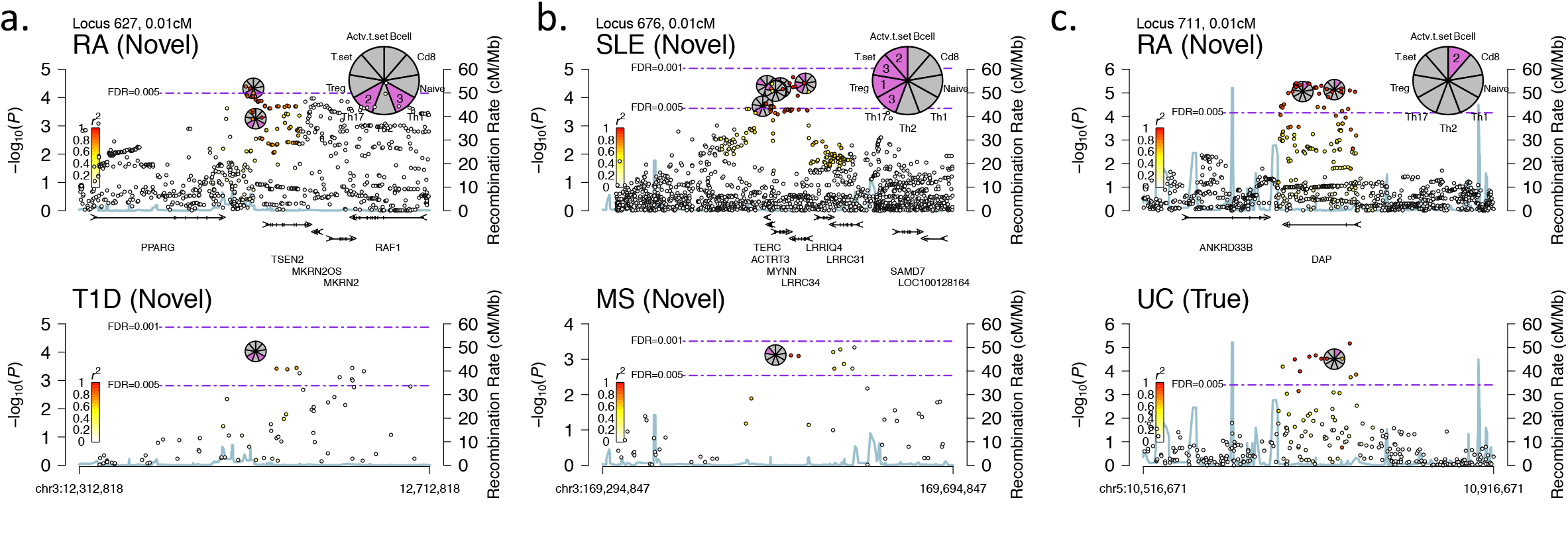
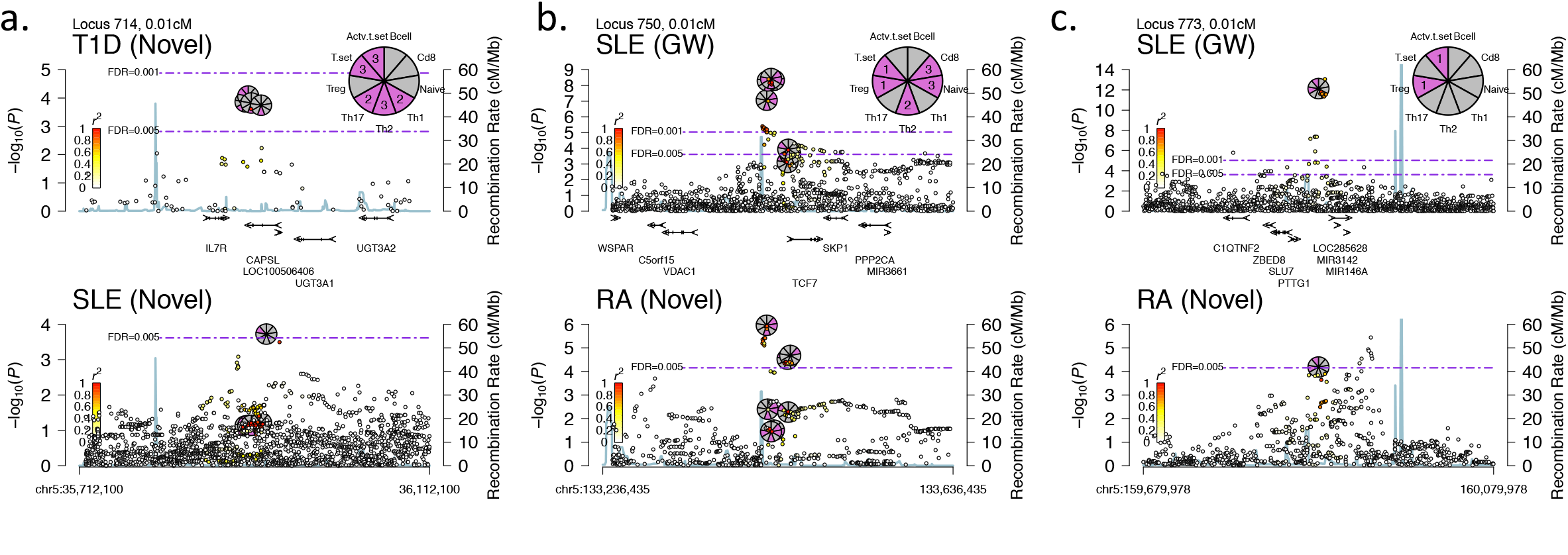
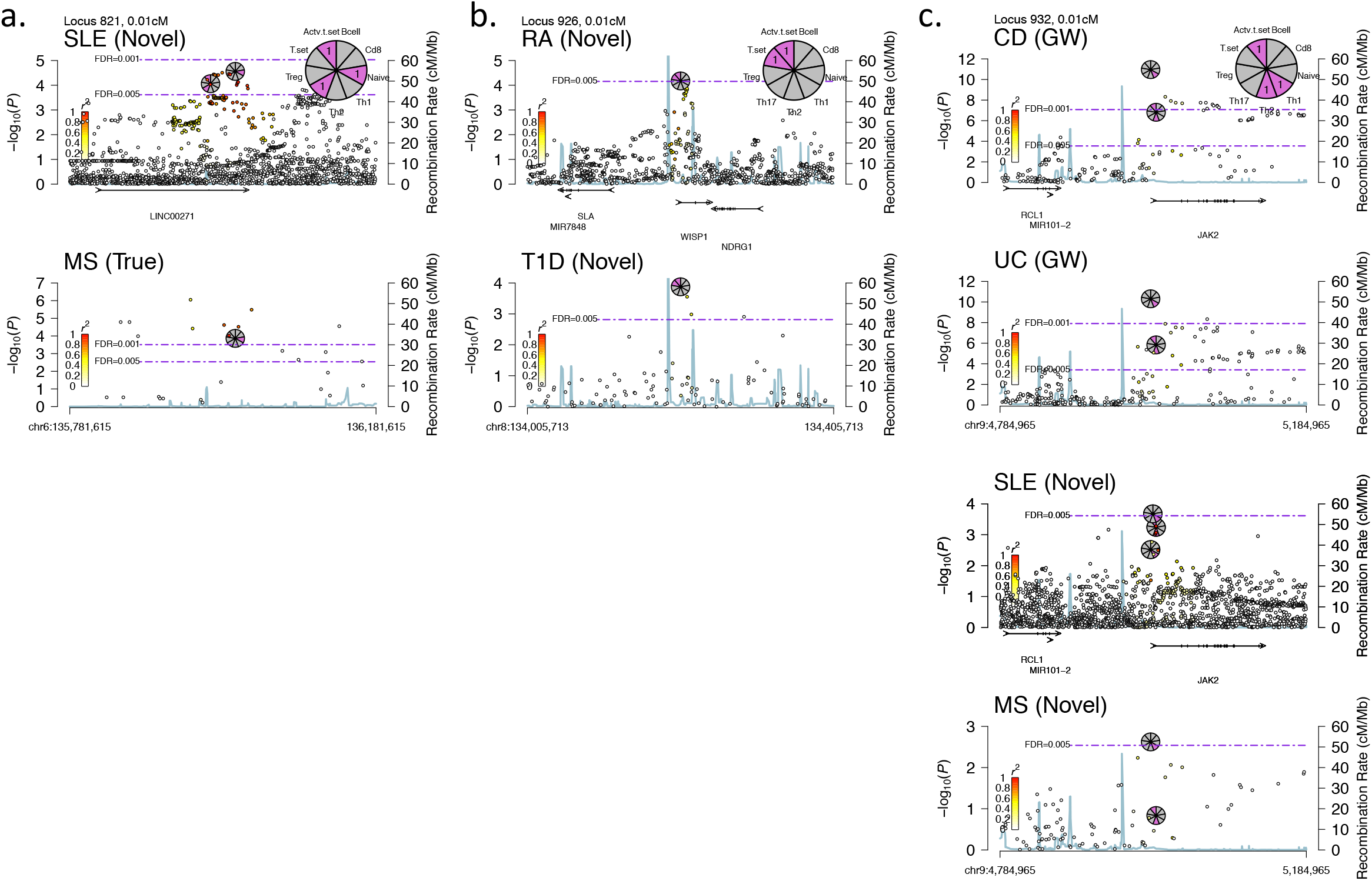
RMPs for novel replicating loci identified in this study. We present RMPs for novel loci found in these study.

**Figure S28.**
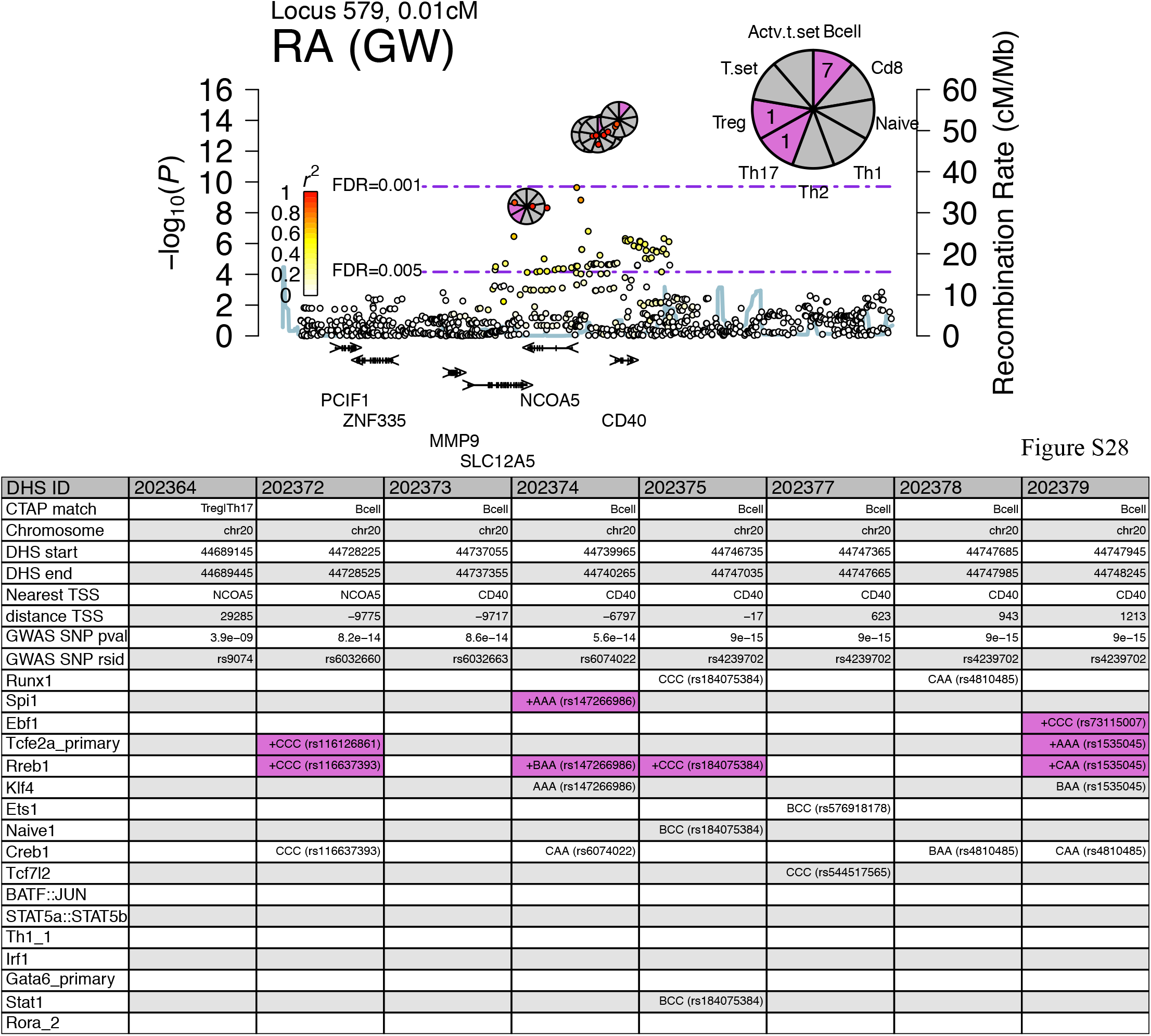
Example RMP with the attached regulatory information table. For each generated RMP we also included an information table giving similar information as that found in the network, but in a tabular format and on a per locus scale. For example, the table here lists all the polymorphic TF binding sites found in the locus with their three letter grades and intersecting SNP rsIDs. This example is for the *CD40* gene region. The table has a column for each associated DHS. In total, there were eight associated DHSs in the region. The first row specifies the accessibility profile memberships assigned to each DHS. In cases were multiple profiles were assigned, profiles were delimited by a vertical bar and listed from left to right by their match ranking (better matches to the left). For example, DHS 202364 (leftmost column) was a better match for Treg than for Th17, although both profiles were assigned to it. Rows two to four show the hg19 physical position of the DHS tag SNP. Rows five to six give the nearest TSS gene name and TSS distance from the DHS midpoint (negative distances indicate upstream positions). Note that if multiple TSSs were annotated in Refseq for a given gene, all such TSS positions were considered in the table; however, the RMP only show one of these (the TSS of the longest isoform by our convention). At times, this may lead to some discrepancies between the table’ reported TSS distance, and what one can see qualitatively from the RMP. Rows seven to eight specify the DHS tag SNP rsID and GWAS *P* value. Following rows specify for each polymorphic binding site, if it was found in a given DHS, and if was found, provide the three-letter grade for the best matching sequence as well as the intersecting SNP rsID. ‘+’ binding sites were further visually highlighted in purple. These are the most probable, functional TF binding sites, given that they were found in a DHS of an accessibility profile the TFBS was enriched in. Taken together, the table allows efficient identification of candidate regulatory SNPs, together with their cellular contexts. In this example, the +AAA SPI1 binding site in DHS 202374 assigned to a B-cell accessibility profile, and the +AAA TCFE2A binding site in DHS 202379 that was also assigned to B cells, are the two top candidates suggested for follow up by our criteria.

**Figure S29-S30.**
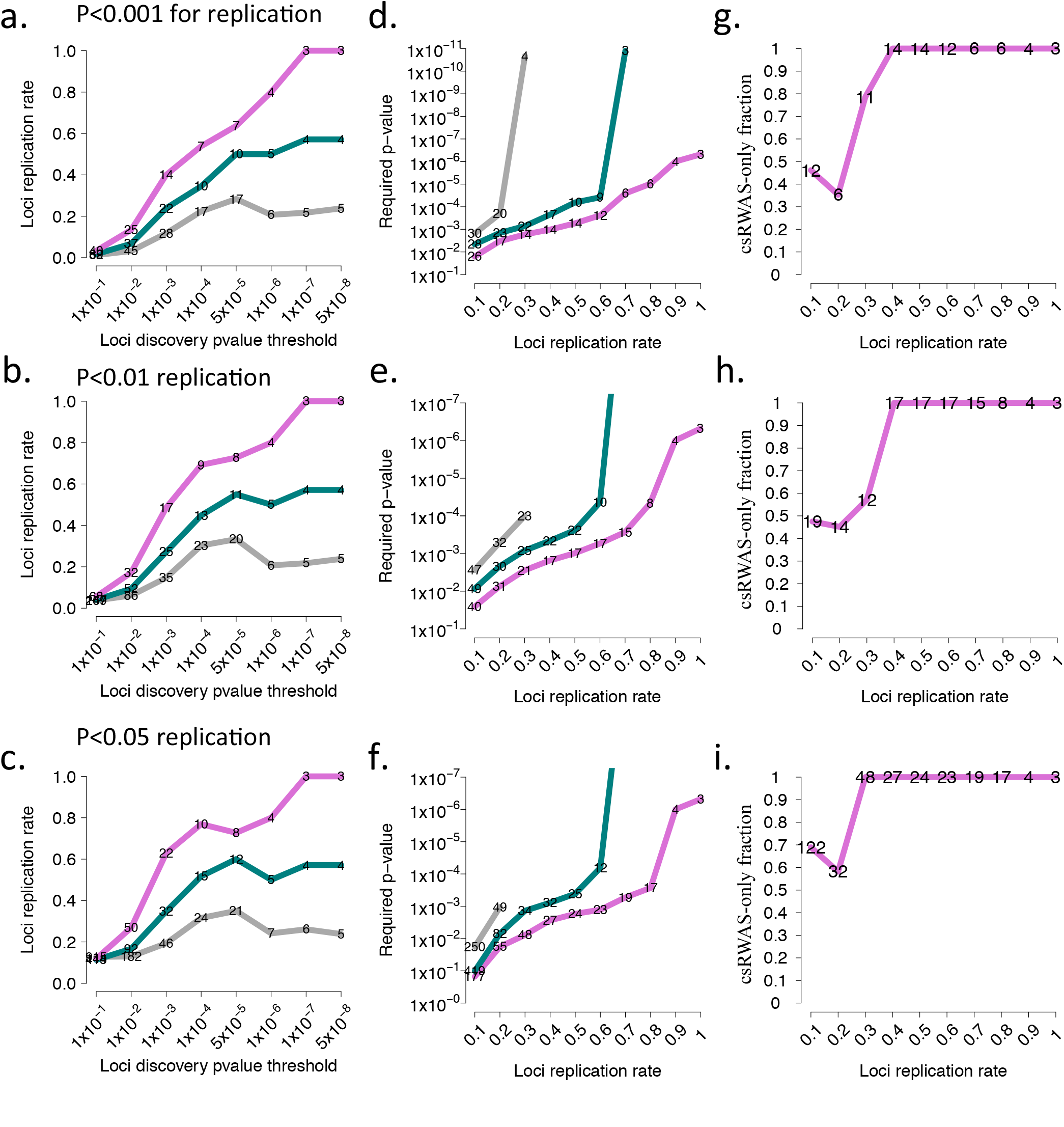
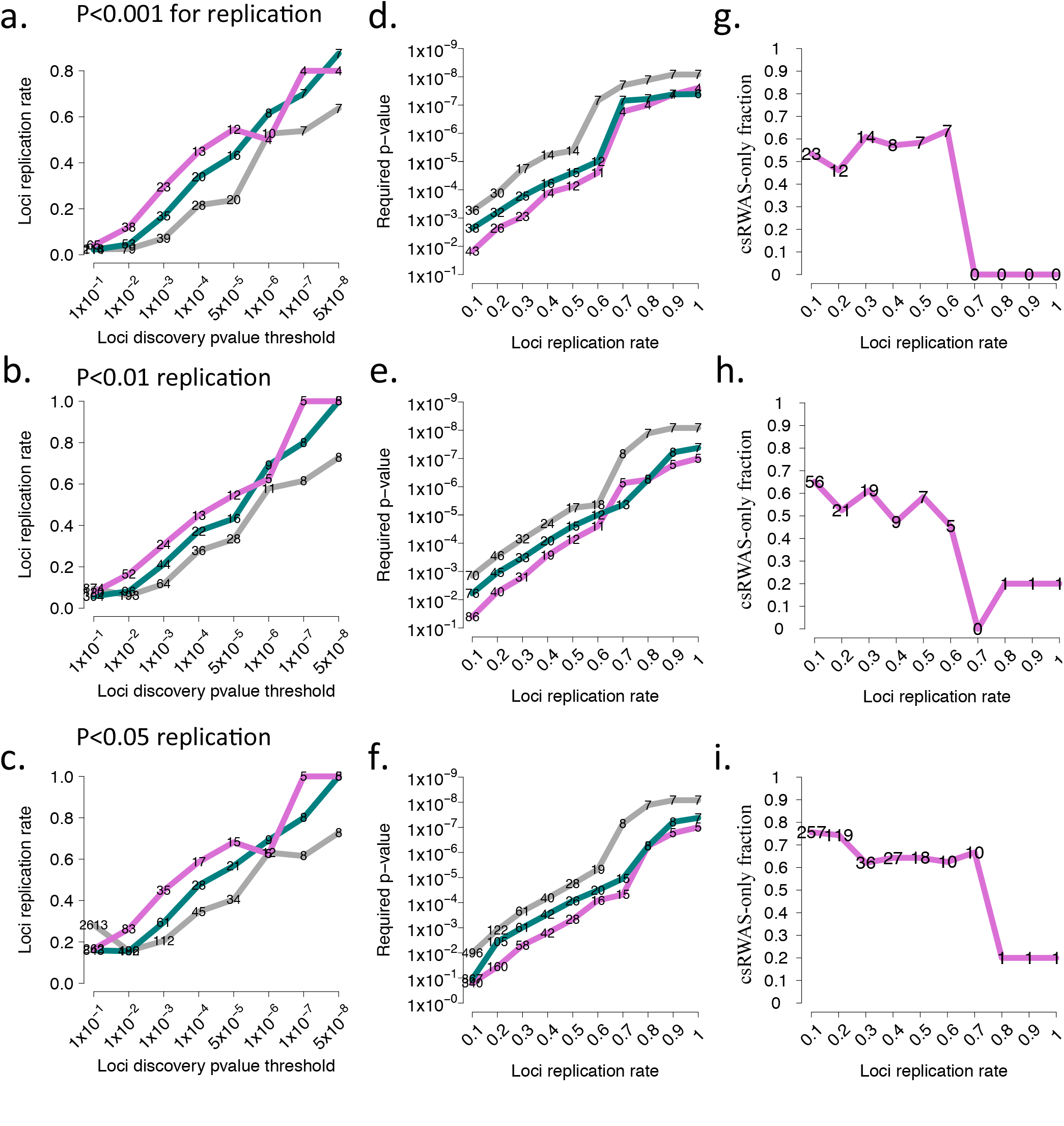
Replication studies comparing csRWAS to GWAS (varying the nominal replication *P* value) We allowed for two additional nominal significance *P* values for loci replication, 0.001 (top three plots), and 0.05 (bottom three plots), in addition to 0.01 used for main results (middle three plots). Figure S29 shows the results for CD, and Figure S30 for RA. The results were consistent across these three *P*-value thresholds (for each column compare the middle plot to the upper and lower plots). For a description of each type of the three plots, see the legend for Figure 7.

## Online methods

### Obtaining and preprocessing of DNase-seq data

DNase-seq data was obtained from ENCODE^16,19^ and the Roadmap Epigenomics project^17^. ENCODE data was downloaded from: http://hgdownload.cse.ucsc.edu/goldenPath/hg19/encodeDCC/wgEncodeUwDgf/. For each cell type, the “NarrowPeak” file (defining the peak regions) and the “Signal” file (providing the number of DNase1 cuts observed for each nucleotide in the genome) were downloaded.

Roadmap Epigenomics project files were downloaded in bam format from GEO^72^, as NarrowPeak and Signal files were not provided. The aligned reads in .bam format were then used as input to hotspot^6^ (the same algorithm used by ENCODE) to generate NarrowPeak files for Roadmap data, and to generate signal files by an in house script. Unlike all the other DNase-seq data, the CD8+ T cell sample from Roadmap was sequenced using 100bps paired-ends (compared to 36bp single end for all other samples). To put the data on the same footing we trimmed the pair-end data to 36 bps using the BBMap suite: https://www.biostarhandbook.com/unit/bbmap-help.html. We also noted that Illumina adapters were common in the reads (likely because of short selected insert size, coupled with the longer reads), so we removed these adapter sequences using BBMap as well before continuing to map paired-reads to the hg19 genome using bowtie2^73^.

In total we obtained DNase-seq data for the following fifteen cell types (in parentheses we provide the short name we used for these samples throughout the manuscript):

1. ENCODE: Embryonic stem cells (hESCH7)
  a. wgEncodeUwDgfH7esPkV2.narrowPeak.gz
  b. wgEncodeUwDgfH7esSig.bigWig
2. Roadmap: Fetal Brain (FetalBrain)
  a. NarrowPeak file generated from .bam using hotspot
  b. Signal file generated using in house script
3. ENCODE: Brain Astrocyte (BrainAstro)
  a. wgEncodeUwDgfNhaPk.narrowPeak.gz
  b. wgEncodeUwDgfNhaSig.bigWig
4. ENCODE: Skeletal Myoblasts (Myoblast)
  a. wgEncodeUwDgfHsmmPk.narrowPeak.gz
  b. wgEncodeUwDgfHsmmSig.bigWig
5. ENCODE: Dermal fibroblasts (Fibrobalst)
  a. wgEncodeUwDgfNhdfadPkV2.narrowPeak.gz
  b. wgEncodeUwDgfNhdfadSig.bigWig
6. ENCODE: Esophageal epithelium (Epithelial)
  a. wgEncodeUwDgfHeePk.narrowPeak.gz
  b. wgEncodeUwDgfHeeSig.bigWig
7. Roadmap: CD34+ hematopoietic Stem cells (Cd34+ HematoSC)
  a. NarrowPeak file generated from .bam using hotspot
  b. Signal file generated using in house script
8. ENCODE: Monocytes (Monocyte)
  a. wgEncodeUwDgfMonocd14ro1746Pk.narrowPeak.gz
  b. wgEncodeUwDgfMonocd14ro1746Sig.bigWig
9. ENCODE: B cells (B cell)
  a. wgEncodeUwDgfCd20ro01778Pk.narrowPeak.gz
  b. wgEncodeUwDgfCd20ro01778Sig.bigWig
10. Roadmap: CD8+ T cells (Cd8+)
  a. NarrowPeak file generated from .bam using hotspot
  b. Signal file generated using in house script
11. ENCODE: Naive CD4+ T cells (Naive CD4+ T cell)
  a. wgEncodeUwDgfCd4naivewb11970640Pk.narrowPeak.gz
  b. wgEncodeUwDgfCd4naivewb11970640Sig.bigWig
12. ENCODE: Th1 T cells (Th1)
  a. wgEncodeUwDgfTh1wb33676984Pk.narrowPeak.gz
  b. wgEncodeUwDgfTh1wb33676984Pk.Sig.bigWig
13. ENCODE: Th2 T cells (Th2)
  a. wgEncodeUwDgfTh2Pk.narrowPeak.gz
  b. wgEncodeUwDgfTh2Sig.bigWig
14. ENCODE: T-regulatory (Treg)
  a. wgEncodeUwDgfTregwb78495824Pk.narrowPeak.gz
  b. wgEncodeUwDgfTregwb78495824Sig.bigWig
15. ENCODE: Th17 (Th17)
  a. wgEncodeUwDgfTh17Pk.narrowPeak.gz
  b. wgEncodeUwDgfTh17Sig.bigWig

### Collapsing DHS annotations from multiple files

Let *N* be the number of DNase-seq peak files, and denote all the peaks with significance score above a set threshold (here set to 300), by *H*=*[h^1^, h^2^,…, h^N^]*, where the significance scores were as reported by the hotspot peak detection algorithm^6^. Also, let each peak have the following information associated with it: chromosome, start position, end position, and a score. We defined a DHS locus as the most extreme 5’ and 3’ positions of any overlapping DHS peaks in the set H. We further refined the start and end positions of each locus using the less significant peaks as described next. Taking all peaks not considered so far, i.e. peaks with a significance score below 300, we allowed an extension of a DHS start and end positions, if any of the additional peaks overlapped with one of the previously defined DHSs, and had a more extreme 5’ or 3’ bp positions. This second step does not change the number of DHSs, which was determined by the more significant peaks, but allows for a refinement of loci start and end position to include all relevant sequences for the considered cell types. The result was an annotation of DHS coordinates across the genome, guarantying that each identified DHS was of a high-significance in at least one of the examined cell types.

### Cleavages per kilo base per million (CPKM) as a continuous measure of accessibility for DHSs

DNase-seq measures chromatin accessibility by quantifying how many times each bp position was cleaved and therefore sequenced as the 5’ nucleotide of a read. Equipped with the DHS annotation defined above, one natural measure to quantify accessibility across samples was to calculate the number of cleavages per kilobase per million (CPKM) for each DHS and for each cell type. A small pseudo CPKM value, equal to the 0.05 percentile of all raw CPKMs, was added to each CPKM value to avoid zeros or very small CPKMs. The result was a full, continuous matrix of DHSs by cell types, with each entry holding the accessibility value of a given DHS in a given cell type.

### Matching DHSs to cell-type accessibility profiles

We quantified how well each DHS matched to each of a set of predefined accessibility profiles. There were two main types of accessibility profiles: single cell-type profiles, e.g. to identify Th17-specific DHSs, and high-order profiles, e.g. to identify DHSs accessible in all activated CD4+ T cells (Th1, Th2, Treg, and Th17).

#### Single cell-type specific accessibility profiles

These scores were used to match DHSs to the cell type they were most specific to. Let *x_ij_*, denote the CPKM value of DHS *i* in cell-type *j*. The specificity score of DHS *i* to cell-type *j, s_ij_*, was then calculated as:

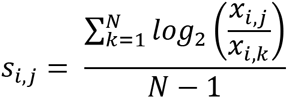

where *N* is the total number of cell types. That is, the average of the log2 fold-change of a given DHS accessibility in a given cell type, compared to the accessibility of the same DHS in all other cell types. *s_i,j_=1* for example would indicate that, on average, DHS *i* is two times more accessible in cell type *j* that in the other cell types, and a value of 2 would indicate that DHS *i* is four times more accessible in cell type *j* than in the other cell types (note the log2 space). As a practical matter, since T-cells had very similar accessibility patterns to one another (as expected for closely related cells) and we wanted to find the differences between T-lineages, when calculating the specificity scores for each T lineage, we first averaged out the CPKM values of all non-T, differentiated cell types (i.e. excluding hESC and fetal brain cell types), into one meta-sample of CPKM value. We then calculated specificity scores as described above. The result was a higher weight given to differences between T-cell lineages over differences between T-cells and other differentiated cell types.

#### High-order accessibility profiles

These scores were used to match DHSs to a subset of cell types they were most specific to. For this we used a correlation approach. Let *h* be an ordering of the fifteen cell types, *h*=(*hESC, Fetal Brain,…, Th17*), and *X* be the CPKM matrix of size *m x n*, where *n* is the number of cell types and *m* is the number of DHSs (columns ordered as in *h*). Also, let *y* be a “fishing” vector with one entry per cell-type (also ordered as in *h*). Then, to determine the match score of DHS *i* with a high-order profile, we assigned *y_j_=1*, if cell-type *j* belonged to that accessibility profile, and *y_j_=0*, otherwise. For example, if the profile of interest was the activated CD4+ T set, and positions 12-15 corresponded to the activated T-cell types: Th1, Th2, Treg, and Th17, then y would equal (0,0,0,0,0,0,0,0,0,0,0,1,1,1,1). We then calculated the correlation between *y* and each DHS (row) in *X*. These correlations were used to decide which DHS belonged to a high-order profile based on either a rank cutoff (controlling for the number of DHSs per profile), or a correlation cutoff (controlling for the minimum correlation of DHSs per profile).

#### Ubiquitous accessibility profile

This score was used to identify ubiquitous DHSs that were robustly accessible across all cell types. The correlation approach cannot work in this case, as the standard deviation of the vector is zero (a vector of 1’s), and the correlation undefined. For this we first narrowed down to DHSs that had a peak in all examined cell-types (from all called DHSs, not limiting to those with score > 300). This resulted in a set of 21,178 DHSs. We prioritized among these DHSs (e.g. to choose the top 600 most ubiquitous DHSs as was done for the DNA motif enrichment analysis) by ranking based on two properties: DHS average CPKM across cell types (from high to low), and the DHS standard deviation across cell types (from low to high), resulting in two ranked lists of length *L*=21,178. DHSs that ranked high in both lists were those found to have similar high accessibility across all cell types. We than rank-combined the two lists by taking their rank-sum for DHS *i* as computed by: *r_i_*=(*r_i,1_/L*+*r_i,2_*/*L*)/*2*, where *r_i,1_* is the DHS rank in list 1 and *r_i,2_* is the DHS rank in list 2. Using these rank-combined scores we ranked among ubiquitous DHSs to prioritize the most ubiquitous DHSs.

### Removing HLA region

We removed the HLA region of the human genome (chr:6:25,000,000-35,000,000; hg19 build) from consideration in all of our analyses. This was done to avoid allowing this highly autoimmunity relevant DNA region from dominating our genome-wide results. Examining the accessibility profiles of DHSs in this region can be the focus of future works.

### Short sequence motif enrichment in DHSs

We identified TF binding sites in DHSs of a given accessibility profile using MEME-ChlP command line tool version 4.1.0^74^. Specifically we used the MEME^75^ and DREME^76^ de novo motif discovery algorithms that were implemented as part of MEME-ChlP. As input we used DNA sequences (100bps centered on DHS midpoints) of the top 600 DHSs matching each profile. All motifs of length greater than 6bps, with an *E*-value<10^-10^, and that were present (one or more occurrences) in at least 60 sequences (10% of input sequences) were considered as enriched. Enriched sequence motifs were matched to known TF binding sites using Tomtom^77^, against two comprehensive databases of known binding sites: Jaspar-2014^78^ and UniPROBE update 2015^79^. Both databases were downloaded from the MEME-CHIP website: http://meme-suite.org/doc/download.html?man_type=web on January 5^th^ 2016. In cases were two or more similar motifs were found for a given accessibility profile (e.g. if both MEME and DREME found the same motif), the motif with the higher significance was kept. This step was handled automatically by MEME-CHIP. MEME motifs with a Tomtom match score <0.01 were given the matched TF name. DREME motifs with a Tomtom match score <0.1 were given the matched TF name. For DREME we used a more lenient cutoff because DREME motifs were guaranteed to be shorter than 8bps by the algorithm’s design and this led to many partial matches of lower TOMTOM scores to known TF binding sites, when compared to the longer (on average) MEME motifs. Motifs not matched to a known TF binding site were considered novel and annotated using the nomenclature: [profile name]_[count]. These novel motifs were considered in the same manner as known TF binding sites for subsequent analyses (i.e., we did not limit subsequent analyses to known TF binding sites). Lastly, promoter regions (within 2kb of an annotated TSS) were not considered to avoid identifying core promoter elements and to instead favor TF binding sites.

Command line used: meme-chip-noecho-oc $PATH_OUT$PATH_OUT_SUFFIX - index-name $PATH_OUT_SUFFIX.html-db $FILE_MOTIFS_JASPAR-db$FILE_MOTIFS_UNIPROBE-meme-mod zoops-meme-minw 6 -meme-maxw 18- meme-nmotifs 6 --meme-minsites 60 -dreme-e 0.05 -dreme-m 10-centrimo-local - centrimo-score 5 -centrimo-ethresh 10 $PATH_SEQ$SEQ_FILE. Where “$” variables indicate directory and file locations on our computer.

### GO enrichment analysis

We associated DHSs to the nearest gene transcript-start-site (TSS) as a proxy for likely regulated genes, and tested the top ranking 600 unique genes per profile for GO function enrichment using an implementation of the hypergeometric test in the FUNC software package ^29^. We considered all enriched GO terms per profile at FDR<0.05 and *P* value<0.001. We post-processed the enriched terms found, using the refinement algorithm in FUNC that filters out higher level GO categories, which were not enriched after removing genes from lower level GO enriched categories. We only considered enriched GO categories supported by more than 5 genes. From the remaining GO terms, we reported the top six terms per profile.

### Gene expression analysis

Tissue and cell type expression data was obtained from two sources that were based on two different technologies:

1. Array based expression data: the Human GNF1H tissue-specific expression dataset^80^ downloaded via the BioGPS website (http://biogps.org/), with expression data available for 13,066 genes across 84 tissues/cell types.
2. Sequencing based expression data: the GTEx expression database^31^, V6 (dbGaP Accession phs000424.v6.p1), with expression data available for 54,148 genes and noncoding RNAs across 53 tissues and cell types (after ignoring genes with the same symbol name associated to more than one loci in the genome, to remove ambiguity). In the GTEx database we averaged for each gene in each tissue or cell type its RPKM values across all individuals to obtain one RPKM value per tissue or cell type. To allow comparison between genes, the expression of each gene was normalized across all tissues and cell types in each dataset (BioGPS/GTEx) to obtain tissue or cell type specific Z-score expression values.

For BioGPS, we used the expression data across the following six tissues or cell types: bronchial epithelial cells, CD19+ B cells, CD4+ T cells, skeletal muscle, CD14+ monocytes (GNF1H dataset) and whole brain. For GTEx, we used the expression data across the following six tissues or cell types: Esophagus mucosa, whole blood, EBV transformed lymphocytes, skeletal muscle, brain cortex, and brain amygdala. These tissues and cell types where chosen because they were related to one or more of the accessible cell types in the profiles we assayed. For both datasets we assigned genes to DHSs by the rule of nearest TSS. DHSs assigned to genes with no expression data were ignored. For each accessibility profile we extracted the top 600 unique genes and plotted their *Z* score boxplot, in each tissue or cell type, comparing them side-by-side.

### Alternative assignment of DHSs to cell-type accessibility profiles based on binary accessibility values

We compared two approaches for grouping DHSs into accessibility profiles: using CPKM values as described above, or using binary assignment of DHSs to cell-types (the method used by ENCODE for example). In the latter case, non-overlapping DHSs found in only one cell type were considered as cell-type specific, and similarly, DHSs found only in a subset of cell-types comprising a high-order profile, were assigned to that accessibility profile. Note that the CPKM approach is fuzzy in that a DHS were assigned a match score to any profile, and thereby not a priori limiting the number of identified DHSs per profile. In comparison, the binary approach was not fuzzy in that each DHS can belong to only one profile. Note also, that in both cases the same single annotation of DHSs was used, i.e. the only difference is in what values the entries in the accessibility matrix can take.

### Obtaining GWAS data

We obtained genotype data from the Wellcome Trust Case Control Consortium (WTCCC) for the following diseases: type 1 diabetes (T1D)^41^, type 2 diabetes (T2D)^41^, Crohn’s disease (CD)^41^, ulcerative colitis (UC)^37^, and multiple sclerosis (MS)^39^. Data for additional GWAS was obtained as summary statistics from the: Rheumatoid Arthritis Consortium International for Immunochip (RACI)^36^, International Inflammatory Bowel Disease Genetics Consortium (IIBDGC)^38,81^, Coronary Artery Disease Genetics Consortium (C4D)^82^, International Consortium for Blood Pressure (ICBP)^83^, DIAbetes Genetics Replication And Meta-analysis (DIAGRAM)^84^, Genetic Investigation of ANthropometric Traits (GIANT)^85,86^, International Genomics of Alzheimer’s Project (IGAP)^87^, Psychiatric Genomics Consortium (PGC)^87,88^, and the Euro-Canadian systemic lupus erythematosus GWAS (EC-SLE)^40^. All GWAS used with details about population, number of cases, number of controls, SNP number (tag or imputed), primary publication pubmed ID (PMID), if the trait was considered as autoimmune by us, and the consortium that reported the data, can be found in the readme file for table S2.

### GWAS processing

We analyzed the WTCCC datasets starting with the primary genotype data. We used the plink whole-genome association tool set^89^. We removed individuals inferred to be related (plink generated pi-hat larger than 0.12), individuals with more than 10% missing genotypes, and individuals with a reported sex not matching the expected X to autosome ratio. We filtered SNPs with >10% missingness, with a minor allele frequency (MAF) < 0.005, and for which missingness was significantly correlated with phenotype (P<0.05 after Bonferroni correction for the number of SNPs in the study). We ran principal component analysis (PCA) using EIGENSOFT^90^ after pruning for linkage disequilibrium (LD) and removing large LD blocks^91^. Individuals inferred to be of non-European ancestry by EIGENSOFT were removed from all subsequent analysis (five iterations of outlier removal, where outliers were determined by exceeding six standard deviations along one of the first ten PC’s.). SNP *P* values were determined using Plink’s logistic regression option with the top ten PC’s provided as covariates to help control for possible hidden population structure among cases and controls. Finally, we estimated genomic inflation factors for each GWAS using the R package GenABEL^92^, estlambda(method="median"), i.e. 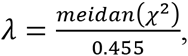 using all TSS-distal SNPs as input (greater than 30kb away of any TSS), as was previously suggested^23^. We corrected all studies for inflation using the above lambdas, including studies for which we directly downloaded summary statistics.

### Enrichment of GWAS associations with DHSs stratified by cell type accessibility profiles

Each DHS was assigned the GWAS SNP with the minimum *P*-value in an interval of 2500 bps surrounding the DHS midpoint. For each GWAS, DHSs with no such SNP were discarded. We ranked the top 5,000 remaining DHSs by their assigned SNP *P*-values, from low to high, and calculated a cumulative enrichment score for each profile, defined as the proportion of DHSs with a match score greater than the 95^th^ percentile of all match scores for that accessibility profile. More formally, assume without loss of generality that there is only one GWAS and accessibility profile, and denote by *c* the 95% percentile-score of all DHS match scores for that accessibility profile (i.e. 5% of DHSs will have a score larger than c). We then add one DHS into consideration at a time (where DHSs were ordered by their significance) and calculate the proportion of DHSs with a match score larger than *c*, as each DHS is added into consideration. The expectation is that this cumulative proportion will fluctuate around 5%, with larger deviation in the beginning, as only a small number of DHSs contributed to the proportion up to that point. The standard deviation with each additional DHS was estimated using a binomial sampling process with *n* as the number of DHSs seen to that point (where *n=1, 2,…, 5000)*, and *p=0.05* is the constant probability of success. The enrichment curve was then simply defined as this cumulative proportion line (see examples in Fig S9a). The significance of this enrichment curve was calculated as the area under the curve (AUC), where the maximal possible AUC was normalized to 1. The expectation is therefore an AUC of 0.05. We estimated the standard deviation of this expectation, using 10,000 simulations, with 5000 DHSs ordered at random (not by significance). The null distribution of the summary statistics and the observed summary statistics for each profile and GWAS pair were shown in Fig. S9b. Since multiple enrichments were considered, we applied a Bonferroni correction cutoff (*p*<0.01 after correction) to determine significant enrichments, and visualized these significant profile and GWAS pairs in a heatmap, as seen in Fig. S10a.

### A context specific regulatory-wide association study

We tested DHSs specific to adaptive-immune cell types for genetic associations with GWAS of autoimmune diseases. Each DHS was assigned the GWAS SNP with the minimum *P* value in an interval of 2500 bps of the DHS midpoint. For each GWAS, DHSs with no such SNP were. The total pool of DHSs for each study was then selected as follows: the union of the top 5,000 DHSs matching each of the following cell-type-specific activity profiles (B cell, CD8+ T, Naïve CD4+ T, Th1, Th2, Th17 and Treg), and all DHSs with a correlation match-score > 0.7 with the T set or Activated CD4+ T set. Note that this procedure can match a DHS to more than one activity profile. Out of the 348,527 DHSs, this procedure selected at most 62,586 unique DHSs. This number was smaller for non-imputed GWAS data, as many DHSs would not be tagged by a SNP within the 2500 bps interval. Assuming a human genome size of 3x10^9^, and a DHS size of 300 bps (which is enough to capture the signal observed for most DHSs, see Fig. 2b and Fig. S2), the total number of base pairs covered by all 348,527 DHSs was approximately 105 million (3.5% of genome), and the total covered by adaptive-immune DHSs was 11 million (0.37% of the genome). To determine the significance of DHS-associations we used an empirical accessibility-based FDR with a cutoff of 0.005. For each study, *P* value thresholds corresponding to a given FDR were determined from a null distribution of *P* values comprised of all non-immune DHS *P* values. The pool of non-immune DHSs was determined as follows: we ordered DHSs nine times by their profile match scores to each of the nine adaptive-immune-specific accessibility profiles (i.e. B cell specific, CD8+ T-cell specific,…, T set specific, and activated CD4+ T set), generating nine ranked lists of DHSs. We then rank-combined these nine lists into one ranked list, v, using the equation (per DHS 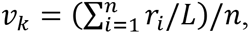 where *n* is the number of adaptive-immune accessibility profiles, *r_i_* is the rank of DHS *k* in profile *i=1,…n*, and *L* is the total number of DHSs. The bottom-ranked entries in *v* represent the DHSs that were least accessible in the adaptive-immune cell types. Using this property, we selected the DHSs ranking in the bottom half of *v*, as the set of non-immune DHSs, and used their *P* values for the null distribution. As before, for each GWAS, DHSs with no tag SNP within 2.5 kb were discarded. The corresponding *P* value, p, for a given FDR threshold *f* was then determined as the threshold, *p*, for which a proportion of *ƒP*-values was smaller than *p*. That is, for FDR<0.005 this procedure will find the *P* value cutoff where a proportion of 0.005 of the non-immune DHSs have equal or smaller *P* values that that (i.e., an estimate of the proportion of DHSs expected by chance at this *P* value threshold).

We grouped associated DHSs into loci in two manners. The first was when considering each study independently. The second was when considering DHSs associated to any of the six autoimmune diseases, as if they came from a single GWAS study. The latter strategy helped in determining which loci were shared across autoimmune diseases. We determined SNP loci memberships using a genetic distance cutoff of 0.1cM or 0.01cM, if the studies were considered individually, and 0.01cM and 0.0025cM, if all studies were considered as one. We focused on results from all GWAS together, with a 0.01cM cutoff, to leverage the related autoimmune diseases for replication, and to help ensure that replicating loci were more likely due to shared associated DHSs than some more general relevance of a larger region around the DHSs for the disease. The HapMap Phase II data^93^ was used to assign genetic position in cM to each GWAS SNP. A DHS (or GWAS SNPs not in the HapMap data) were assigned a genetic position using linear interpolation between the nearest flanking HapMap SNPs. If a DHS did not have a right or left flanking SNP (may happen at chromosomes ends) we used linear extrapolation from the pair of rightmost or leftmost HapMap SNPs, for the right and left sides of the chromosome, respectively. DHSs or SNPs were then grouped into loci using the following procedure:

#### Pseudo-code

Starting with chromosome one, assign the leftmost associated DHS (most 5’) to current_locus=1, and repeat the following steps:

If the next leftmost DHS is within genetic distance *d* from the previously assigned DHS, add it to the current_locus.

Otherwise assign the DHS to current_locus=current_locus+1

Repeat until all DHSs in the chromosome have been assigned to a locus. Repeat this procedure for each chromosome.

### Classifying DHS associations

We used several groupings to inform about the reliability of each association and to define which associations are novel findings to this work. The first grouping was based on the commonly used genome-wide significance threshold of *5x10^-8^*. Specifically, associated DHSs were assigned into four bins:

- GW, if their tag SNP reached genome-wide significance of *5x10^-8^*,
- Locus GW (lGW), if another GWAS SNP within 0.1cM or 100kb of the DHS center reached genome-wide significance,
- True, if a GWAS SNP within 0.1cM or 100kb of DHS center reached genome-wide significance as reported by all studies for that disease in the NHGRI GWAS catalog^4^, and
- Novel otherwise.

The GWAS catalog^4^ was downloaded on May 08 2016, from:

https://www.ebi.ac.uk/gwas/api/search/downloads/full. We lifted the genome coordinates in the GWAS catalog (GRCh38.p5) to hg19 using liftOver (https://genome.ucsc.edu/cgi-bin/hgLiftOver). For each of the six autoimmune diseases we generated smaller catalog tables by searching for rows matching the following DISEASE.TRAIT terms:

RA (*n*=275): “Rheumatoid arthritis” or “Rheumatoid arthritis (ACPA-negative)”.

CD (*n*=330): “Crohn’ disease”, “Inflammatory bowel disease”, or “Inflammatory bowel disease (early onset)”.

UC (*n*=266): “Ulcerative colitis”, “Ulcerative colitis or Crohn’ disease”, “Inflammatory bowel disease”, or “Inflammatory bowel disease (early onset)”.

T1D (*n*=93): “Type 1 diabetes”.

MS (*n*=165): “Multiple sclerosis”.

SLE (*n*=118): “Systemic lupus erythematosus”.

The numbers in parentheses indicate how many rows (associations) in the catalog matched to each disease. We then mapped all reported lead SNPs to centimorgans as we described before. These smaller catalogs were used to determine which DHSs were defined as ‘True’.

Above we described how DHSs were classified. For a given disease, an associated locus (comprised of one or more associated DHSs) was said to belong to one of these categories based on the highest precedence DHS for that disease in that locus.

The second grouping was based on cross-replication of a DHS locus across two or more of the six autoimmune diseases we analyzed. Specifically, this grouping was determined by our grouping of DHSs into loci, using a genetic distance threshold of 0.01, and considering all GWAS together. An associated locus was then said to have cross-replicated if the DHSs comprising that locus, came from two or more diseases. Note that this would mean that the reported replication rates are all underestimates, as in practice, some of the GWAS we had were rather old, and either did not use imputation, or imputed but not to the same density or quality as more recent GWAS (e.g. due to lower density of genotyping array), leaving some DHSs untagged, and therefore decreasing possible replication rates.

The third grouping was testing if the association was previously reported in the GWAS catalog-i.e. IN vs. OUT of the catalog. A DHS was said to be IN the catalog if the SNP tagging that DHS was within 0.1cM or 100kb from a catalog lead SNP of genome-wide significance for that same autoimmune disease. Otherwise the DHS was said to be out of the catalog. Interestingly, out of the 649 “GW” DHSs we identified, 89 were not in the catalog (although having genome-wide significance in the original studies). With this respect, we noted that only one or two GWAS SNPs supported some GW loci, i.e. they did not have a nice peak structures as may be expected by the LD structure in a locus. We scanned each locus for the number of SNPs with *P* value < 10^-3^ in a region of 0.01cM around each associated DHS. A DHS was then flagged as poor quality if it had fewer than four SNPs with *p<1x10^-3^* supporting its association. In this work we used poor loci only as evidence for replication of non-poor loci, but not otherwise considered.

### KEGG enrichment analysis of associated genes

We assigned genes to DHSs by the rule of nearest TSS. We considered all associated DHSs discovered with an FDR of 0.005 for any of the six autoimmune diseases, which were not flagged by us as poor, for a total of 528 unique genes. We used WEB-GESTALT^94^ to identify enriched KEGG terms, retaining significant pathways with six or more genes, and a multiple-testing adjusted *P*-value smaller than 0.01 (as provided by the software). For a reference gene lists we used genes (nearest TSS) from the DHSs serving as input for csRWAS, i.e. genes most proximal to the adaptive-immune-specific DHSs. This was done to evaluate the contribution of the genetic variation associated with autoimmune diseases to pathways enriched over what was to be expected for adaptive-immune-specific DHSs.

### Confirming the presence of a stretch enhancer in a given cell type and gene

To confirm if a gene was under the control of a stretch enhancer (also known as a super enhancer), as was done for the *Blk* and *Irf5* examples given in the main text, we queried the dbSUPER database^95^, which summarizes stretch enhancer status from several studies across cell types, and links to the UCSC genome browser to allow an examination of the raw ChIP-seq signal in that region. We used the UCSC genome browser to make sure that the stretch enhancer regions were overlapping the cell-type-specific DHS stretches we identified.

### Characterizing transcription factor binding sites within DHSs

Transcription factor binding sites (TFBSs) are relatively short and degenerate in nature, making finding them in the genome many times possible by chance alone, without them carrying any function. Since DHSs mark the presence of some DNA biding protein, finding TFBSs in a DHS region increases the likelihood of that TFBS being functional. However, we already described how TFBSs within DHSs were enriched in a profile dependent manner. For example, Th17 specific DHSs were uniquely enriched for a Rorc TFBS. This makes finding a Rorc TFBS in say two DHSs, one specific to Th17 and the other for B cells, more likely to be functional in the first DHS than in the second. We note these more likely TFBSs as <u>congruent</u> TFBSs. When we discuss the letter grades, these TFBSs would be prefixed by ‘+’. We also detected the subset of TFBSs where a 1000 Genomes Project^96^ European SNP was intersecting their predicted binding sites. We next describe the details of the process of identifying TFBSs in DHSs, and how the subset of polymorphic TFBSs was deduced.

From the table in Figure 3a we de novo identified TFBSs enriched in one of the nine adaptive-immune-specific accessibility profiles. For these TFBSs we obtained cognate binding motifs as position weight matrices (PWMs)^97^ from Jaspar-2014^78^ and UniPROBE update 2015^79^. PWMs were available for the following TFBSs: SPI1, CEBPB, TAL1::GATA1, EBF1, TCFE2A_primary, RREB1, KLF4, ETS1, RUNX1, CREB1, TCF7L2, BATF::JUN, STAT5a::STAT5b, IRF1, GATA6_primary (same DNA binding domain as for GATA3), STAT1, and RORA_2 (same DNA binding domain as for RORC) (as identified by TOMTOM described earlier). Three enriched sequence motifs did not have a good match to a known TFBS and were named by our convention as [profile_name]_[integer], namely: HematoSC_1, Naïve_1, and Th1_1. For these motifs we used the output PWMs from MEME-CHIP. Taking PWMs as input we scanned all DHSs for TFBS occurrences using FIMO^98^. For each DHS a 300 bps region, centered on the DHS midpoint, was scanned. An initial scan was performed on the hg19 reference genome as retrieved using the R bioconductor^99^ library BSgenome.Hsapiens.UCSC.hg19. A secondary scan was performed on this reference sequence, after we mutated it by inserting alternate alleles at each SNP position. We refer to all variants throughout the text as SNPs although some belonged to small insertions and deletions (indels). The second scan was used to identify TFBSs created or improved when the alternate alleles were present, and to estimate the impact of SNPs on the TFBS binding affinity. We applied the following rules when mutating the reference sequences with the alternative alleles: 1. We mutated the reference sequence of a DHS beginning with the leftmost SNP and moving SNP-by-SNP to the rightmost SNP. For example, if a deletion took place, and then a SNP was found within this already deleted reference sequence, that SNP was not used as the region harboring the SNP was already removed. 2. In cases where more than one alternative allele was reported we used the alternative allele with the highest reported European MAF. All SNPs with a MAF>0 in the CEU (Europeans) samples of the 1000 Genomes phase 3 data were considered. In total 900,147 DHS-intersecting SNPs were identified, an average of 2.58 SNPs per DHS. Out of these, 156,126 (17.3%) were quite common (0.15<MAF), 82,261 (9%) were not as common (0.05<MAF<=0.15), and the majority of 661,760 (73.5%) were quite rare (MAF<=0.05) in the CEU samples. We also considered the MAF of these same SNPs in Asians (as a subset of the GWAS we used came from Asian/European mixed samples, as in the transethnic RA meta-analysis study). For EAS (east Asians) 1000 Genome samples, 147,345 (16.4%) were quite common (0.15<MAF), 66,148 (7.3%) were not as common (0.05<MAF<=0.15), and 686,654 (76.3%) were quite rare (MAF<=0.05). We kept records of every TFBSs found in the reference or alternate DHS sequences, with a P-value smaller than 0.01. The command line used for FIMO was: fimo --bgfile motif-file --max-strand --text --thresh 0.01 $PWM_FILE $DHS_SEQUENCE_FILE > $FIMO_OUTPUT_FILE.

### Assigning letter grades for polymorphic transcription factor binding sites

A small proportion of predicted TFBSs in either the reference or the alternate DHS sequences had an intersecting SNP. We focused on these polymorphic TFBSs (pTFBSs) as they provide a resource to design follow up studies to determine causality of genetic associations. Since we identified the accessibility profiles as a key factor in other aspects of this work (e.g. enriching for genetic associations or providing a regulatory context), we also annotated the subset of congruent TFBSs (as defined for characterizing TFBSs). Lastly, we also considered SNP MAF as a relevant factor as it could be relevant in deciding which SNP to follow up on. All of the above factors per pTFBSs were integrated into a simple three-letter grade string, making them human accessible as described next.

We first identified all cases where a TFBS was found with a SNP intersecting it, denoting these as pTFBSs. For a given DHS, multiple pTFBSs can be found for the same TF. In such cases we retained the pTFBS with the highest FIMO score for further analyses (i.e., the highest score of the best match for each TF, when considering both the reference and alternative sequences for each DHS). We annotated these pTFBSs with a three-letter code. The first letter captures the goodness-of-fit between the cognate motif and the identified TFBS, the second letter captures the MAF of the intersecting SNP in CEU samples (Europeans), and the third letter captures the MAF in EAS samples (Asians).

The letter grade alphabet was (“+”, “A”, “B”, “C”). The first letter reported the TFBS score (in the following order of precedence), with very good matches to the consensus-binding site receiving an A grade (*P* value < 10^-4^ in either primary or secondary scan), medium matches a B (10^-3^ <P value <= 10^-4^ in either primary or secondary scan), and more degenerate matches a C (10^-2^ <P value <= 10^-3^ in either primary or secondary scan). The second letter indicated the MAF of the intersecting SNP in Europeans, with A denoting common (0.15<MAF), B not as common (0.05<MAF<=0.15) and C rare SNPs (MAF<=0.05). Similarly, the third letter indicated the MAF in Asians. For congruent TFBSs we added a plus sign prefix. A couple of examples should make these three-letter grades clear. Assume the examined DHS matched a B-cell specific accessibility profile and that two pTFBSs were found in this DHS, SPI1 with *p<10^-4^* (enriched in B cell specific DHSs), and RORC with *10^-2^<p<10^-3^* (not enriched in B-cell specific DHSs). The SPI1 match was intersected by a common SNP in both CEU and EAS, and the RORC match was intersected by a rare SNP in both CEU and EAS. The SPI1 TFBS would therefore be annotated as +AAA, indicating the TFBS was found in a context it was enriched in (+), the TFBS was a good match to the cognate sequence motif of SPI1, and that the SNP intersecting it was common in both populations. The RORC TFBS would therefore be annotated as CCC, indicating the TFBS was degenerate (a bad match to the cognate sequence motif of RORC) and that the SNP intersecting it was rare in both populations. Reading the letter grades saves the need for many back-references to the primary data (i.e. SNP MAFs, TFBS P-values, and DHS accessibility profiles), helping to decide in this example that the SPI1 pTFBS looks like a more promising TFBS to follow up on.

### A cross disease hierarchical network model

We constructed a hierarchical network model with five levels. This network was visualized in the open source cross-platform network visualization tool, Cytoscape^43^, and made available for dynamic exploration using the same tool (supplementary file 1). The trait nodes (autoimmune diseases) were placed at level one, associated loci (loci nodes) at level two, DHS tagging SNPs (SNP nodes) at level three, associated DHSs (DHS nodes) at level four, and DHS most proximal TSS genes (gene nodes) at level five. Edges were directed from nodes at a lower level to the nodes at the level directly above it. Connected to DHS nodes were two types of DHS information nodes: one for the nine adaptive-immune-specific accessibility profiles, and another for the nineteen TFBSs enriched in adaptive-immune-specific DHSs. Accessibility profile nodes were connected to the DHSs that matched them (profile to DHS edge direction), e.g. the B-cell accessibility profile node connected to B cell specific DHSs. Similarly, TFBS nodes connected to all DHSs harboring congruent polymorphic TFBSs for that TF, e.g. the ETS1 node connected to all DHSs harboring a congruent (‘+’ grade), polymorphic ETS1 motif. Edges connecting TFBSs to DHSs were given as a name their three-letter grades.

To aid interpretations, we colored nodes for genes, TFBSs, and accessibility profiles by their most likely accessibility profile, which was calculated as follows. For TFBSs the color was determined as the color matching the accessibility profile where the motif was most enriched in (in terms of percentages, as per Fig. 3a). For example, RUNX1 received a color matching the Th2 accessibility color. For genes, we counted the accessibility profiles from all associated DHSs that had that gene as their most proximal TSS. The profile corresponding to the highest count was then selected for that gene. In cases of a tie, we took the best match accessibility profile per DHS (a DHS may be matched to more than one profile), and recalculated the counts. If still a tie, we used the accessibility profile with the lowest profile-match-rank for the tied profiles.

The topology of the network and directionality of the edges were designed to aid in exploration of the network through descending along incoming edges, or ascending along outgoing edges. This was key for allowing exploration of the results, and for generating subnetworks per gene, loci, or any other starting node type. For example, to study the role of B cells in autoimmune diseases, one can select the B-cell accessibility profile node, then select all adjacent nodes (DHSs), then deselect the B-cell accessibility profile node, and continue up the network through outgoing edges to get the genes, and down the network through incoming edges to link to the SNPs, loci, and diseases. The B-cell accessibility profile node can then be reselected and a subnetwork generated. This is an example of how to query the network for a particular insight, but users are encouraged to come up with their own queries. Exploring the network assumes familiarity with Cytoscape for which many tutorials are available online.

### Generating regulatory Manhattan plots

Regulatory Manhattan plots (RMPs) were generated by the R statistical software^100^. The recombination rates were directly obtained from HapMap Phase II data^93^. To calculate the linkage disequilibrium quantity, r^2^, between an associated DHS tagging SNP and nearby SNPs, we used the phased genotypes of the CEU European individuals from the 1000 Genomes Phase I data^101^ and the value was calculated by the command --hap-r2 implemented in VCFtools^102^. We augmented the typical Manhattan plots using regulatory information as described in the main text and the figure legends. One RMP was generated per locus across all six autoimmune diseases to allow visual evaluation of cross-disease status. RMPs highlight SNPs tagging associated DHSs by plotting a small pie chart in their place. Each such pie chart has nine wedges, a wedge per adaptive-immune-specific accessibility profile. Purple wedges indicate the profile membership of DHSs tagged by a given SNP. To facilitate loci interpretation, in cases where multiple associated DHSs were found, we placed a larger, summary pie chart on the top right of the lead RMP. These larger pie charts count how many times each accessibility profile was assigned to any of the associated DHSs in a locus. Examining profile counts provides useful information about the prominent cell type context of an association. We also provided with each RMP, a subtable reporting further information about associated DHSs and the polymorphic TFBSs (pTFBS) they harbored. Specifically, these tables report which pTFBSs were present in a locus (per DHS), and show their letter grades to help decide on pTFBSs for follow up work. Entries with congruent pTFBSs (prefixed by a ‘+’) were further highlighted in purple. RMPs were ordered by disease from top to bottom by their most significant DHS *P* values. The center of RMPs was determined as the average of all associated DHS positions. That is, the center of the plot gravitates towards the region of highest associated DHSs density. All RMPs were 400kb long. Loci longer than 150kb were not plotted, but RMPs from the lower cM cutoff of 0.025cM would likely have broken such long loci into smaller chunks.

RMPs are available for download as zipped files (Supplementary files 2 and 3). We made the size of each RMP pdf file very small (~50 kb), so opening many RMPs at once should not be a problem on any modern computer.

### Testing why DHSs of genome-wide significance were not found in the GWAS catalog

Out of the 89 “GW” DHS associations that were not in the catalog although they reached genome-wide significance, 46 were of poor quality and were thus likely removed by the authors as part of some quality control step. Out of the remaining 43 DHS associations 23 were associated with SLE, where the original GWAS data^40^ was yet to be incorporated into the GWAS catalog as of the time of our download (May 08 2016). Out of the remaining 20 DHS associations, 11 were associated with RA in two independent loci that cross-replicated (Fig. S15). One locus surrounding *FAM231B* was reported in of the original GWAS^36^ (their supplementary table 1) but the second, surrounding *ZFP36L1* was not reported; perhaps because it did not replicate in their stage 2 and stage3 studies. It was unclear to us why the first locus was not reported in the catalog, suggesting that the catalog is still incomplete with respect to reported associations. Out of the remaining 9 DHS associations 4 were associated with CD, in two loci (Fig. S16). The first locus surrounding *TTC33* was reported in the original study as part of a larger locus upstream of the *PTGER4* gene^38^ (their table 1, reported in hg18 coordinates on chr5, we found this on chr16 in hg19 coordinates). This locus was reported in the catalog as well but did not meet our genetic or physical DHS distance criteria to be reported by us as cataloged. The second locus surrounding *BRD7* cross-replicated in our study with UC, but was not reported in the original study or in the catalog. Given that this locus reached genome-wide significance and replicated with UC, we suggest that this is a true genome-wide association. Out of the remaining 5 DHS associations, 3 were associated with MS in one independent locus surrounding *RMI2* that cross-replicated with T1D and CD (Fig. S17). The locus was reported in the original study^38^ (their supplementary table A), using the hg18 name for *RMI2 (C16orf75)*, but was not found in the catalog. The remaining 2 DHSs were associated with UC in two independent loci. The first locus in the *REPS1* gene region seems like an artifact showing the limitations of our suggested heuristic to flag poor associations (locus 831, 0.01cM, see RMP). The second locus in the *NR6A1* was also of somewhat questionable quality (locus 953, 0.01cM, see RMP). In support of these loci being of poor quality, both failed to cross-replicate here. Thus, these loci may not have met some quality control steps taken by the authors. We would note that although we did our best to understand why GW loci were not in the catalog, and sometimes not reported by the original authors of the study, we do not imply that our analysis relying solely on GWAS summary statistics, could replicate all quality control steps that may have been taken by the original authors. As such, the fact that some loci were not reported by the authors but seemed to be of high quality to us, does not preclude these loci still being poor by some other criterion that we did not or could not evaluate.

### Replication studies

The gold standard for validation of any genetic study is replication in an independent sample^63^. We evaluated the validity of associations found by GWAS as a function of discovery P-value threshold, and contrasted it with the same for RWAS (all DHSs) and csRWAS (DHSs specific to adaptive-immune-specific accessibility profiles). For this analysis, DHSs were associated to their nearest GWAS SNP to allow fair comparisons with GWAS (note that before DHSs were assigned the minimum P-value SNP in a 2.5kb region surrounding each DHS). For rheumatoid arthritis (RA) we had two independent studies, one of participants of Asian descent (n=22,515) and the other of participants of a European descent (*n*=58,284). For Crohn’s disease (CD) we had two non-independent studies, one of UK-descent participants (*n*=4,664), and the other of a meta-analysis of several European descent CD studies, which included the UK study (*n*=21,389). We removed the shared UK-participants from the CD meta-analysis as follows.

The CD meta-analysis^38^ in addition to reporting the meta *Z*-scores and meta *P*-values, also reported the per-study *Z*-scores for each SNP. We recalculated the meta *Z*-scores without the UK-participant using Stouffer’s method for combining *Z*-scores, thereby making the newly-calculated meta-analysis independent from the UK study. Specifically, the *Z*-scores for each SNP were combined as follows:, 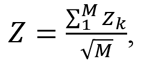 where *M* denotes the number of CD studies, and *Z_k_* the SNP *Z*-score in GWAS *k*.

Next, we used the smaller GWAS for each disease for discovery, at a range of discovery *P*-value thresholds, and estimated the replication success of discovered associations in the larger GWAS.

Formally, Let *S_1_* and *S_2_* denote the sets of SNP *P*-values in the discovery and replication GWAS, respectively. We first restricted the analysis to the intersection of these two sets, only considering SNPs that were measured in both studies. Denote the intersection of *S_1_* and *S_2_* as *S*. From here on we only consider SNPs in S. Let *S_d,p_* be the set of SNPs with a *P*-value smaller than a set threshold, *p*, in the discovery GWAS. We defined the subset of SNPs in *S_d,p_* that were below nominal significance of 0.01 in the replication study as replicating SNPs. Denote this subset of replicating SNPs by *S_r,p_*. Then the replication rate of SNPs at a given discovery *P* value threshold was simply:

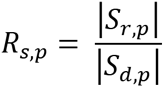

where in *R, s* denotes the study (CD or RA) and *p* the discovery *P*-value threshold. Vertical bars denote the cardinality of the set (i.e. the number of SNPs in the set). That is, the replication rate was the ratio of discovered and replicating SNPs, to all discovered SNPs.

SNP replication rate however is not a good indicator for the replication rate of independent associations, as SNPs within a DNA region are correlated by linkage disequilibrium. The association units in GWAS are typically defined as independent loci, i.e. a set of genomic regions that are not in LD with one another. We defined associated GWAS loci using the pseudo code we described earlier for DHS associations with a genetic distance cutoff *d*=0.1 cM.

We now had a grouping of discovered SNPs, *S_d,p_*, to a set of independent loci, *L_d,p_*. The subset of replicating loci, *L_r,p_*, was then determined as those loci where one or more of the associated SNPs comprising them replicated with nominal significance of 0.01. Using *L_d,p_* and *L_r,p_* the loci replication rate per SNP, per discovery P value, was calculated as:

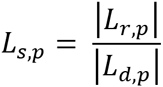

where in *L*, *s* denotes the study (CD or RA) and *p* the discovery *P*-value threshold.

For RWAS and csRWAS, the number of discovered and replicated SNPs was calculated as follows. For a given *P*-value threshold we first determined which DHSs where associated in the discovery study. As each DHS was assigned a single SNP, we could map DHSs to SNPs, and then to loci as determined for GWAS. The number of RWAS or csRWAS discovered loci, was then simply the subset of GWAS loci that mapped to at least one associated DHS. That is, for a given significance threshold, GWAS defined the total possible number of discovered loci, and the total number of replicating loci. A DHS locus was then considered as replicating if one of the discovered DHS SNPs within it replicated. It is important to understand that although GWAS defines the set of all discovered loci, and the subset of discovered and replicating loci, csRWAS and RWAS identified a subset of these loci, and therefore may have very different replication rates than GWAS, or one another.

We calculated the reciprocal discovery *P* value required to attain a given replication rate, by calculating the replication rates at finely increasing discovery *P* value thresholds, and keeping the *P* values thresholds when a particular replication rate was attained (replication rates were calculated using the above procedure).

### Code availability

We aimed to describe our approach in detail to allow replication. In line with this we provided command line calls with arguments when using common tools (e.g. MEME-CHIP), parameters when using online tools (e.g. WEB-GESTALT), pseudo code when a simple procedure can be described concisely (loci assignments), and functions with their parameters when using statistical packages (e.g. GenABEL). For all other types of analysis code is available upon request.

## Supplementary information

Tables S1-S5 with readme files per table (tables are in a tab delimited format), the Cytoscape network with a readme file, and RMPs for all loci with readme file. Links to download the files are provided below. Click on a link, or copy and paste it to a web browser. A Google drive interface will open up showing all the zipped files (e.g. a table and a readme file), find the download icon (a down facing arrow on top of the page), click to download and unzip the files.

### Supplementary tables

Supplementary Table 1 (101 MB):

Accessibility annotation, CPKM matrix, and match-scores per accessibility profile https://drive.google.com/open?id=0B0i1cwUJRRHwcEdXbGJ6YWZaNHM

Supplementary Table 2 (76 MB):

DHSs to GWAS proxy SNPs (both to nearest SNP, and to minimum P-value SNP in a 2.5kb region around DHS midpoint) https://drive.google.com/open?id=0B0i1cwUJRRHwT3hmc0ZNSGtCbHM

Supplementary Table 3 (122 KB):

Significant associations (per DHS) https://drive.google.com/open?id=0B0i1cwUJRRHwV2tSeFRMX2NXdVU

Supplementary Table 4 (23 KB):

Significant associations (per locus) https://drive.google.com/open?id=0B0i1cwUJRRHwWXU3XzZhR1J4MWc

Supplementary Table 5 (93MB):

DHSs to scored, polymorphic TF binding sites https://drive.google.com/open?id=0B0i1cwUJRRHwNm9SVjBvcGoza3c

### Cytoscape network file

Supplementary file 1, cM cutoff of 0.01, FDR<0.005 (458 KB): https://drive.google.com/open?id=0B0i1cwUJRRHwTm01VmdXN20wWjQ

### Regulatory Manhattan plots (RMPs)

Supplementary file 2, cM cutoff of 0.01, FDR<0.005 (72 MB):
https://drive.google.com/open?id=0B0i1cwUJRRHwQU0yVGlzYzlzUkE

Supplementary file 3, cM cutoff of 0.0025 (only used to break down loci that were too large, >150kb, under the 0.01cM cutoff), FDR<0.005 (65 MB):
https://drive.google.com/open?id=0B0i1cwUJRRHwYnpOLWw2U1NnV2M

## Figures

All figures appear below but are also available as two downloadable .pdf files from the following private links.

Main figures, private links:

https://drive.google.com/open?id=OB_nf7cPOLTBSSVpuajRxQlZySVk

Supplemental figures, private link:

https://drive.google.com/open?id=OB_nf7cPOLTBSQUtfZOpUV3hQLW8

